# Structural requirements for dihydrobenzoxazepinone anthelmintics: actions against medically important and model parasites - *Trichuris muris*, *Brugia malayi*, *Heligmosomoides polygyrus* and *Schistosoma mansoni*

**DOI:** 10.1101/2020.11.17.384933

**Authors:** Frederick A Partridge, Carole JR Bataille, Ruth Forman, Amy E Marriott, Josephine Forde-Thomas, Cécile Häberli, Ria L Dinsdale, James DB O’Sullivan, Nicky J Willis, Graham M Wynne, Helen Whiteland, John Archer, Andrew Steven, Jennifer Keiser, Joseph D Turner, Karl F Hoffmann, Mark J Taylor, Kathryn J Else, Angela J Russell, David B Sattelle

**Author notes:** Corresponding authors: (DBS), (KJE), (AJR). These authors contributed equally (Frederick A Partridge, Carole JR Bataille, Ruth Forman, Amy E Marriott, Josephine Forde-Thomas, Cécile Häberli).

## Abstract

Nine hundred million people are infected with the soil-transmitted helminths *Ascaris lumbricoides* (roundworm), hookworm, and *Trichuris trichiura* (whipworm). However, low single-dose cure rates of the benzimidazole drugs, the mainstay of preventative chemotherapy for whipworm, together with parasite drug resistance, mean that current approaches may not be able to eliminate morbidity from Trichuriasis. We are seeking to develop new anthelmintic drugs specifically with activity against whipworm as a priority, and previously identified a hit series of dihydrobenzoxazepinone (DHB) compounds that block motility of *ex vivo Trichuris muris.* Here we report a systematic investigation of the structure-activity relationship of the anthelmintic activity of DHB compounds. We synthesised 47 analogues, which allowed us to define features of the molecules essential for anthelmintic action, as well as broadening the chemotype by identification of dihydrobenzoquinolinones (DBQ) with anthelmintic activity. We investigated the activity of these compounds against other parasitic nematodes, identifying DHB compounds with activity against *Brugia malayi* and *Heligmosomoides polygyrus*. We also demonstrated activity of DHB compounds against the trematode *Schistosoma mansoni,* a parasite that causes schistosomiasis. These results demonstrate the potential of DHB and DBQ compounds for further development as broad-spectrum anthelmintics.

**Author summary:** Around a billion people are infected by the soil transmitted helminths *Ascaris,* hookworm and whipworm. In the case of whipworm, the benzimidazole drugs, which are distributed to school children in affected areas, have low cure rates. This means that finding an improved treatment for whipworm is a priority. We previously identified five DHB compounds in a screen for new compounds active against whipworm. Here we systematically dissect these molecules, making 47 modified versions of the compounds. This allowed us to define the features of these compounds that are important for activity against whipworm. We also demonstrate activity of DHB compounds against other parasitic nematodes, and against *Schistosoma mansoni,* a trematode parasite. These results show the potential for further development of DHB compounds as broad-spectrum anthelmintics.

## Introduction

900 million people are infected with soil-transmitted helminths causing a global burden of around two million disability-adjusted life years [1,2]. Because of this, the World Health Organization has set a goal to achieve and maintain elimination of soil-transmitted helminth morbidity by 2030 [3]. Huge mass drug administration efforts are underway, distributing hundreds of millions of doses of benzimidazole drugs (albendazole and mebendazole) to school-age children in affected areas annually or biannually as preventative chemotherapy (PCT).

Benzimidazole drugs are partially effective against whipworm *(Trichuris trichiura)* when administered as a course of treatment, reaching cure rates of around 43% [4]. However, for mass drug administration, practicalities and scale mean that only one dose is given. In contrast to *Ascaris lumbricoides,* where a single dose of benzimidazole drugs cures around 90-95% of infected individuals, the single dose cure rate for whipworm is low, around 30% [5,6]. The current mass drug administration protocol may therefore not be able to break transmission and reduce the prevalence of moderate to heavy whipworm infections to below 2% as required to eliminate morbidity [3]. Due to the poor single dose efficacy of the benzimidazole drugs against whipworm, there have been extensive efforts to identify more efficacious drug combinations [6]. Of these the most promising to date is a combination of albendazole plus the N-type nicotinic acetylcholine receptor agonist oxantel pamoate, which has a single dose cure rate reported to be between 31 and 83% [7–10]. A second drug, moxidectin, also shows promise to be added to albendazole for improved control of whipworm [7].

Of concern, however, is the possibility that drug resistance may become prevalent, derailing the push towards control of whipworm. Currently there is only indirect evidence of this possibility. In a meta-analysis, it has been shown that egg reduction rates and cure rates after albendazole treatment are decreasing over time [11]. Polymorphisms in the beta-tubulin gene that are associated with benzimidazole resistance are found in populations of human whipworm, and the frequency of these polymorphisms increased after albendazole treatment [12,13]. The Starworms project will establish a valuable system to monitor benzimidazole drug efficacy and the potential emergence of anthelmintic resistance due to soil-transmitted helminth control programs [14]. Because of these two problems – low efficacy of existing drugs against whipworm, and concerns about development of resistance to these drugs – we and others have been pursuing a strategy of identifying new anti-whipworm compounds, via a mixture of repurposing and *de novo* small molecule screening [15–22].

Beyond soil transmitted helminths (STH), lymphatic filariasis and schistosomiasis are two medically important tissue helminthiases prioritised for global or regional elimination via mass PCT, as outlined in the WHO Roadmap 2030 implementation targets [23]. A related filarial nematode, *Onchocerca volvulus*, is also targeted for regional elimination. Reliance on a few, or in the case of onchocerciasis and schistosomiasis, a single chemotherapeutic agent (ivermectin and praziquantel, respectively) used ‘en masse’ for PCT, is a vulnerability of current elimination strategies, considering the potential for development of drug resistance. As with STH, annual or semi-annual mass drug administrations extending upward of 20 years are required to break transmission with current drugs due to incomplete adulticidal / selective larvicidal activity profiles of the implemented anti-filarial or schistosomicidal agents. Alternative strategies, for instance, development of a short-course curative treatment for filariasis, would be a step-change to reduce elimination time frames [24,25].

We previously described a hit series of five dihydrobenz[*e*][1,4]oxazepin-2(3*H)*-one (DHB) compounds with anthelmintic activity against *ex vivo T*. *muris* [18]. Here we report our progress in expanding this hit series and understanding the relationship between structure and anthelmintic activity. We also extend our investigations of the activity of the DHB compounds against *Brugia malayi,* a causative agent of lymphatic filariasis, *Heligmosomoides polygyrus bakeri*, a mouse gastrointestinal nematode model, and the human blood fluke, *Schistosoma mansoni*.

## Materials and Methods

### Ethics statement

All experimental procedures involving *T. muris* were approved by the University of Manchester Animal Welfare and Ethical Review Board and performed within the guidelines of the Animals (Scientific Procedures) Act, 1986.

All experiments involving *Brugia malayi* were approved by the ethical committees of the University of Liverpool and Liverpool School of Tropical Medicine (LSTM) and conducted under Home Office Animals (Scientific Procedures) Act 1986 (UK) requirements and the ARRIVE guidelines.

The work on *Heligmosomoides polygyrus* was approved by the local veterinary agency, based on Swiss cantonal and national regulations (permission no. 2070).

For experiments involving *Schistosoma mansoni,* all procedures performed on mice adhered to the United Kingdom Home Office Animals (Scientific Procedures) Act of 1986 (project licenses PPL 40/3700 and P3B8C46FD) as well as the European Union Animals Directive 2010/63/EU and were approved by Aberystwyth University’s (AU) Animal Welfare and Ethical Review Body (AWERB).

### Chemical synthesis

Compounds were synthesised from commercially available starting materials, and fully characterised by Nuclear Magnetic Resonance Spectroscopy and Mass Spectrometry. Full experimental details and analytical data are provided in the Supporting Information.

### Isolation of *T. muris* adults

Male and female severe combined immunodeficient (SCID) mice were bred in house at the University of Manchester and used at age 8-12 weeks. Mice were maintained at a temperature of 20-22°C in a 12h light, 12h dark lighting schedule, in sterile, individually ventilated cages with food and water *ad lib.*

The parasite was maintained and the infectivity of the administered *T. muris* eggs was assessed as previously described [26,27]. For generation of adult *T. muris* worms 150 infective eggs were given per oral gavage in water to each SCID mouse. 35 days post infection mice were sacrificed via schedule one methods. At necropsy the caecae and colons were removed, opened longitudinally and washed with pre-warmed RPMI-1640 media supplemented with penicillin (500U/ml) and streptomycin (500μg/ml). Adult *T. muris* worms were gently removed using fine forceps under a dissecting microscope and maintained at 37°C in RPMI-1640 media supplemented with penicillin (500U/ml) and streptomycin (500μg/ml).

### *T. muris* adult motility assay

Single adult worms were placed in microplate wells containing 100 μL of RPMI-1640 medium, penicillin (500 U/mL), streptomycin (500 μg/mL) and 1 μl (1% v/v) dimethylsulfoxide (DMSO) or compound dissolved in DMSO. Assay plates were incubated at 37°C with 5% CO_2_. The INVAPP system was used to quantify worm motility [28,29]. Movies of the whole plate were recorded (20 frames, 100 ms interval) and motility determined by thresholding the variance of each pixel in the image over time [30]. Compounds were initially tested at 100 μM. Those showing activity were also tested at lower concentrations, typically 50 and 75 μM, and EC_50_ estimates were measured for compounds of interest using the a log-logistic model and the R package *drc* [31].

### *B. malayi* parasite production

The life cycle of *B. malayi* was maintained in *Aedes aegypti* mosquitoes (Liverpool strain) and inbred Mongolian gerbils housed at the Biomedical Services Unit, University of Liverpool under specific pathogen-free conditions. Microfilariae were harvested from experimentally infected Mongolian gerbils via catheterization under anaesthesia and fed to mosquitoes in human blood at 20,000 mf / ml using artificial membranes heated to 37 °C. Mosquitoes were reared for 14 days with daily sugar-water feeding to allow development to larval stage (*Bm*L3). At day 14, *Bm*L3 were collected from infected mosquitoes by stunning at 4 °C, crushing and concentrating using a Baermann’s apparatus and RPMI-1640 media. Male IL-4Rα^−/−^IL-5^−/−^ BALB/c mice (gifted by Prof. Achim Hoerauf, University of Bonn, Germany) aged 6-8 weeks, weighing 18-24 g were infected intraperitoneally with 150 *Bm*L3 and left for 12 weeks to develop to patent adult stage as previously detailed [32].

### *B. malayi* microfilaria assay

*Brugia malayi* microfilariae (mf) were harvested from Mongolian gerbils via intraperitoneal lavage and purified using PD-10 columns (Amersham). Mf densities were then adjusted to 8,000 / well in complete medium consisting of RPMI-1640 supplemented with 1% penicillin-streptomycin, 1% amphotericin B and 10% FBS within 96 well plates.

33 test compounds (10 mM stock in 100% DMSO) were initially tested against mf. Compounds were diluted to 10 μM in complete medium and added to the plated mf. Three replicates were used for each compound, and each plate included ivermectin (50 μM) as a positive control and DMSO (0.5% v/v) as a negative control. Assay plates were incubated for 6 days at 37°C, 5% CO_2_. Mf were scored daily for motility as a proxy of nematode health, using a 5 point scoring system (4 = fully motile, 0 = no motility) as described previously [33]. Compounds found to reduce motility were progressed to a secondary screen, whereby the MTT assay was employed at day 6 to assess parasite viability quantitatively. For this, excess media was removed from wells and mf were incubated with 0.5 mg/ml MTT (3-(4,5-Dimethylthiazol-2-yl)-2,5-Diphenyltetrazolium Bromide (Merck) in PBS at 37°C for 90 min. After washing in PBS and centrifugation, mf pellets were incubated in 100% DMSO for 1 h at 37°C to solubilize the blue formazan product. Samples were read at OD 490 nm on a 96-well plate reader (Varioskan, Bio-Rad). Compounds exhibiting the greatest activity on parasite viability were progressed further for drug dose titration assays.

### *B. malayi* adult assay

Adult female *B. malayi* of 12-24 weeks of age were isolated from susceptible IL-4Rα^−/−^IL-5^−/−^ immunodeficient mice, washed in PBS and added to lymphatic endothelial cell co-cultures (HMVECdly; LEC; Lonza) at a density of two parasites per well. Successful test compounds from the mf assay were diluted to 10 μM and added to the trans-wells in 6 ml endothelial basal media with supplements (EGM-2 MV; Lonza). Twelve replicates (n = 6 wells) were set up per group, with flubendazole (10 μM; Sigma) and DMSO (0.5% v/v) added as controls. Plates were incubated for 14 days at 37 °C, 5% CO_2_ with daily motility scoring, as above. Individual parasites were taken for MTT analysis at day 14.

### Heligmosomoides polygyrus

*H. polygyrus* larvae (L3) were obtained by filtering the faeces of infected mice and cultivating the eggs on an agar plate for 8-10 days in the dark at 24°C. 30-40 L3 were placed in each well of a 96-well plate for each compound in the presence of 100 μl RPMI 1640 (Gibco, Waltham MA, USA) culture medium supplemented with 5% amphotericin B (250 μg/ml, Sigma-Aldrich, Buchs, Switzerland) and 1% penicillin 10,000 U/ml, and streptomycin 10 mg/ml solution (Sigma-Aldrich, Buchs, Switzerland) with the test drugs (100 μM concentration). Worms were kept at room temperature for 72 h and for evaluation 50-80 μl of hot water (≈80°C) was added to each well and the larvae that responded to this stimulus (the moving worms) were counted. The proportion of larval death was determined. Compounds were tested in duplicate at 100 μM. Control wells were included in each experiment, which included the highest amount of solvent (1% DMSO).

### *S. mansoni* Roboworm assay

*Biomphalaria glabrata* (NMRI and the previously described pigmented strains [34]) infected previously with *S. mansoni* (Puerto Rican strain) miracidia were exposed for 1.5 hrs under light at 26 °C. Cercariae were collected and mechanically transformed into schistosomula as previously described [35]. Mechanically-transformed schistosomula were subsequently prepared for high throughput screening (HTS) on the Roboworm platform according to Crusco *et. al.* [36]. All compounds were tested in duplicate during dose response titrations (50 μM, 40 μM, 30 μM, 20 μM, 10 μM in 0.625% DMSO). Assay controls included 10 μM (in 0.625% DMSO) auranofin (positive control; Sigma-Aldrich, UK) and 0.625% DMSO (negative control). Schistosomula phenotype and motility were quantified after 72 hr co-culture with compounds as previously described [37]. Compounds passing both phenotype (−0.15) and motility (−0.35) thresholds were classified as hits. Z’ scores for all assays were above 0.35 [38].

## Results

### Novel DHB Chemistry

We have recently reported the identification of five dihydrobenzoxazepinone (DHB) hit compounds as a new family of molecules active against *T. muris* adult motility [18]. Further, one of the compounds **OX02983** was also found to be efficacious at reducing the ability of eggs to establish infection *in vivo*. As we identified a limited number of active DHB family members in the first instance via a library screen, we aimed to investigate the DHB chemotype systematically with the goal of understanding their structure-activity relationships (SARs) and improving potency. **OX02983** was used as a starting point of our investigation.

The synthesis used to prepare **OX02983** was adapted to systematically alter all the different cycles **A-D**, as shown in Fig 1. The first step was a reductive amination of the requisite aminobromobenzoate with the desired aldehyde to install cycles **A** and **C**. This was followed by a ring-closure step to generate cycle **B** and finally a cross-coupling reaction to add cycle **D** (Scheme 1).

**Fig 1.**
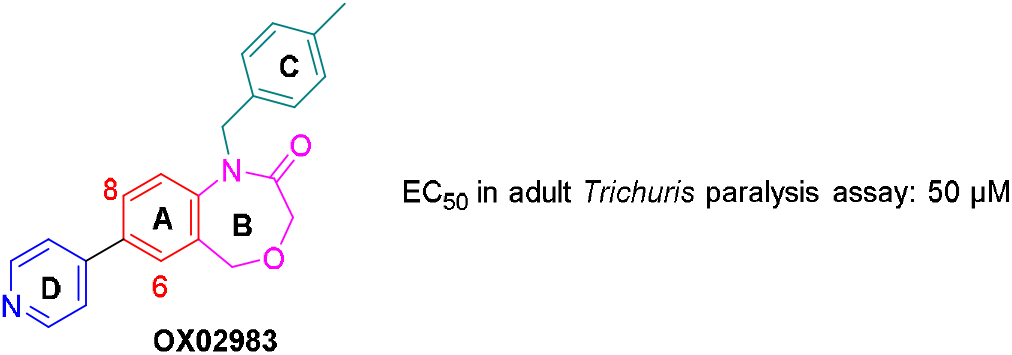
Structure of OX02983 highlighting the four cycles labelled A-D.

**Scheme 1.**
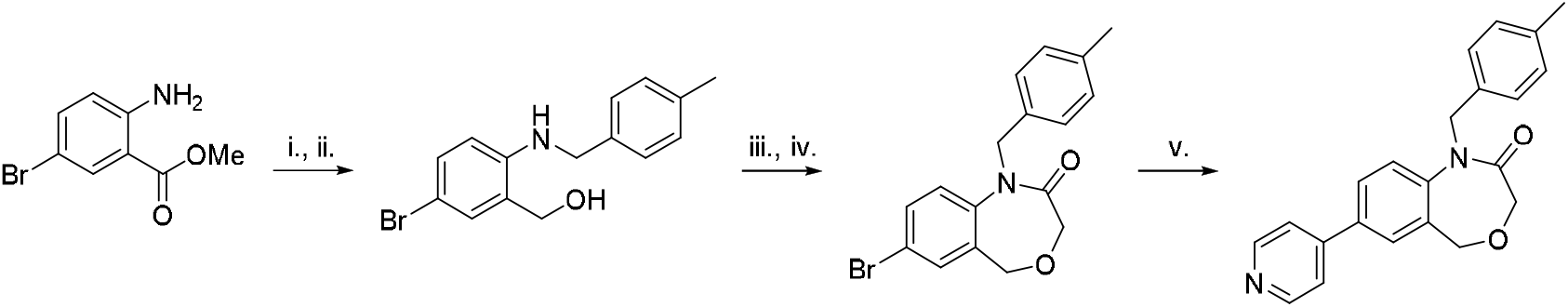
Representative scheme for synthesis of DHB compounds. i. 4-methylbenzaldehyde (1.5 eq.), AcOH (0.5 eq.), NaBH(OAc)_3_, CH_2_Cl_2_, 0 °C to rt, 48 h; ii. LiAlH_4_ (1 M in THF, 3.5 eq.); iii. chloroacetyl chloride (2.0 eq.), NEt_3_ (2.0 eq.), THF, 0 °C to rt, 16 h; iv. NaOH (10 N, aq.), rt, 2 h; v. 4-pyridyl-B(OH)_2_ (1.1 eq.), Pd(dppf)Cl_2_ (5 mol%), K_2_CO_3_ (3.0 eq.), 1,4-dioxane/H_2_O (4:1), 90 °C, 18 h.

It was decided to conduct a systematic structure activity relationship (SAR) investigation and alter the four different cycles within the structure of **OX02983** to understand their importance in the activity against *T. muris* with a view to improving efficacy. As the synthesis is linear, it was logical to investigate from **A** to **D**. We therefore started with core **B**, to ascertain the importance of regiochemistry and relative orientation of the substituents (Table 1). All the prepared compounds were screened using an automated adult *T. muris* motility assay [28] at 100 μM. Active compounds were also tested at lower concentrations and/or an EC_50_ value determined to assess their relative activity.

**Table 1.**
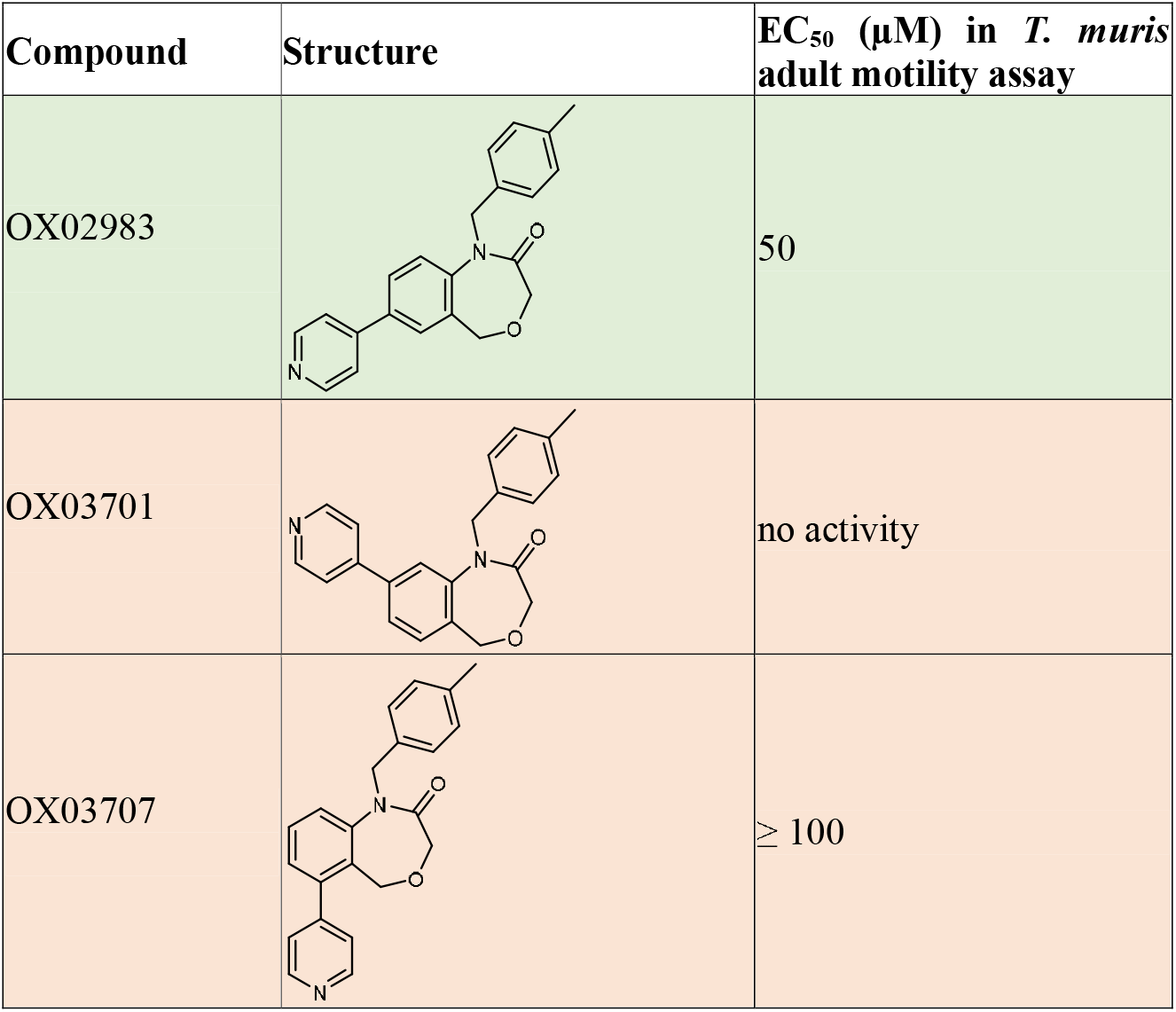
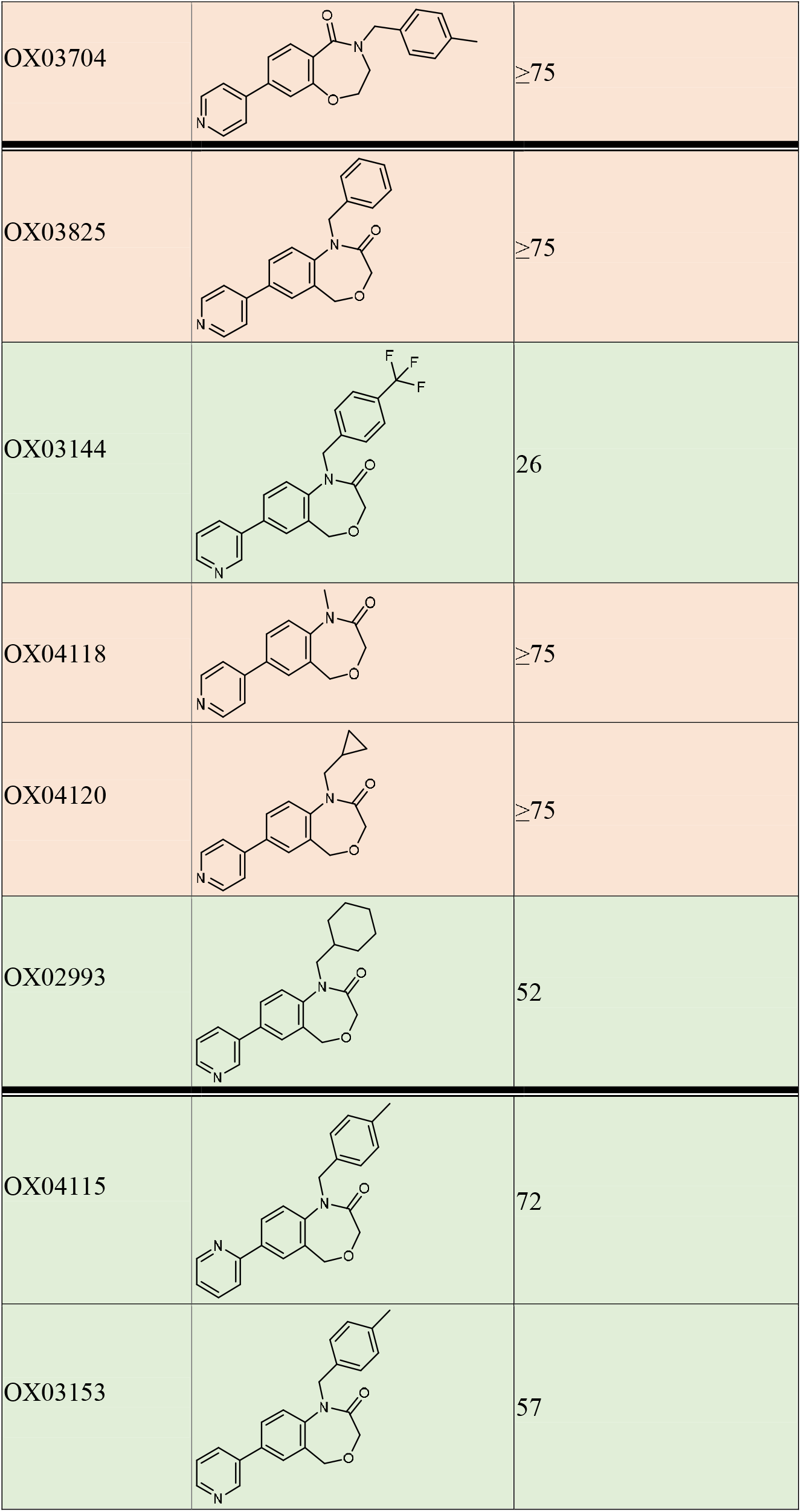
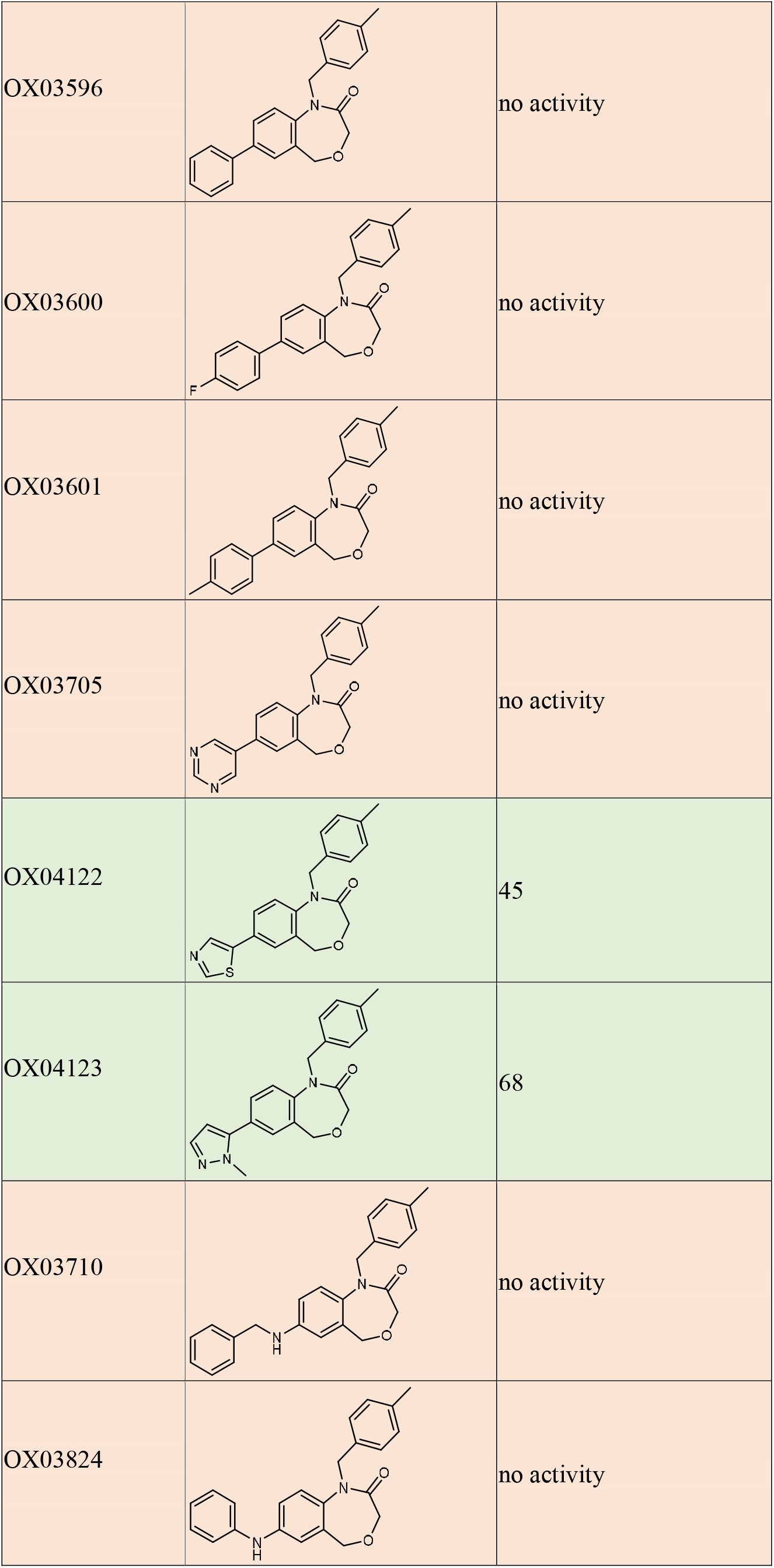
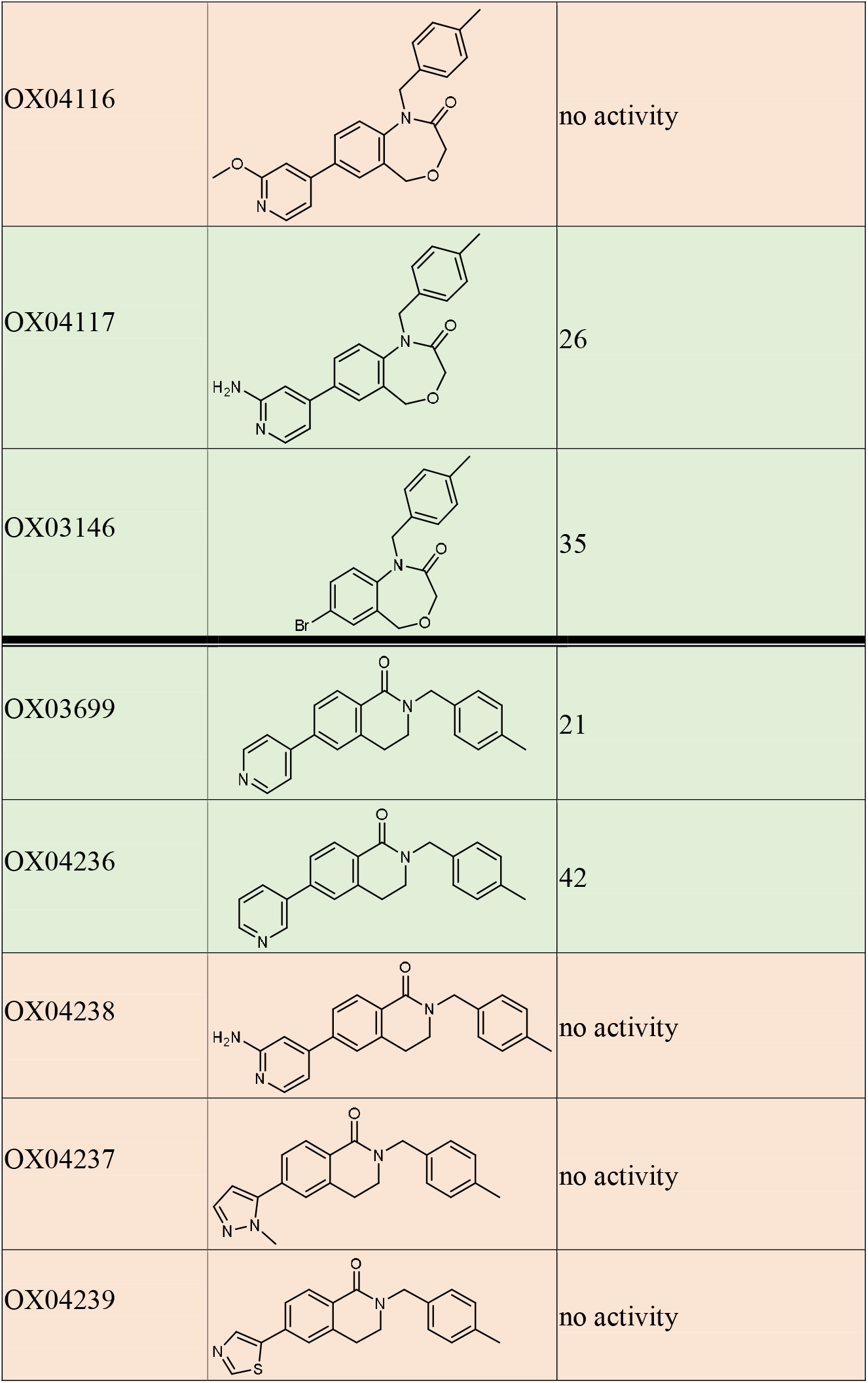
Structures and EC_50_ of representative DHB compounds in the *T. muris* adult motility assay. All compounds investigated in this study are described in the S2 File. No activity means no clear reduction in motility when tested at 100 μM. Where an EC_50_ estimate is shown, it was calculated using a log-logistic model using the R package drc [31].

| Compound | Structure | EC<sub>50</sub> (μM) in *T. muris* adult motility assay |
| --- | --- | --- |
| OX02983 |  | 50 |
| OX03701 |  | no activity |
| OX03707 |  | ≥ 100 |
| OX03704 | | $\geq 75$ |
| OX03825 | | $\geq 75$ |
| OX03144 |  | 26 |
| OX04118 | | $\geq 75$ |
| OX04120 | | $\geq 75$ |
| OX02993 |  | 52 |
| OX04115 |  | 72 |
| OX03153 |  | 57 |
| OX03596 |  | no activity |
| OX03600 |  | no activity |
| OX03601 |  | no activity |
| OX03705 |  | no activity |
| OX04122 |  | 45 |
| OX04123 |  | 68 |
| OX03710 |  | no activity |
| OX03824 |  | no activity |
| OX04116 |  | no activity |
| OX04117 |  | 26 |
| OX03146 |  | 35 |
| OX03699 |  | 21 |
| OX04236 |  | 42 |
| OX04238 |  | no activity |
| OX04237 |  | no activity |
| OX04239 |  | no activity |

Using the appropriate starting materials (see S1 File for details of the syntheses), the different structural analogues **OX03701**, **OX03707** (where the 4-pyridyl ring is in position 8 and 6 of the bicyclic core respectively, see Fig 1) and **OX03704**(the reverse amide equivalent of **OX02983**) were prepared using a similar synthesis to **OX02983** (Table 1). Interestingly, none of the structural analogues exhibited any activity in our *ex vivo* adult *T. muris* motility assay, revealing that the regiochemistry within **OX02983** is important for its activity. The next step was to investigate cycle **C**; a small set of amines was used in the reductive amination step to prepare analogues **OX04118**, **OX04120**, **OX02993**, **OX03825**, **OX03144** bearing methyl, cyclopropyl, cyclohexyl, benzyl and *p*-trifluoromethylbenzyl groups respectively. From those, only the cyclohexyl substituted derivative **OX02993** and the *p*-trifluoromethylbenzyl substituted derivative **OX03144** showed activity in the motility assay, with EC_50_ values of 52 μM and 26 μM respectively. The next step was to vary cycle **D**, while keeping cycles **A**-**C** constant to allow a comparison with **OX02983**. Suzuki reactions were therefore carried out on the 7-bromo precursor with an array of boronic acids and esters. The regioisomers of the pyridyl ring (**D**) were tolerated with *meta* and *para* giving the best activity. Analogues where the pyridyl ring was replaced with an aryl substituent were all inactive, be they unsubstituted (**OX03596**), substituted with an electron withdrawing group (4-F, **OX03600**), or an electron donating group (4-Me, **OX03601**) (Table 1). Different heterocycles were also trialled in place of the pyridine; a similar level of activity was obtained with the isosteric thiazole (**OX04122**, EC_50_ of 45 μM) and the methylimidazole (**OX04123**, EC_50_ of 68 μM) analogues. Substituting with a pyrimidine (**OX03705**) led to a loss of activity, leading us to hypothesize that the basicity of the substituent may be of importance to the activity. Following this, we prepared phenylamine and benzyl amine-substituted analogues **OX03824** and **OX03710**, but neither exhibited activity against *T. muris*, suggesting incorporating a linker between cycles **A** and **D** was not tolerated. We then turned our interest to substituted pyridyl, and although the methoxy substituted pyridyl (**OX04116**) was not active, the amino pyridyl **OX04117** displayed modestly improved activity than **OX02983**(EC_50_ 26 μM), which may be related to its moderately higher basicity.

In an effort to improve the efficacy further, we looked at more drastic modification to core **B**, by contracting the ring by removing the oxygen atom. Forge (Cresset) was used to overlay **OX02983** and its six-membered ring analogue **OX3699**; a good fit was obtained (~79% similarity) suggesting dihydrobenzoquinolinones (DBQ) as possible candidates for further improvement (Fig 2).

**Fig 2.**
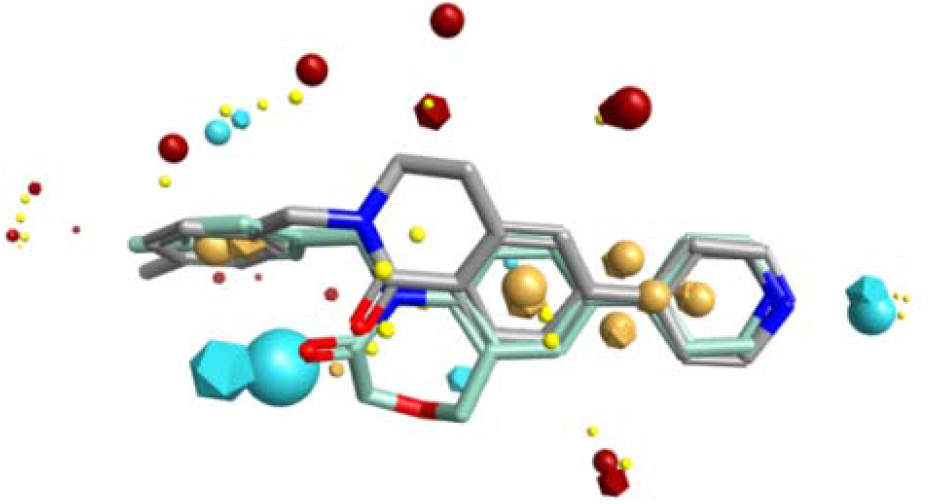
Overlay of **OX02983** (light blue) and **OX3699** (grey); blue spheres represent negative electrostatic field, red spheres represent positive electrostatic field, brown spheres represent hydrophobicity and small yellow sphere represent the van der Waals force.

DBQs have been investigated quite extensively in medicinal chemistry; examples have been reported as antiviral agents through inhibition of HIV replication [39,40]. Other analogues were found to inhibit WDR5 protein-protein interactions, leading to inhibition of cancer cell proliferation [41–43]. ^3-5^

It was decided to prepare a small number of compounds only using those substituents and comparable regiochemistry that gave the most potent analogues so far.

The synthesis started with substitution at *N*2 of 6-bromo-3,4-dihydroisoquinolinone with 4-methylbenzyl bromide, followed by a Suzuki coupling reaction with the requisite boronic acid to afford the desired ring contracted **OX02983** mimic (Scheme 2).

**Scheme 2.**
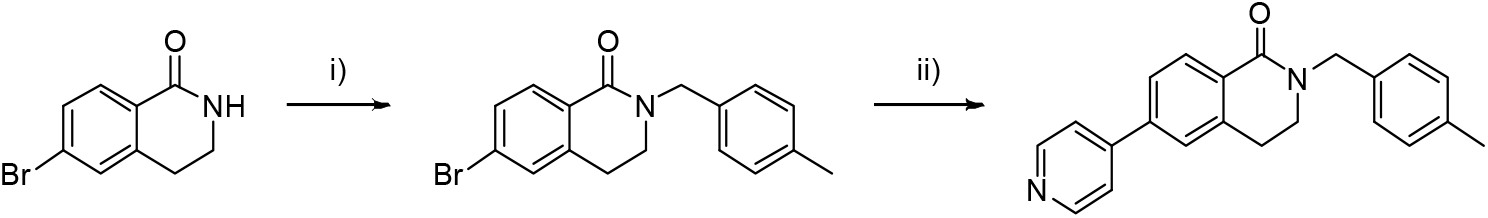
Synthesis of **OX3699**: Reagents and conditions: i) 4-methylbenzyl bromide (2.0 eq.), NaH (1.5 eq.), DMF, rt, 16 h (96 %); ii) 4-pyridyl-B(OH)_2_ (1.1 eq.), Pd(dppf)Cl_2_ (5 mol%), K_2_CO_3_ (3.0 eq.), 1,4-dioxane/H_2_O (4:1), 90 °C 18 h.

The DBQ bearing the 3-and 4-pyridyl substituents (**OX03699** and **OX04236**) were active in the motility assay and led to similar EC_50_s to the best results from the DHB series (with EC_50_ values of 21 μM and 46 μM respectively). Unfortunately, as soon as we moved away from the simple pyridyl substituent, all activity in the motility assay was lost again. The 2-amino pyrid-5-yl, the best example of ring **D** in the DHB series, was surprisingly inactive (**OX04238** EC_50_ >100 μM *vs*. **OX04117** EC_50_ 26 μM). Similarly, the methyl imidazole and the thiazole-substituted analogues (**OX04237** and **OX4739** respectively), also exhibited no activity in the motility assay, in contrast to their DHB counterparts suggesting that SARs did not correlate between the DHB and DBQ series. As the best results from the DQB and the DHB series were largely similar, we felt that this alternative core was not going to enhance substantially the potency of the compounds.

Collectively, these data have improved our understanding or provided insights into the SARs of the DHB/DQB family of compounds. The structure of cycles **A** and **B** in **OX02983** were found to be critical to activity; variations of the toluyl group for ring **C** generally also led to inactive compounds. Some variations of cycle **D** were tolerated, and there appeared to be a preference for a basic site within the substituent. However, although we were able to alter the structure resulting in loss of activity, we were unable to improve, only retain, activity.

Apart from the representative compounds presented in Table 1, further similar analogues and all synthetic precursors were prepared and tested (S2 File). Together it gave us a library of 47 compounds that could then be used against different parasite species to understand whether these compound series showed broad-spectrum anthelmintic activity.

### DHB compounds are active in models of a range of helminth infections

Whipworm is only one of many widely prevalent human helminth infections, and there are continuing efforts to improve drug treatments for these diseases. There have been recent successes, such as the approval of the veterinary medicine moxidectin for onchocerciasis [44], and the establishment of the triple therapy albendazole, diethylcarbamazine citrate plus ivermectin as an improved microfilaricide treatment for lymphatic filariasis suitable for mass drug administration [45,46]. However, sub-optimal efficacy, problematic contraindications, and concerns that mass drug administration could lead to the spread of drug resistance, mean that repurposing of veterinary anthelmintics, improving drug combinations, and the development of new anthelmintics remain priorities [47–49].

Development of a new anthelmintic is a long and expensive process, and funding for neglected tropical diseases is limited. Furthermore, multiple parasitic helminths, for example the soil transmitted nematodes, *Ascaris*, *Trichuris* and the human hookworms, the vector transmitted filarial nematodes, and the *Schistosoma* trematodes, are often endemic in the same regions. It would, therefore, be helpful if a new drug would have activity against several target species and which worked across the Nematoda and Platyhelminthes phyla. We therefore wanted to investigate whether the DHB series of compounds had a range of activities beyond *Trichuris*.

#### Activity against B. malayi

*B. malayi* is one of the tissue dwelling nematode parasites responsible for human lymphatic filariasis [50]. We first examined single dose efficacy of 33 DHB compounds at 10 μM against the *B. malayi* mf larval stage, with motility scored every 24 hours. The results after six days are shown in Fig 3A. Ivermectin was used as a positive control in the mf assay. **OX02983** showed the most promise in this assay, reducing average motility to a score of 1. From this primary screen, 15 compounds that were determined to significantly impact *B. malayi* mf motility, plus an additional 5 with no discernible effect, were retested in a secondary screen (Fig 3B). These results confirmed the significant reduction in motility caused by 11 compounds and confirmed the paralytic effect of **OX02983**. After six days drug exposure, mf were also tested for metabolic activity, a measure of parasite viability, using the MTT assay (Fig 3C). **OX02983** and **OX03153**, in particular, showed activity in this assay, significantly reducing *B. malayi* mf MTT reductase activity on average by 53% and 82%, respectively (1-way ANOVA with Holm-Sidak’s multiple comparison tests, P<0.01 and P<0.0001). To determine the dose-dependent efficacy of **OX02983** and **OX03153**, they were tested in a concentration response 6-day experiment (dose range 0.016-50 μM) using MTT reductase activity as a quantitative viability readout (Fig 3D,E). From this an EC_50_ concentration of 5.5 μM was determined for **OX02983** and 26.7 μM for **OX03153**.

**Fig 3.**
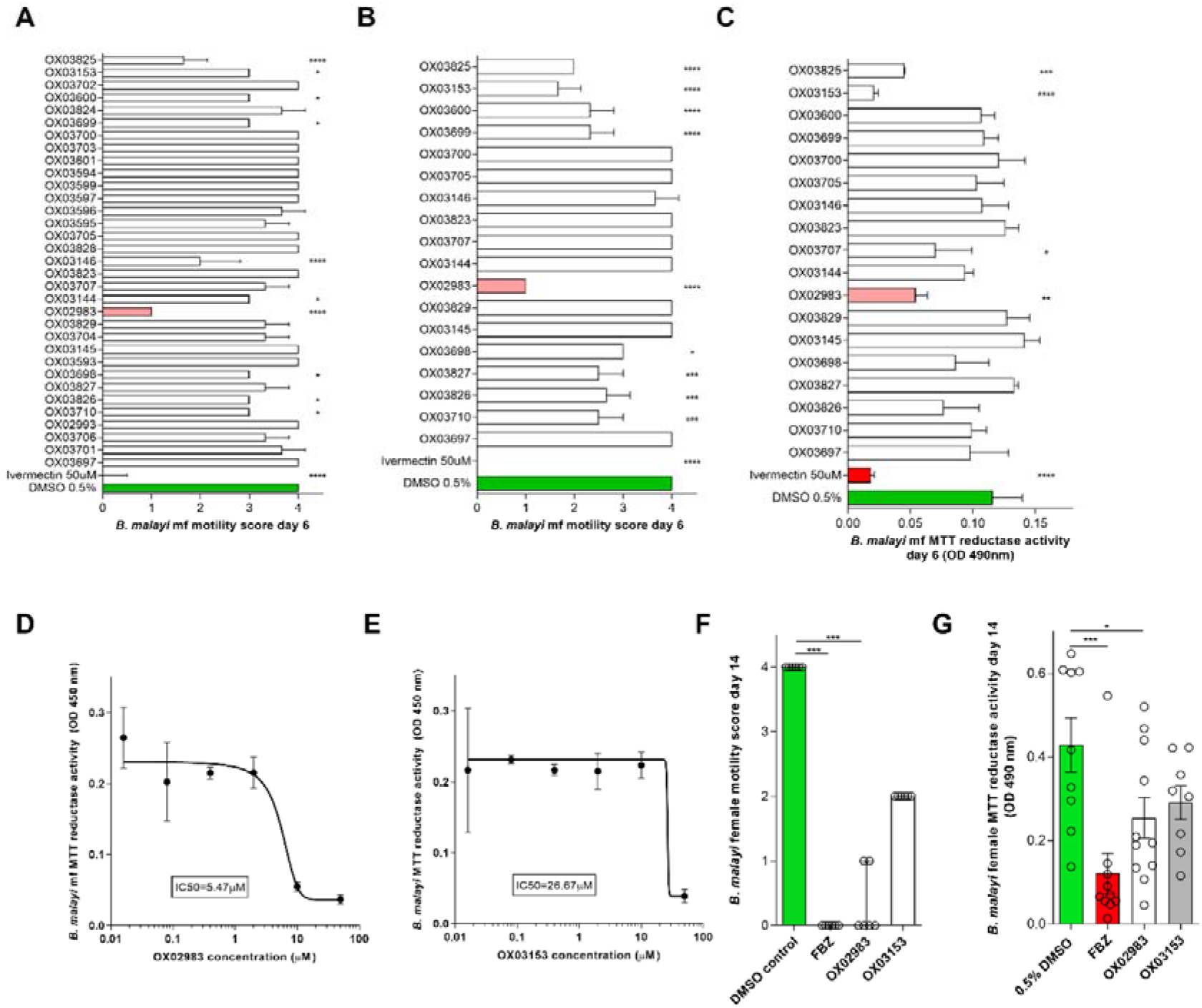
Activity of 33 DHB compounds against *B. malayi* microfilariae and adults. (A) Primary screen – assessment of *B. malayi* mf motility (5 point scoring system) after six days continuous exposure to 35 test compound screened at 10 μM in triplicate. Ivermectin (50 μM) was the positive control. (B) Confirmatory mf motility and (C) metabolic activity screening of 15 active compounds identified in (A) and five inactive compounds (10μM in triplicate). (D, E) 50% inhibitory concentration (EC_50_) assays of active compounds **OX02983** and **OX03153** on *B. malayi* microfilarial metabolic activity after 6-day continuous exposure. Metabolic activity (C-E) was assessed by colormetric MTT assay, data is optical density of mf extracts measured at 490nm. (F) Effects on adult female *B. malayi* motility and (G) metabolic activity following 14 day continuous exposure to **OX02983** or **OX03153**(10 μM). Flubendazole (10 μM) was used as a positive control in the assay. Data plotted is mean ± SD of 3 replicates (A-E) median and range of 6 replicates (F) and mean ± SEM of 9-11 replicates (G). Significant differences were determined by 1-way ANOVA with Holm-Sidak multiple comparisons test (A-C and G) or Kruskal-Wallis with Dunn’s multiple comparisons test (F). Significance is indicated ****P<0.0001, ***P<0.001, **P<0.01 and *P<0.05.

Due to their efficacy against *B. malayi* mf, **OX02983** and **OX03153** were advanced for *in vitro* activity against adult *B. malayi*, utilising a novel, long-term adult worm lymphatic endothelial cell bilayer co-culture system. Adult female *B. malayi* exposed to vehicle control retained full survival and motility in culture over 14 days whereas the positive control, flubendazole (10 μM) mediated complete paralytic activity by day 14 (Kruskal Wallis with Dunn’s multiple comparisons tests, P<0.001) and significantly reduced metabolic activity by an average of 72% (1-way ANOVA with Holm-Sidak’s multiple comparison tests, P<0.001) (Fig 3F-G). **OX02983**(10 μM) also mediated significant anti-filarial activities against adult *B. malayi* by day 14. Motility was completely hindered in 4/6 adult parasites by **OX02983**(Kruskal Wallis with Dunn’s multiple comparisons tests, P<0.001), whilst **OX03153** mediated a 50% partial reduction in adult motility. OX02983 also significantly impacted on adult female *B. malayi* metabolic activity, on average by 41% (1-way ANOVA with Holm-Sidak’s multiple comparison tests, P<0.05). Taken together, these results are encouraging because they show that compounds that are active against *T. muris* (a clade I nematode according to the phylogeny of Blaxter) are also active against evolutionarily-distant nematodes, as *B. malayi* is a clade III nematode [51].

#### Activity against H. polygyrus

*H. polygyrus bakeri* is an intestinal nematode parasite of laboratory mice [52]. It is a strongylid nematode, related to human hookworm species. 31 DHB compounds were tested at 100 μM against *ex vivo H. polygyrus* L3 stage worms (n = 2). The results are shown in Fig 4. The cut-off used to determine hits in this assay is 50% larval death [17]. Two compounds, **OX03594** and **OX03599**, exceeded this level of larval death and were therefore considered active. They did not however reach the threshold for good activity (75%). Given the modest activity of these compounds against *H. polygyrus* we have not further pursued this direction at this point. Activity of DHB compounds against nematodes in three of the five clades of the phylum Nematoda, according to the phylogeny of Blaxter, supports the potential for development of a pan-nematode control agent from this compound series [51].

**Fig 4.**
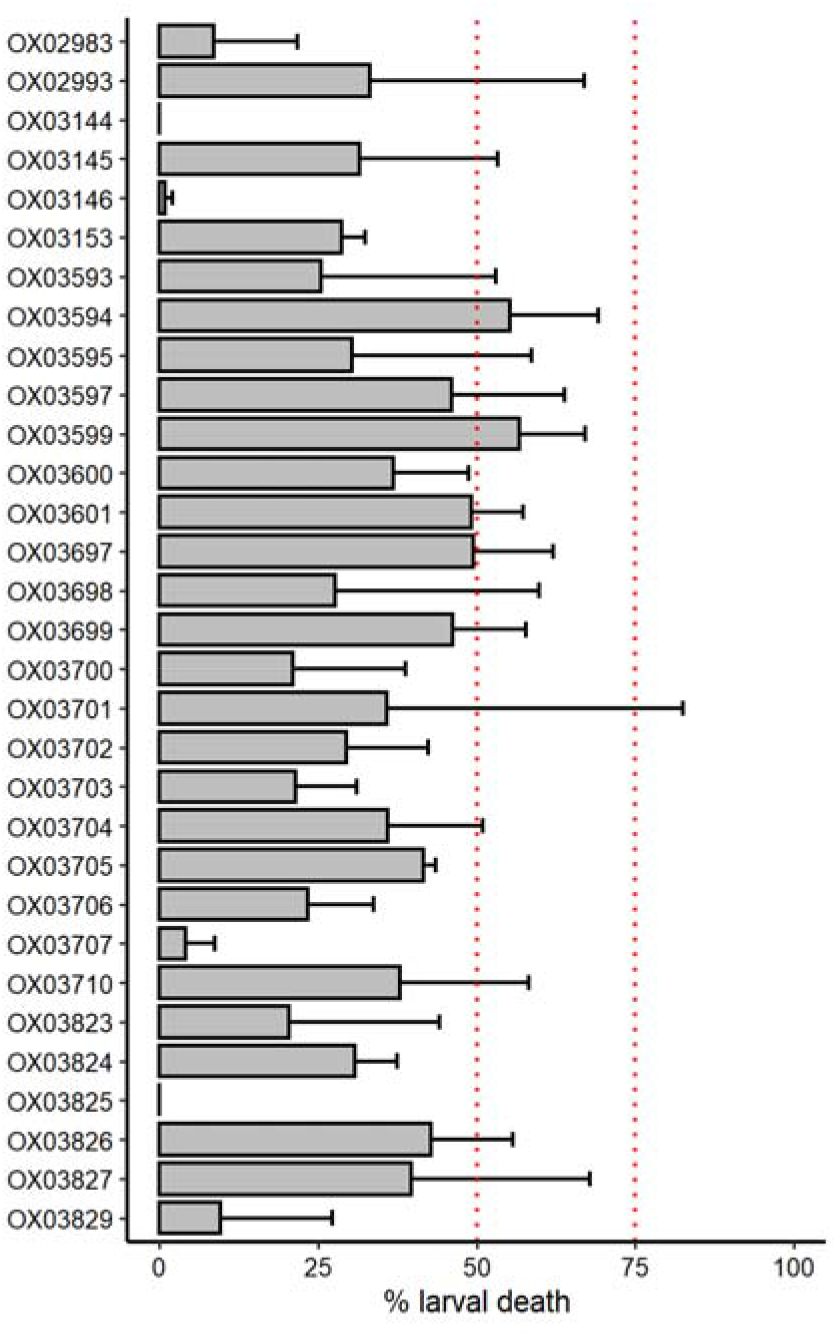
Measurement of the activity of 31 DHB compounds against *H. polygyrus* L3 stage worms. Larval death is measured as the proportion of worms that respond to stimulus. Compounds were tested in duplicate at 100 μM. Dashed lines indicates the cut-off (50%) used to determine hits in this assay and the cut-off for good activity (75%) [17]: <50%, not active, 50-75% moderate activity, >75% good and >90% excellent activity.

#### Activity against S. mansoni schistosomula

Compared to *T. muris, H. polygyrus* and *B. malayi,* which are all parasitic worms within the phylum Nematoda, *S. mansoni* is a more evolutionary distinct helminth – a trematode within the Platyhelminthes phylum. It is a human parasite that infects around 150 million people, causing schistosomiasis [1]. Praziquantel, an N-acylated quinoline-piperazinone, is the basis of schistosomiasis treatment and is safe and efficacious against adult worms of all *Schistosoma* spp. as a monotherapy. However, there is concern about the emergence of drug resistance [53] and praziquantel has lower efficacy against juvenile forms, so immature parasites may survive drug exposure and continue the infection.

We screened 30 DHB compounds against *S. mansoni* schistosomula at 50 μM using the RoboWorm system. This is an imaging-based screen that measures two parameters, motility and “phenotype,” an assessment of morphological and other features [37]. Auranofin, an inhibitor of *S. mansoni* thioredoxin glutathione reductase (TGR) activity [54] was the positive control in this experiment. The results are shown in Fig 5. The cut-offs for defining hit compounds in this assay have been previously defined [37,55]. Nine compounds were hits in this assay for both motility and phenotype measurements. Concentration-response curves were measured for these compounds (Table 2), with EC_50_ values in the range 14-41 μM. It is encouraging that DHB compound series members show activity against such evolutionarily distant pathogens to whipworm, particularly as DHB compounds show little or no cytotoxicity in mammalian cell culture – so these compounds are not broadly toxic [18].

**Fig 5.**
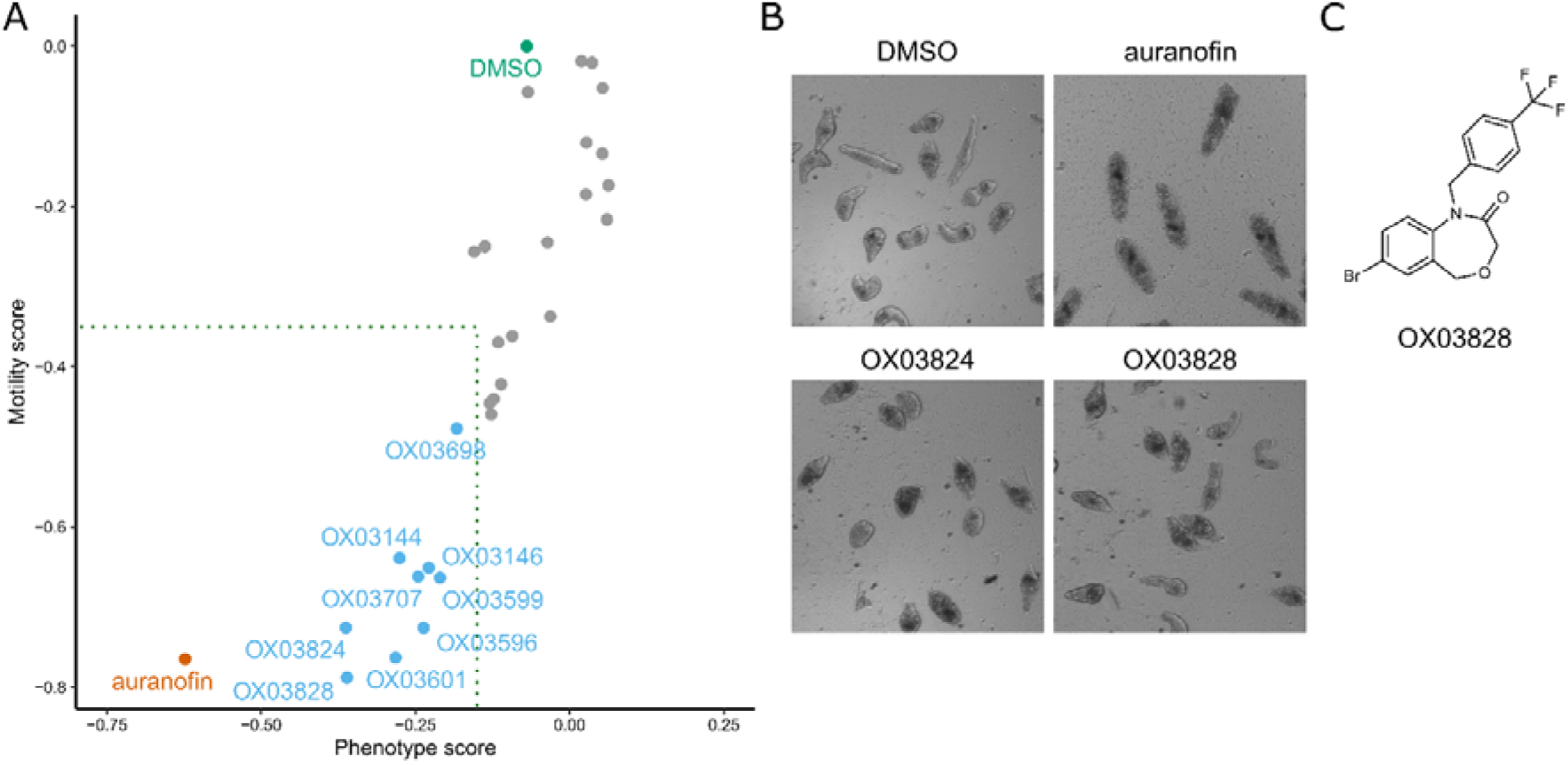
Measurement of the activity of 30 DHB compounds against *S. mansoni* schistosomula using the RoboWorm platform. (A) Each point is the measured effect of one compound on the two parameters – motility and phenotype. The phenotype score is calculated by a computational model that assesses morphological and texture properties of the schistosomula [37]. Compounds were screened at 50 μM. Auranofin was the positive control (screened at 10 μM). Dotted box indicates the threshold for activity in this assay: −0.15 for phenotype and −0.35 for motility; compounds must be below this threshold for both parameters to be considered a hit [37,55]. Compounds were screened in duplicate on two or three separate occasions and the data represents the average scores of these experiments. (B) Representative images of schistosomula treated with controls, **OX03824** and **OX03828**. (C) Structure of **OX03828**. The structure of **OX03824** is shown in Table 1.

**Table 2.** EC_50_ values for compounds active in the *S. mansoni* schistosomula Roboworm assay. Compounds were screened at 10, 20, 30, 40 and 50 μM and EC_50_ values were calculated for each of the screening parameters, phenotype and motility.

| Compound | EC<sub>50</sub> in *S. mansoni* schistosomula Roboworm assay |  |
| --- | --- | --- |
| | (μM) | |
|  | phenotype score | motility score |
| OX03144 | 28.4 | 28.0 |
| OX03146 | 25.5 | 25.6 |
| OX03596 | 29.1 | 26.8 |
| OX03599 | 14.2 | 18.3 |
| OX03601 | 24.4 | 20.6 |
| OX03698 | 28.5 | 26.6 |
| OX03707 | 32.7 | 29.3 |
| OX03824 | 40.9 | 36.8 |
| OX03828 | 25.7 | 20.7 |

## Discussion

### Investigation of the DHB structure-activity relationship

We previously identified a small hit series of five DHB compounds with activity against *T. muris* adult motility [18]. In medicinal chemistry, it is important to understand how variations in the structure of the compound affect activity, as this allows us to discover the critical aspects of the compound for target binding, with the overall aim of increasing potency as well as improving physicochemical properties. We therefore embarked upon a systematic, structure-activity relationship investigation, taking advantage of the convenient synthesis of the DHBs, which allowed us to systematically alter the four cyclic components of this class of compounds. A total of 47 variant compounds were synthesised in this work.

This work has enabled us to define certain essential features of the anti-whipworm DHB compounds. The 4-pyridyl ring (cycle D in Fig 1) must be in the 7 position, unlike the analogues **OX03701** and **OX03707**. The amide moiety of the oxazepinone ring must be as in **OX02983**, and not as in **OX03704**. The oxazepinone nitrogen can be substituted with methylbenzyl, cyclohexyl and *p-*trifluromethylbenzyl (**OX02983**, **OX02993**, and **OX03144**), but not methyl, cyclopropyl, or benzyl groups. We also investigated in detail the replacement of cycle B. We found that removal of the oxygen from the DHB core was also consistent with similar activity to **OX02983**– the dihydrobenzoquinolinone compounds **OX03699** and **OX04236** had EC_50_ values of 21 and 42 μM respectively.

### Targeting multiple helminth species with DHB family members

Despite being unable to improve efficacy against *Trichuris* substantially through structural modifications, we were able to demonstrate activity of our compounds against other helminth parasites. In drug discovery for NTDs, pan-anthelmintic activity is desirable given that polyparasitism in the target population is the norm. Thus, being able to target multiple species of helminths with a single drug administered via mass drug administration programmes is of significant benefit. Of particular note was the commonality in DHB compounds active against *T. muris* that were also active against the tissue dwelling nematode parasite *B. malayi.* The ability of the DHB compounds to act against different clades within the nematode phylum is not unprecedented, indeed the co-administration of albendazole with ivermectin is currently advocated for control of *Trichuris*, and the same drug combination (in some situations supplemented with diethylcarbamazine) is widely used against lymphatic filariasis [46]. Indeed, the large-scale efforts to treat lymphatic filariasis have indirectly enhanced the number of people being treated for soil transmitted helminths [56]. Similarly, the alternative drug combination of albendazole and moxidectin is also being explored for the treatment of Trichuriasis given that moxidectin is an approved treatment for onchocerciasis [57].

In contrast, there are currently no drugs used in MDA that have demonstrated cross-phyla efficacy against both schistosomes and nematodes. Currently only praziquantel is used for preventative chemotherapy against schistosomes, although co-administration with albendazole is recommended where STHs are co-endemic [58]. Therefore it was notable that DHB family members could work across phyla, showing some activity against both schistosomes and nematodes.

### Conclusions

In this study we have investigated the structure-activity relationship of the DHB compounds, defined essential features for anthelmintic action, and broadened the active series by the discovery of dihydrobenzoquinolinone compounds with activity against *T. muris* adult motility. We have also demonstrated that DHB and related compounds have activity against multiple helminths across different phyla – against the nematodes *B. malayi* and *H. polygyrus* as well as *T. muris,* and against the trematode *S. mansoni.* What we have not achieved however, is the substantive improvement in potency from the 20-50 μM range that would be desirable to progress this series with confidence to *in vivo* testing. Open science, where information is disclosed more freely than in traditional models, is proposed to accelerate drug discovery and make it more cost efficient, especially in the context of neglected diseases [59,60]. We have therefore decided to report our progress at this point. We note that we do not yet know the target of the DHB/DBQ compounds in helminths. Identifying this target may facilitate the boost in activity we are striving for.

## Acknowledgments

JDT, MJT and AEM were supported by a National Centre for Replacement, Refinement and Reduction of Animals in Research Studentship (NC3R Studentship NC/M00175X/1) and a Bill and Melinda Gates Foundation award to the Liverpool School of Tropical Medicine (BMGF OPP1054324).

FAP and DBS are supported by Medical Research Council grant MR/N024842/1 and a UCL/Wellcome Trust Translational Partnership Pilot Grant.

JFT, HW and KFH acknowledge the Welsh Government Life Sciences Research Network Wales and the Wellcome Trust Pathfinder (201008/Z/16/Z) schemes for financial support of the Roboworm platform.

## Supporting information Captions

**S1 File. Supporting information for synthetic chemistry**

**S2 File. Summary table of compound structures and assay results.** Shaded results are those compounds that were active in each assay.

## References

1. GBD 2017 Disease and Injury Incidence and Prevalence Collaborators. Global, regional, and national incidence, prevalence, and years lived with disability for 354 diseases and injuries for 195 countries and territories, 1990-2017: a systematic analysis for the Global Burden of Disease Study 2017. Lancet. 2018;392: 1789–1858. doi:10.1016/S0140-6736(18)32279-7

2. Kyu HH, Abate D, Abate KH, Abay SM, Abbafati C, Abbasi N, et al. Global, regional, and national disability-adjusted life-years (DALYs) for 359 diseases and injuries and healthy life expectancy (HALE) for 195 countries and territories, 1990–2017: a systematic analysis for the Global Burden of Disease Study 2017. The Lancet. 2018;392: 1859–1922. doi:10.1016/S0140-6736(18)32335-3

3. World Health Organization. 2030 targets for soil-transmitted helminthiases control programmes. 2020 [cited 21 Apr 2020]. Available: http://www.who.int/intestinal_worms/resources/9789240000315/en/

4. Palmeirim MS, Ame SM, Ali SM, Hattendorf J, Keiser J. Efficacy and Safety of a Single Dose versus a Multiple Dose Regimen of Mebendazole against Hookworm Infections in Children: A Randomised, Double-blind Trial. EClinicalMedicine. 2018;1: 7–13. doi:10.1016/j.eclinm.2018.06.004

5. Keiser J, Utzinger J. Efficacy of current drugs against soil-transmitted helminth infections: Systematic review and meta-analysis. JAMA. 2008;299: 1937–1948. doi:10.1001/jama.299.16.1937

6. Moser W, Schindler C, Keiser J. Drug Combinations Against Soil-Transmitted Helminth Infections. Adv Parasitol. 2019;103: 91–115. doi:10.1016/bs.apar.2018.08.002

7. Barda B, Ame SM, Ali SM, Albonico M, Puchkov M, Huwyler J, et al. Efficacy and tolerability of moxidectin alone and in co-administration with albendazole and tribendimidine versus albendazole plus oxantel pamoate against *Trichuris trichiura* infections: a randomised, non-inferiority, single-blind trial. Lancet Infect Dis. 2018;18: 864–873. doi:10.1016/S1473-3099(18)30233-0

8. Speich B, Ame SM, Ali SM, Alles R, Huwyler J, Hattendorf J, et al. Oxantel Pamoate–Albendazole for *Trichuris trichiura* Infection. New England Journal of Medicine. 2014;370: 610–620. doi:10.1056/NEJMoa1301956

9. Speich B, Ali SM, Ame SM, Bogoch II, Alles R, Huwyler J, et al. Efficacy and safety of albendazole plus ivermectin, albendazole plus mebendazole, albendazole plus oxantel pamoate, and mebendazole alone against *Trichuris trichiura* and concomitant soil-transmitted helminth infections: a four-arm, randomised controlled trial. The Lancet Infectious Diseases. 2015;15: 277–284. doi:10.1016/S1473-3099(14)71050-3

10. Speich B, Moser W, Ali SM, Ame SM, Albonico M, Hattendorf J, et al. Efficacy and reinfection with soil-transmitted helminths 18-weeks post-treatment with albendazole-ivermectin, albendazole-mebendazole, albendazole-oxantel pamoate and mebendazole. Parasit Vectors. 2016;9: 123. doi:10.1186/s13071-016-1406-8

11. Moser W, Schindler C, Keiser J. Efficacy of recommended drugs against soil transmitted helminths: systematic review and network meta-analysis. BMJ. 2017;358: j4307. doi:10.1136/bmj.j4307

12. Diawara A, Drake LJ, Suswillo RR, Kihara J, Bundy DAP, Scott ME, et al. Assays to Detect β-Tubulin Codon 200 Polymorphism in *Trichuris trichiura* and *Ascaris lumbricoides*. PLoS Negl Trop Dis. 2009;3: e397. doi:10.1371/journal.pntd.0000397

13. Diawara A, Halpenny CM, Churcher TS, Mwandawiro C, Kihara J, Kaplan RM, et al. Association between Response to Albendazole Treatment and β-Tubulin Genotype Frequencies in Soil-transmitted Helminths. PLoS Negl Trop Dis. 2013;7: e2247. doi:10.1371/journal.pntd.0002247

14. Vlaminck J, Cools P, Albonico M, Ame S, Chanthapaseuth T, Viengxay V, et al. Piloting a surveillance system to monitor the global patterns of drug efficacy and the emergence of anthelmintic resistance in soil-transmitted helminth control programs: a Starworms study protocol. Gates Open Res. 2020;4: 28. doi:10.12688/gatesopenres.13115.1

15. Elfawal MA, Savinov SN, Aroian RV. Drug Screening for Discovery of Broad-spectrum Agents for Soil-transmitted Nematodes. Sci Rep. 2019;9: 12347. doi:10.1038/s41598-019-48720-1

16. Hurst RJ, Hopwood T, Gallagher AL, Partridge FA, Burgis T, Sattelle DB, et al. An antagonist of the retinoid X receptor reduces the viability of *Trichuris muris* in vitro. BMC Infectious Diseases. 2014;14: 520. doi:10.1186/1471-2334-14-520

17. Keiser J, Panic G, Adelfio R, Cowan N, Vargas M, Scandale I. Evaluation of an FDA approved library against laboratory models of human intestinal nematode infections. Parasit Vectors. 2016;9: 376. doi:10.1186/s13071-016-1616-0

18. Partridge FA, Murphy EA, Willis NJ, Bataille CJR, Forman R, Heyer-Chauhan N, et al. Dihydrobenz[*e*][1,4]oxazepin-2(3*H*)-ones, a new anthelmintic chemotype immobilising whipworm and reducing infectivity *in vivo*. PLoS Negl Trop Dis. 2017;11: e0005359. doi:10.1371/journal.pntd.0005359

19. Partridge FA, Forman R, Willis NJ, Bataille CJR, Murphy EA, Brown AE, et al. 2,4-Diaminothieno[3,2-*d*]pyrimidines, a new class of anthelmintic with activity against adult and egg stages of whipworm. PLoS Negl Trop Dis. 2018;12: e0006487. doi:10.1371/journal.pntd.0006487

20. Preston S, Jiao Y, Baell JB, Keiser J, Crawford S, Koehler AV, et al. Screening of the “Open Scaffolds” collection from Compounds Australia identifies a new chemical entity with anthelmintic activities against different developmental stages of the barber’s pole worm and other parasitic nematodes. Int J Parasitol Drugs Drug Resist. 2017;7: 286–294. doi:10.1016/j.ijpddr.2017.05.004

21. Tyagi R, Maddirala AR, Elfawal M, Fischer C, Bulman CA, Rosa BA, et al. Small Molecule Inhibitors of Metabolic Enzymes Repurposed as a New Class of Anthelmintics. ACS Infect Dis. 2018;4: 1130–1145. doi:10.1021/acsinfecdis.8b00090

22. Weeks JC, Roberts WM, Leasure C, Suzuki BM, Robinson KJ, Currey H, et al. Sertraline, Paroxetine, and Chlorpromazine Are Rapidly Acting Anthelmintic Drugs Capable of Clinical Repurposing. Sci Rep. 2018;8: 975. doi:10.1038/s41598-017-18457-w

23. World Health Organization. Ending the neglect to attain the Sustainable Development Goals: A road map for neglected tropical diseases 2021–2030. World Health Organization; 2020. Available: http://www.who.int/neglected_diseases/resources/who-ucn-ntd-2020.01/en/

24. Boussinesq M, Fobi G, Kuesel AC. Alternative treatment strategies to accelerate the elimination of onchocerciasis. Int Health. 2018;10: i40–i48. doi:10.1093/inthealth/ihx054

25. Taylor MJ, Hoerauf A, Townson S, Slatko BE, Ward SA. Anti-*Wolbachia* drug discovery and development: safe macrofilaricides for onchocerciasis and lymphatic filariasis. Parasitology. 2014;141: 119–127. doi:10.1017/S0031182013001108

26. Sahputra R, Ruckerl D, Couper KN, Muller W, Else KJ. The Essential Role Played by B Cells in Supporting Protective Immunity Against *Trichuris muris* Infection Is by Controlling the Th1/Th2 Balance in the Mesenteric Lymph Nodes and Depends on Host Genetic Background. Front Immunol. 2019;10: 2842. doi:10.3389/fimmu.2019.02842

27. Wakelin D. Acquired immunity to *Trichuris muris* in the albino laboratory mouse. Parasitology. 1967;57: 515–524. doi:10.1017/s0031182000072395

28. Partridge FA, Brown AE, Buckingham SD, Willis NJ, Wynne GM, Forman R, et al. An automated high-throughput system for phenotypic screening of chemical libraries on *C. elegans* and parasitic nematodes. International Journal for Parasitology: Drugs and Drug Resistance. 2018;8: 8–21. doi:10.1016/j.ijpddr.2017.11.004

29. Buckingham SD, Partridge FA, Poulton BC, Miller BS, McKendry RA, Lycett GJ, et al. Automated phenotyping of mosquito larvae enables high-throughput screening for novel larvicides and smartphone-based detection of larval insecticide resistance. bioRxiv. 2020; 2020.07.20.211946. doi:10.1101/2020.07.20.211946

30. Buckingham SD, Partridge FA, Sattelle DB. Automated, high-throughput, motility analysis in *Caenorhabditis elegans* and parasitic nematodes: Applications in the search for new anthelmintics. International Journal for Parasitology: Drugs and Drug Resistance. 2014;4: 226–232. doi:10.1016/j.ijpddr.2014.10.004

31. Ritz C, Baty F, Streibig JC, Gerhard D. Dose-Response Analysis Using R. PLOS ONE. 2015;10: e0146021. doi:10.1371/journal.pone.0146021

32. Pionnier N, Sjoberg H, Furlong-Silva J, Marriott A, Halliday A, Archer J, et al. Eosinophil-Mediated Immune Control of Adult Filarial Nematode Infection Can Proceed in the Absence of IL-4 Receptor Signaling. The Journal of Immunology. 2020;205: 731–740. doi:10.4049/jimmunol.1901244

33. Turner JD, Pionnier N, Furlong-Silva J, Sjoberg H, Cross S, Halliday A, et al. Interleukin-4 activated macrophages mediate immunity to filarial helminth infection by sustaining CCR3-dependent eosinophilia. PLOS Pathogens. 2018;14: e1006949. doi:10.1371/journal.ppat.1006949

34. Geyer KK, Niazi UH, Duval D, Cosseau C, Tomlinson C, Chalmers IW, et al. The *Biomphalaria glabrata* DNA methylation machinery displays spatial tissue expression, is differentially active in distinct snail populations and is modulated by interactions with *Schistosoma mansoni*. PLoS Negl Trop Dis. 2017;11: e0005246. doi:10.1371/journal.pntd.0005246

35. Colley DG, Wikel SK. *Schistosoma mansoni:* simplified method for the production of schistosomules. Exp Parasitol. 1974;35: 44–51. doi:10.1016/0014-4894(74)90005-8

36. Crusco A, Bordoni C, Chakroborty A, Whatley KCL, Whiteland H, Westwell AD, et al. Design, synthesis and anthelmintic activity of 7-keto-sempervirol analogues. Eur J Med Chem. 2018;152: 87–100. doi:10.1016/j.ejmech.2018.04.032

37. Paveley RA, Mansour NR, Hallyburton I, Bleicher LS, Benn AE, Mikic I, et al. Whole Organism High-Content Screening by Label-Free, Image-Based Bayesian Classification for Parasitic Diseases. PLOS Neglected Tropical Diseases. 2012;6: e1762. doi:10.1371/journal.pntd.0001762

38. Zhang J, Chung T, Oldenburg K. A Simple Statistical Parameter for Use in Evaluation and Validation of High Throughput Screening Assays. J Biomol Screen. 1999;4: 67–73. doi:10.1177/108705719900400206

39. Johns BA, Velthuisen EJ, Weatherhead JG. Tetrahydroisoquinoline derivatives. WO2017093938A1, 2017. Available: https://patents.google.com/patent/WO2017093938A1/fi

40. Zhao XZ, Metifiot M, Smith SJ, Maddali K, Marchand C, Hughes SH, et al. 6,7-Dihydroxyisoindolin-1-one and 7,8-Dihydroxy-3,4-Dihydroisoquinolin-1(2H)-one Based HIV-1 Integrase Inhibitors. Curr Top Med Chem. 2016;16: 435–440. doi:10.2174/1568026615666150813150058

41. Shen J, Xiong B, Zhao Y, Li J, Geng M, Ma L, et al. 1,2,3,4-Tetrahydroisoquinolin-1(H)-one and 1(2H)-phthalazinone derivatives as histone deacetylase inhibitors and their preparation, pharmaceutical compositions and use in the treatment of cancer. CN107879975A, 2018.

42. Lee T, Alvarado JR, Tian J, Meyers KM, Han C, Mills JJ, et al. Wdr5 inhibitors and modulators. WO2020086857A1, 2020.

43. Tian J, Teuscher KB, Aho ER, Alvarado JR, Mills JJ, Meyers KM, et al. Discovery and Structure-Based Optimization of Potent and Selective WD Repeat Domain 5 (WDR5) Inhibitors Containing a Dihydroisoquinolinone Bicyclic Core. J Med Chem. 2020;63: 656–675. doi:10.1021/acs.jmedchem.9b01608

44. Opoku NO, Bakajika DK, Kanza EM, Howard H, Mambandu GL, Nyathirombo A, et al. Single dose moxidectin versus ivermectin for *Onchocerca volvulus* infection in Ghana, Liberia, and the Democratic Republic of the Congo: a randomised, controlled, double-blind phase 3 trial. Lancet. 2018;392: 1207–1216. doi:10.1016/S0140-6736(17)32844-1

45. King CL, Suamani J, Sanuku N, Cheng Y-C, Satofan S, Mancuso B, et al. A Trial of a Triple-Drug Treatment for Lymphatic Filariasis. New England Journal of Medicine. 2018. doi:10.1056/NEJMoa1706854

46. World Health Organization. Guideline: Alternative mass drug administration regimens to eliminate lymphatic filariasis. 2017. Available: http://whqlibdoc.who.int/publications/2011/9789241501811_eng.pdf

47. Kuesel AC. Research for new drugs for elimination of onchocerciasis in Africa. Int J Parasitol Drugs Drug Resist. 2016;6: 272–286. doi:10.1016/j.ijpddr.2016.04.002

48. Vercruysse J, Albonico M, Behnke JM, Kotze AC, Prichard RK, McCarthy JS, et al. Is anthelmintic resistance a concern for the control of human soil-transmitted helminths? International Journal for Parasitology: Drugs and Drug Resistance. 2011;1: 14–27. doi:10.1016/j.ijpddr.2011.09.002

49. Wang W, Wang L, Liang Y-S. Susceptibility or resistance of praziquantel in human schistosomiasis: a review. Parasitol Res. 2012;111: 1871–1877. doi:10.1007/s00436-012-3151-z

50. Taylor MJ, Hoerauf A, Bockarie M. Lymphatic filariasis and onchocerciasis. The Lancet. 2010;376: 1175–1185. doi:10.1016/S0140-6736(10)60586-7

51. Blaxter ML, Ley PD, Garey JR, Liu LX, Scheldeman P, Vierstraete A, et al. A molecular evolutionary framework for the phylum Nematoda. Nature. 1998;392: 71–75. doi:10.1038/32160

52. Behnke JM, Menge DM, Noyes H. *Heligmosomoides bakeri:* a model for exploring the biology and genetics of resistance to chronic gastrointestinal nematode infections. Parasitology. 2009;136: 1565–1580. doi:10.1017/S0031182009006003

53. Crellen T, Walker M, Lamberton PHL, Kabatereine NB, Tukahebwa EM, Cotton JA, et al. Reduced Efficacy of Praziquantel Against *Schistosoma mansoni* Is Associated With Multiple Rounds of Mass Drug Administration. Clin Infect Dis. 2016;63: 1151–1159. doi:10.1093/cid/ciw506

54. Kuntz AN, Davioud-Charvet E, Sayed AA, Califf LL, Dessolin J, Arnér ESJ, et al. Thioredoxin Glutathione Reductase from *Schistosoma mansoni:* An Essential Parasite Enzyme and a Key Drug Target. PLOS Med. 2007;4: e206. doi:10.1371/journal.pmed.0040206

55. Whatley KCL, Padalino G, Whiteland H, Geyer KK, Hulme BJ, Chalmers IW, et al. The repositioning of epigenetic probes/inhibitors identifies new anti-schistosomal lead compounds and chemotherapeutic targets. PLOS Neglected Tropical Diseases. 2019;13: e0007693. doi:10.1371/journal.pntd.0007693

56. Farrell SH, Coffeng LE, Truscott JE, Werkman M, Toor J, de Vlas SJ, et al. Investigating the Effectiveness of Current and Modified World Health Organization Guidelines for the Control of Soil-Transmitted Helminth Infections. Clin Infect Dis. 2018;66: S253–S259. doi:10.1093/cid/ciy002

57. Keller L, Palmeirim MS, Ame SM, Ali SM, Puchkov M, Huwyler J, et al. Efficacy and Safety of Ascending Dosages of Moxidectin and Moxidectin-albendazole Against *Trichuris trichiura* in Adolescents: A Randomized Controlled Trial. Clin Infect Dis. 2020;70: 1193–1201. doi:10.1093/cid/ciz326

58. World Health Organization. Preventive chemotherapy in human helminthiasis. Coordinated use of anthelminthic drugs in control interventions: a manual for health professionals and programme managers. 2006. Available: https://apps.who.int/iris/handle/10665/43545

59. Partridge FA, Forman R, Bataille CJR, Wynne GM, Nick M, Russell AJ, et al. Anthelmintic drug discovery: target identification, screening methods and the role of open science. Beilstein J Org Chem. 2020;16: 1203–1224. doi:10.3762/bjoc.16.105

60. Todd MH. Six Laws of Open Source Drug Discovery. ChemMedChem. 2019;14: 1804–1809. doi:10.1002/cmdc.201900565

