## Supplementary material for "Structural requirements for dihydrobenzoxazepinone anthelmintics: actions against medically important and model parasites - *Trichuris muris*, *Brugia malayi*, *Heligmosomoides polygyrus* and *Schistosoma mansoni*": S1 File

### S1 File. Supporting information for synthetic chemistry

#### General Procedures

**General Procedure 1**

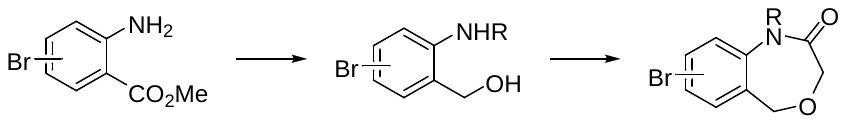

To a solution of the requisite aniline (1.0 eq.) in CH_2_Cl_2_ (6 mL/mmol) at room temperature was added the desired aldehyde (1.5 eq.) and AcOH (0.5 eq.). The resulting solution was then cooled to 0 °C before addition of NaBH(OAc)_3_ (2.0 eq.). The mixture was warmed slowly to room temperature and stirred for 16 h before addition of CH_2_Cl_2_ (5 mL/mmol) and NaHCO_3_ (sat. aq., 10 mL/mmol), and the aqueous layer extracted with CH_2_Cl_2_ (5x10 mL/mmol). The combined organic layers were dried (Na_2_SO_4_) and concentrated *in vacuo* and the crude residue was purified by column chromatography on silica gel to afford the desired methyl ester. A solution of LiAlH_4_ (1.0 M in THF, 3.5 eq.) was added dropwise over a period of 5-10 min to an ice-cold solution of the requisite methyl ester (1.0 eq.) in THF (7 mL/mmol), and the reaction mixture was stirred for 1 h. NH_4_Cl (sat. aq. sol, 3 mL/mmol) was then added and the aqueous layer extracted with EtOAc (3 x 35 mL/mmol), the combined organic layers were dried (Na_2_SO_4_) and concentrated *in vacuo*. The resulting alcohol was used in the next step without further purification. To an ice-cold solution of desired alcohol in THF (2 mL/mmol) was added Et_3_N (2.0 eq.), followed by chloroacetyl chloride (4.4 eq.). The reaction was warmed to room temperature and stirred for 16 h, at which time the crude reaction mixture was passed through a short pad of silica gel, eluted with EtOAc and the solution concentrated *in vacuo*. The crude residue was then dissolved in *^i^*PrOH (2 mL/mmol) before addition of NaOH (10 N, aq., 10 eq.). The reaction was stirred at room temperature for 2 h, diluted with CH_2_Cl_2_ (5 mL/mmol), washed with brine (ca 5 mL/mmol) and further extracted with CH_2_Cl_2_ (5 mL/mmol). The combined organic layers were dried (Na_2_SO_4_), concentrated *in vacuo* and the resulting residue was purified by column chromatography on silica gel.

**General Procedure 2**

NaHCO_3_ (1.5 M, aq., 3 eq.) was added to a solution of the bromide (1.0 eq.) and the boronic acid (1.3 eq.) in DMF (3 mL/mmol) in a microwave vial and the vessel was degassed with argon. The requisite Pd catalyst (5 mol%) was then added, the microwave vial was sealed and irradiated at 150 °C for 30 min. The reaction mixture was allowed to cool, diluted with EtOAc (5 mL/mmol), washed with 0.5 M aq. LiCl (3 x 3 mL/mmol). The combined organic layers were dried (Na_2_SO_4_), then concentrated *in vacuo*. Purification by column chromatography on silica gel (solvents as stated) afforded the desired product.

**General Procedure 3**

The requisite bromide (600 mg, 2.79 mmol, 1.0 eq.), K_2_CO_3_ (1.16 g, 8.37 mmol, 3.0 eq.), the requisite boronic acid (572 mg, 3.07 mmol, 1.1 eq.), and Pd(dppf)Cl_2_ (100 mg, 0.140 mmol, 5 mol%) were added sequentially to a microwave vial equipped with a magnetic stirrer bar. The reaction vessel was fitted with a rubber septum and purged with N_2_ for 5 min, before addition of a degassed solution of 1,4-dioxane/water (5:1, 8 mL) *via* syringe. The vial was then sealed and the reaction heated to 100 °C for 18 h. The mixture was cooled down, diluted with EtOAc (30 mL), and washed with a 50/50 solution of water and brine (2 x 30 mL). The organic phase was dried (Na_2_SO_4_) and concentrated *in vacuo*. Purification by column chromatography on silica gel (solvents as stated) afforded the desired product.

**General Procedure 4** (Coppola et al., 2004)

To a solution of the requisite amide (1 eq.) in DMF (3 mL/mmol) was added the desired aryl bromide (1.3 eq.) followed by NaH (60% in oil, 1.3 eq.) portion-wise, and the resulting solution stirred for 18 h. The reaction was quenched with NH_4_Cl (sat. aq. sol.), extracted with EtOAc (3 x 1 mL/mmol), dried (Na_2_SO_4_), then concentrated *in vacuo*. The compound was purified by flash column chromatography (silica gel).

#### Experimental data

**3700 8-bromo-1-(4-methylbenzyl)-1,5-dihydrobenzo[e][1,4]oxazepin-2(3H)-one**

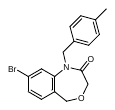

Following **general procedure 1**, (4-bromo-2-((4-methylbenzyl)amino)phenyl)methanol (704 mg, 2.29 mmol) (obtained from the corresponding methyl-2-amino-4-bromobenzoate and benzaldehyde) afforded the title compound **3700** as a white solid (482 mg, 61%), after purification on silica gel (EtOAc:pentane, 1:9); mp = 155-157 ºC (EtOAc); ^1^H NMR (500 MHz, CDCl_3_) δ 7.45 (d, *J* = 1.9 Hz, 1H), 7.36 (dd, *J* = 8.1, 1.8 Hz, 1H), 7.16 (d, *J* = 8.1 Hz, 1H), 7.13 (d, *J* = 8.2 Hz, 2H), 7.07 (d, *J* = 8.0 Hz, 2H), 5.04 (s, 2H), 4.45 (s, 2H), 4.04 (s, 2H), 2.29 (s, 3H); ^13^C NMR (126 MHz, CDCl_3_) δ 168.3, 143.9, 137.6, 133.7, 131.9, 129.8, 129.5, 128.5, 127.9, 124.8, 123.6, 77.4, 77.4, 77.2, 76.9, 67.6, 67.5, 50.6, 21.2; *m/z* LRMS (ESI^+^): 370 [M+Na]^+^; HRMS (ESI^+^): calc. for C_17_H_17_O_2_N^81^Br [M+H]^+^ 348.0417, found 348.0418.

**3702 8-bromo-3,4-dihydrobenzo[f][1,4]oxazepin-5(2H)-one 68** (Zablocki et al., 2016)

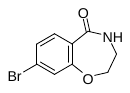

Sodium azide (86 mg, 1.32 mmol, 1.5 eq.) was added over 45 min to an ice-cold solution of 7-bromochroman-4-one (200 mg, 0.881 mmol) in methanesulfonic acid (3 mL) and the resulting solution stirred at RT for 16 h. Conc. HCl (1 mL) was added, and the resulting solid filtered and washed with water, to afford the title compound **3702** as a beige solid (140 mg, 66%); ^1^H NMR (500 MHz, CDCl_3_) δ 7.89 (d, *J* = 8.5 Hz, 1H), 7.25 (dd, *J* = 8.5, 1.9 Hz, 2H), 7.21 (d, *J* = 1.9 Hz, 1H), 6.56 (s, 1H), 4.43 – 4.38 (m, 2H), 3.52 (dd, *J* = 9.6, 5.0 Hz, 2H); ^13^C NMR (126 MHz, CDCl_3_) δ 169.0, 156.2, 133.8, 127.4, 126.1, 124.3, 121.8, 72.9, 41.9; ; *m/z* LRMS (ESI^+^): 243 [M+H]^+^; HRMS (EI): calc for C_9_H_8_^81^BrNO_2_ (M^+^) 242.9718 found 242.9707.

**3703 8-bromo-4-(4-methylbenzyl)-3,4-dihydrobenzo[f][1,4]oxazepin-5(2H)-one**

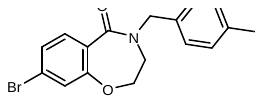

Following **general procedure 4**, amide **3702** (130 mg, 0.537 mmol) and 4-methylbenzyl bromide (199 mg, 1.07 mmol) afforded the title compound as a white solid (137 mg, 74%) after purification on silica gel (EtOAc:pentane, 1:9); mp = 64-66 °C (EtOAc); ^1^H NMR (500 MHz, CDCl_3_) δ 7.78 (d, *J* = 8.4 Hz, 1H), 7.30 (dd, *J* = 8.4, 1.9 Hz, 1H), 7.26 – 7.20 (m, 3H), 7.17 (s, 1H), 7.16 (d, *J* = 10.4 Hz, 2H), 4.77 (s, 2H), 4.16 (dd, *J* = 5.5, 4.6 Hz, 2H), 3.47 – 3.42 (m, 2H), 2.34 (s, 3H); ^13^C NMR (126 MHz, CDCl_3_) δ 167.8, 154.6, 137.8, 133.8, 132.9, 129.7, 128.5, 126.8, 126.4, 125.9, 124.6, 50.9, 45.9, 21.3; *m/z* LRMS (ESI^+^): 370 [M+Na]^+^; HRMS (ESI^+^): calc. for C_17_H_17_O_2_N^81^Br [M+H]^+^ 348.0417, found 348.0417.

**3706 6-bromo-1-(4-methylbenzyl)-1,5-dihydrobenzo[e][1,4]oxazepin-2(3H)-one**

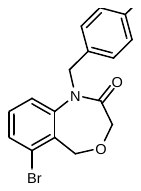

Following **general procedure 1**, (2-bromo-6-((4-methylbenzyl)amino)phenyl)methanol (510 mg, 1.67 mmol) (obtained from the corresponding methyl 2-amino-6-bromobenzoate and 4-methylbenzaldehyde) afforded the title compound **3706** as a white solid (393 mg, 68%) after purification on silica gel (EtOAc:pentane, 1:9); mp = 121 ºC (EtOAc); ^1^H NMR (500 MHz, CDCl_3_) δ 7.50 (dd, *J* = 6.5, 2.6 Hz, 1H), 7.24 (s, 1H), 7.26 – 7.21 (m, 1H), 7.12 (d, *J* = 8.1 Hz, 2H), 7.07 (d, *J* = 8.0 Hz, 2H), 5.06 (s, 2H), 4.74 (s, 2H), 4.04 (s, 2H), 2.30 (s, 3H); ^13^C NMR (126 MHz, CDCl_3_) δ 168.1, 144.1, 137.6, 133.7, 131.2, 130.7, 129.5, 129.4, 128.0, 125.3, 121.1, 77.4, 77.2, 76.9, 50.8, 21.2; *m/z* LRMS (ESI^+^): 370 [M(^81^Br)+Na]^+^; HRMS (ESI^+^): calc. for C_17_H_17_O_2_N^81^Br [M+H]^+^ 348.0417 found 348.0417.

**3827 8-bromo-1-(cyclohexylmethyl)-1,5-dihydrobenzo[e][1,4]oxazepin-2(3H)-one**

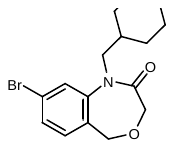

Following **general procedure 1**, ((4-bromo-2-((cyclohexylmethyl)amino)phenyl)methanol (310 mg, 0.863 mmol) (obtained from the corresponding methyl 2-amino-4-bromobenzoate and cyclohexane-carbaldehyde) afforded the title compound **3827** as an off white solid (262 mg, 90%) after purification on silica gel (EtOAc:pentane, 1:4); ^1^H NMR (400 MHz, CDCl_3_) δ 7.41 (d, *J* = 1.9 Hz, 1H), 7.39 (dd, *J* = 8.0, 1.8 Hz, 1H), 7.22 (d, *J* = 8.0 Hz, 1H), 4.62 (s, 2H), 3.95 (s, 2H), 3.77 (d, *J* = 6.8 Hz, 2H), 1.73 – 1.54 (m, 8H), 1.14 (td, *J* = 10.7, 9.0, 2.4 Hz, 3H), 1.01 – 0.87 (m, 2H); ^13^C NMR (101 MHz, CDCl_3_) δ 168.3, 144.3, 132.2, 129.5, 128.2, 124.4, 123.7, , 67.8, 67.7, 52.9, 36.8, 31.3, 26.3, 25.8; *m/z* LRMS (ESI^+^): 362 [M+Na]^+^; HRMS (ESI^+^): calc. for C_16_H_21_NO_2_Br [M+H]^+^ 338.07502, found 338.07519.

**3829 1-benzyl-7-bromo-1,5-dihydrobenzo[*e*][1,4]oxazepin-2(3*H*)-one**

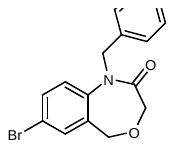

Following **general procedure 1**, (2-(benzylamino)-5-bromophenyl)methanol (940 mg, 3.22 mmol) (obtained from the corresponding methyl 2-amino-5-bromobenzoate and benzaldehyde) afforded the title compound **3829** as a white solid (707 mg, 66%), after purification on silica gel (EtOAc:pentane, 1:9); mp = 115-116 ºC (EtOAc); ^1^H NMR (500 MHz, MeOD) δ 7.61 (dd, *J* = 8.6, 2.3 Hz, 1H), 7.39 (d, *J* = 8.6 Hz, 1H), 7.30 – 7.18 (m, 5H), 5.14 (s, 3H), 4.44 (s, 2H), 3.99 (s, 2H); ^13^C NMR (126 MHz, MeOD) δ 170.3, 142.6, 138.2, 134.2, 134.2, 133.1, 129.7, 129.1, 128.8, 124.9, 120.8, 68.3, 68.1, 51.2, 49.5, 49.3, 49.2, 49.0, 48.8, 48.7, 48.5; *m/z* LRMS (ESI^+^): 356 [M(^81^Br)+Na]^+^354 ; HRMS (ESI^+^): calc. for C_16_H_15_O_2_N^81^Br [M+H]^+^ 334.0260, found 334.0261.

**4119 7-bromo-1-(cyclopropylmethyl)-1,5-dihydrobenzo[e][1,4]oxazepin-2(3H)-one**

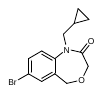

Following **general procedure 1**, (5-bromo-2-((cyclopropylmethyl)amino)phenyl)methanol (280 mg, 0.833 mmol) (obtained from the corresponding methyl 2-amino-5-bromobenzoate and cyclopropane-carbaldehyde) afforded the title compound **4119** as a white solid (262 mg, 90%), after purification on silica gel ((EtOAc:pentane, 1:4); ^1^H NMR (400 MHz, CDCl_3_) δ 7.57 (dd, *J* = 8.6, 2.4 Hz, 1H), 7.51 (d, *J* = 2.3 Hz, 1H), 7.18 (d, *J* = 8.6 Hz, 1H), 4.65 (s, 2H), 3.97 (s, 2H), 3.79 (d, *J* = 7.2 Hz, 2H), 1.10 – 0.95 (m, 1H), 0.49 – 0.37 (m, 2H), 0.28 – 0.19 (m, 2H); ^13^C NMR (101 MHz, CDCl_3_) δ 167.9, 141.9, 133.4, 133.1, 132.0, 123.5, 119.6, 67.6, 67.4, 51.6, 10.2, 4.0; *m/z* LRMS (ESI^+^): 296 [M+Na]^+^; HRMS (ESI^+^): calc. for C_13_H_15_NO_2_Br [M+H]^+^ 296.0286, found 296.0284.

**4118a 7-bromo-1-methyl-1,5-dihydrobenzo[e][1,4]oxazepin-2(3H)-one**

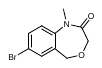

Following **general procedure 4**, bromide **3593** (75 mg, 0.310 mmol) afforded the title compound **4118a** as a white solid (262 mg, 90%), after purification on silica gel (EtOAc:pentane, 1:4); ^1^H NMR (400 MHz, CDCl_3_) δ 7.58 (dd, *J* = 8.6, 2.3 Hz, 1H), 7.50 (d, *J* = 2.3 Hz, 1H), 7.12 (d, *J* = 8.6 Hz, 1H), 4.60 (s, 2H), 4.02 (s, 2H), 3.41 (s, 3H); ^13^C NMR (126 MHz, CDCl_3_) δ 168.3, 142.9, 133.3, 133.2, 131.0, 122.4, 119.3, 67.8, 67.5, 34.8; ; *m/z* LRMS (ESI^+^): 255 [M+H]^+^.

**3698 6-bromo-2-(4-methylbenzyl)-3,4-dihydroisoquinolin-1(2H)-one**

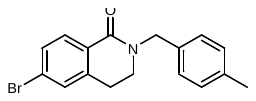

Following **general procedure 4**, 6-bromo-3,4-dihydro-isoquinoline (570 mg, 2.52 mmol) and 4-methylbenzyl bromide (873 mg, 3.28 mmol) afforded the title compound **3698** as a white solid (130 mg, 98%) after purification on silica gel (EtOAc:pentane, 1:4); mp = 97-99 °C (EtOAc); ^1^H NMR (400 MHz, CDCl_3_) δ 8.00 (d, *J* = 8.3 Hz, 1H), 7.33 (dd, *J* = 2.0, 1.0 Hz, 1H), 7.21 (d, *J* = 8.0 Hz, 2H), 7.14 (d, *J* = 7.9 Hz, 2H), 4.73 (s, 2H), 3.46 (dd, *J* = 7.1, 6.2 Hz, 2H), 2.90 (t, *J* = 6.6 Hz, 2H), 2.33 (s, 3H); ^13^C NMR (101 MHz, CDCl_3_) δ 163.9, 140.0, 137.3, 134.2, 130.4, 130.3, 129.9, 129.4, 128.5, 128.2, 128.2, 126.4, 50.2, 45.1, 27.9, 21.2; *m/z* LRMS (ESI^+^): 330 [M+H]^+^; HRMS (EI^+^): calc. for C_17_H_16_^81^BrNO (M^+^) requires 331.0395 found 331.0385.

**3145 1-(cyclohexylmethyl)-8-(pyridin-3-yl)-1,5-dihydrobenzo[e][1,4]oxazepin-2(3H)-one**

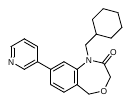

Following **general procedure 3**, bromide **3827** ((400 mg, 1.19 mmol) and 13-pyridyl boronic acid (190 mg, 1.54 mmol) afforded the title product **3145** (398 mg, 99%) as a yellow oil that solidified on standing after purification on silica gel (MeOH:CH_2_Cl_2_, 5%);^1^H NMR (400 MHz, CDCl_3_) δ 8.87 (s, 1H), 8.67 (s, 1H), 7.91 (dt, *J* = 7.9, 1.9 Hz, 1H), 7.52 – 7.41 (m, 3H), 4.73 (s, 2H), 4.02 (s, 2H), 3.88 (d, *J* = 6.9 Hz, 2H), 1.76 – 1.58 (m, 6H), 1.22 – 1.08 (m, 3H), 1.04 – 0.90 (m, 2H); ^13^C NMR (101 MHz, CDCl_3_) δ 168.5, 149.3, 148.3, 143.8, 140.0, 135.6, 134.5, 131.6, 129.1, 125.1, 123.8, 119.7, 67.9, 67.9, 53.0, 36.8, 31.3, 26.3, 25.7; *m/z* LRMS (ESI^+^): 337 [M+H]^+^; HRMS (ESI^+^): calc. for C_21_H_25_N_2_O_2_ [M+H]^+^ 337.19105, found 337.19085.

**3593 1,5-dihydrobenzo[e][1,4]oxazepin-2(3H)-one**

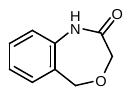

Following **general procedure 1**, title compound **3593** was obtained from *o*-aminobenzyl alcohol (1.00 g, 8.12 mmol) as an orange solid (621 mg, 38%), after purification on silica gel (EtOAc:pentane, 2:3); mp = 98-99 °C (EtOAc); ^1^H NMR (500 MHz, CDCl_3_) δ 9.65 (s, 1H), 8.09 (dd, *J* = 8.1, 1.2 Hz, 1H), 7.23 (dd, *J* = 7.6, 1.6 Hz, 1H), 7.15 (td, *J* = 7.5, 1.2 Hz, 1H), 4.76 (s, 2H); ^13^C NMR (126 MHz, CDCl_3_) δ 164.9, 136.7, 130.0, 129.4, 129.0, 125.2, 122.5, 64.4, 43.1; *m/z* LRMS (ESI^+^): 164 [M+H]^+^; HRMS (ESI^+^): calc. for C_9_H_10_NO_2_ [M+H]^+^ 164.0701, found 164.0706.

**3594 1-benzyl-1,5-dihydrobenzo[e][1,4]oxazepin-2(3H)-one**

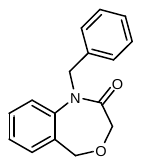

Following **general procedure 4**, amide **3593** (100 mg, 0.612 mmol) and benzyl bromide (145 µL, 1.23 mmol) afforded the title compound as a white solid (43 mg, 28%) after purification on silica gel (EtOAc:pentane, 1:4); mp = 90-92 °C (EtOAc); ^1^H NMR (500 MHz, CDCl_3_) δ 7.40 (ddd, *J* = 8.1, 7.3, 1.7 Hz, 1H), 7.26 (s, 0H), 5.12 (s, 2H), 4.52 (s, 2H), 4.06 (s, 2H); ^13^C NMR (126 MHz, CDCl_3_) δ 168.6, 142.7, 137.2, 130.7, 130.2, 129.6, 128.8, 127.9, 127.7, 126.8, 121.4, 68.1, 67.7, 50.9; *m/z* LRMS (ESI^+^): 276 [M+Na]^+^; HRMS (ESI^+^): calc. for C_16_H_16_NO_2_ [M+H]^+^ 254.1175, found 254.1176.

**3596 1-(4-methylbenzyl)-7-phenyl-1,5-dihydrobenzo[e][1,4]oxazepin-2(3H)-one**

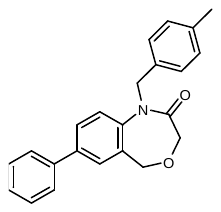

Following **general procedure 2** using Pd(PPh_3_)_4_, bromide **3146** (Partridge et al., 2017) (50 mg, 0.144 mmol), phenylboronic acid (23 mg, 0.187 mmol) afforded the title compound **3596** as yellow solid (14 mg, 28%) after purification on silica gel (EtOAc:pentane, 1:9); mp = 102-105 °C (EtOAc); ^1^H NMR (500 MHz, CDCl_3_) δ 7.61 (dd, *J* = 8.4, 2.2 Hz, 1H), 7.59 – 7.53 (m, 2H), 7.51 (d, *J* = 2.2 Hz, 1H), 7.44 (dd, *J* = 8.4, 6.9 Hz, 2H), 7.37 (d, *J* = 7.4 Hz, 1H), 7.34 (d, *J* = 8.3 Hz, 1H), 7.08 (d, *J* = 7.9 Hz, 2H), 5.11 (s, 2H), 4.58 (s, 2H), 4.11 (s, 2H), 2.30 (s, 3H); ^13^C NMR (126 MHz, CDCl_3_) δ 168.6, 141.8, 139.7, 139.7, 137.4, 134.2, 129.9, 129.5, 129.2, 129.1, 128.7, 127.9, 127.9, 127.0, 121.8, 77.4, 77.2, 76.9, 68.3, 67.8, 50.6, 21.2; *m/z* LRMS (ESI^+^): 344 [M+H]^+^; HRMS (ESI^+^): calc. for C_23_H_22_NO_2_ [M+H]^+^ 344.1645, found 344.1643.

**3599 7-(4-methoxyphenyl)-1-(4-methylbenzyl)-1,5-dihydrobenzo[e][1,4]oxazepin-2(3H)-one**

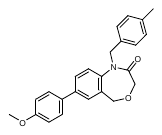

Following **general procedure 2** using Pd(PPh_3_)_4_, bromide **3146** (50 mg, 0.144 mmol) and 4-methoxyphenylboronic acid (29 mg, 0.188 mmol) afforded the title compound **3599** as a yellow solid (11 mg, 21%) after purification on silica gel ((EtOAc:pentane, 1:9); mp = 48-50 °C (EtOAc); ^1^H NMR (500 MHz, CDCl_3_) δ 7.56 (dd, *J* = 8.4, 2.2 Hz, 1H), 7.53 – 7.47 (m, 2H), 7.46 (d, *J* = 2.2 Hz, 1H), 7.31 (d, *J* = 8.4 Hz, 1H), 7.16 (d, *J* = 8.1 Hz, 2H), 7.08 (d, *J* = 7.9 Hz, 2H), 7.02 – 6.94 (m, 2H), 5.10 (s, 2H), 4.56 (s, 2H), 4.10 (s, 2H), 3.85 (s, 3H), 2.29 (s, 3H); ^13^C NMR (126 MHz, CDCl_3_) δ 168.6, 159.6, 141.1, 139.3, 137.4, 134.2, 132.2, 129.9, 129.5, 128.7, 128.2, 128.1, 127.9, 121.8, 114.5, 77.4, 77.2, 76.9, 68.4, 67.8, 55.5, 50.6, 21.2; *m/z* LRMS (ESI^+^): 396 [M+Na]^+^; HRMS (ESI^+^): calc. for C_24_H_24_NO_3_ [M+H]^+^ 374.1750, found 374.1749.

**3600 7-(4-fluorophenyl)-1-(4-methylbenzyl)-1,5-dihydrobenzo[e][1,4]oxazepin-2(3H)-one**

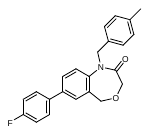

Following **general procedure 2** using Pd(PPh_3_)_4_, bromide **3146** (50 mg, 0.144 mmol) and 4-fluorophenylboronic acid (26 mg, 0.188 mmol) ) afforded the title compound **3600** as a yellow solid (14 mg, 27%) after purification by flash column chromatography (EtOAc:pentane, 15%); mp = 127-129 °C (EtOAc); ^1^H NMR (500 MHz, CDCl_3_) δ 7.56 (dd, *J* = 8.4, 2.3 Hz, 1H), 7.56 – 7.48 (m, 2H), 7.46 (d, *J* = 2.2 Hz, 1H), 7.34 (s, 1H), 7.16 (d, *J* = 8.1 Hz, 2H), 7.13 (t, *J* = 8.7 Hz, 2H), 7.08 (d, *J* = 7.9 Hz, 2H), 5.10 (s, 2H), 4.57 (s, 2H), 4.10 (s, 2H), 2.30 (s, 3H); ^13^C NMR (126 MHz, CDCl_3_) δ 168.6, 163.8, 161.8, 141.7, 138.7, 137.4, 134.1, 130.0, 129.5, 129.1, 128.7, 128.6, 128.5, 127.9, 121.9, 116.1, 115.9, 77.4, 77.2, 76.9, 68.3, 67.8, 50.6, 21.2; *m/z* LRMS (ESI^+^): 385 [M+Na]^+^; HRMS (ESI^+^): calc. for C_23_H_21_FNO_2_ [M+H]^+^ 362.1550, found 362.1546.

**3601 1-(4-methylbenzyl)-7-(p-tolyl)-1,5-dihydrobenzo[e][1,4]oxazepin-2(3H)-one**

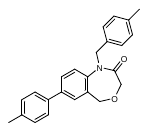

Following **general procedure 2** using Pd(PPh_3_)_4_, bromide **3146** (50 mg, 0.144 mmol) and 4-tolylboronic acid (26 mg, 0.188 mmol) afforded the title compound **3601** as a pale yellow solid (167 mg, 32%) after purification on silica gel (EtOAc:pentane, 15%); mp = 100-101 °C (EtOAc); ^1^H NMR (500 MHz, CDCl_3_) δ 7.49 (d, *J* = 2.2 Hz, 1H), 7.46 (d, *J* = 8.2 Hz, 2H), 7.32 (d, *J* = 8.4 Hz, 1H), 7.26 (s, 6H), 7.19 – 7.14 (m, 2H), 7.08 (d, *J* = 7.9 Hz, 2H), 5.10 (s, 2H), 4.57 (s, 2H), 4.10 (s, 2H), 2.39 (s, 3H), 2.30 (s, 3H); ^13^C NMR (126 MHz, CDCl_3_) δ 168.6, 141.5, 139.6, 137.8, 137.4, 136.8, 134.2, 129.9, 129.8, 129.5, 128.9, 128.4, 127.9, 126.9, 121.7, 68.4, 67.8, 50.6, 21.3, 21.2; *m/z* LRMS (ESI^+^): 380 [M+Na]^+^; HRMS (ESI^+^): calc. for C_24_H_24_NO_2_ [M+H]^+^ 358.1801, found 358.1803.

**3697 methyl 4-(1-(4-methylbenzyl)-2-oxo-1,2,3,5-tetrahydrobenzo[e][1,4]oxazepin-7-yl)benzoate**

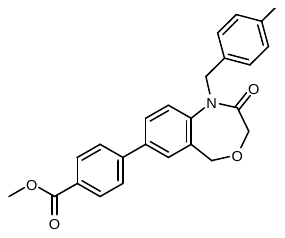

Following **general procedure 2** using Pd(dppf)Cl_2_, bromide **3146** (200 mg, 0.578 mmol) and 4-methyoxycarbonyl-phenylboronic acid (135 mg, 0.751 mmol), afforded the title compound **3697** as a pale yellow solid (51 mg, 22%) after purification on silica gel (EtOAc:pentane, 3:7); mp = 138-140 ºC (EtOAc); ^1^H NMR (500 MHz, CDCl_3_) δ 8.11 (d, *J* = 8.4 Hz, 2H), 7.66 (d, *J* = 2.2 Hz, 1H), 7.63 (d, *J* = 8.6 Hz, 2H), 7.37 (d, *J* = 8.4 Hz, 1H), 7.19 – 7.14 (m, 2H), 7.08 (d, *J* = 7.9 Hz, 2H), 5.12 (s, 2H), 4.59 (s, 2H), 4.11 (s, 2H), 3.94 (s, 3H), 2.30 (s, 3H); ^13^C NMR (126 MHz, CDCl_3_) δ 168.8, 167.2, 144.3, 142.8, 138.6, 137.7, 134.3, 130.7, 130.4, 129.8, 129.7, 129.7, 129.1, 128.1, 127.2, 122.2, 52.6, 50.8, 21.5.; *m/z* LRMS (ESI^+^) 424 [M+Na]^+^; HRMS (ESI^+^): calc. for C_25_H_24_NO_4_ [M+H]^+^ 402.1700, found 402.1701.

**3701 1-(4-methylbenzyl)-8-(pyridin-4-yl)-1,5-dihydrobenzo[e][1,4]oxazepin-2(3H)-one**

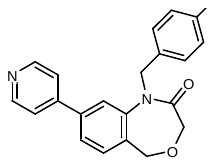

Following **general procedure 2** using Pd(dppf)Cl_2_, bromide **3700** (201 mg, 0.578 mmol) and 4-pyridinylboronic acid (106 mg, 0.867 mmol) afforded the title compound **3701** as a brown solid (44 mg, 24%) after purification on silica gel (MeOH:CH_2_Cl_2_; 2%); mp = 108-110 ºC (MeOH); ^1^H NMR (500 MHz, CDCl_3_) δ 8.69 (d, *J* = 6.1 Hz, 1H), 7.50 (d, *J* = 1.7 Hz, 1H), 7.49 (dd, *J* = 7.7, 1.7 Hz, 1H), 7.45 – 7.39 (m, 3H), 7.17 (d, *J* = 7.9 Hz, 2H), 7.08 (d, *J* = 7.8 Hz, 2H), 5.14 (s, 2H), 4.57 (s, 2H), 4.10 (s, 2H), 2.30 (s, 3H); ^13^C NMR (126 MHz, CDCl_3_) δ 168.5, 150.7, 147.1, 143.5, 140.3, 137.6, 134.0, 131.5, 130.2, 129.6, 128.0, 125.3, 121.7, 120.1, 67.8, 67.8, 50.7, 21.3.; *m/z* LRMS (ESI^+^) 345 [M+H]^+^; HRMS (ESI^+^): calc. for C_22_H_21_N_2_O_2_ [M+H]^+^ 345.1578, found 345.1597

**3704 4-(4-methylbenzyl)-8-(pyridin-4-yl)-3,4-dihydrobenzo[f][1,4]oxazepin-5(2H)-one**

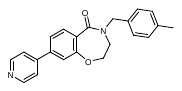

Following **general procedure 2** using Pd(dppf)Cl_2_, bromide **3703** (40 mg, 115 mmol) and 4-pyridinylboronic acid (18 mg, 0.150 mmol) afforded the title compound **3704** as a brown solid (37 mg, 93%) after purification on silica gel (MeOH:CH_2_Cl_2_; 2%); mp = 127-129 ºC (MeOH); ^1^H NMR (500 MHz, CDCl_3_) δ 8.72 (s, 1H), 7.57 (s, 1H), 7.45 (dd, *J* = 8.1, 1.8 Hz, 1H), 7.29 – 7.26 (m, 1H), 7.26 (d, *J* = 3.4 Hz, 1H), 7.17 (d, *J* = 7.8 Hz, 1H), 4.81 (s, 1H), 4.22 (t, *J* = 5.1 Hz, 1H), 3.50 (t, *J* = 5.0 Hz, 1H), 2.35 (s, 2H); ^13^C NMR (126 MHz, CDCl_3_) δ 168.0, 154.6, 149.6, 142.4, 137.8, 133.8, 132.7, 129.7, 128.5, 127.5, 122.0, 120.1, 73.5, 50.9, 46.0, 21.3; *m/z* LRMS (ESI^+^) 345 [M+H]^+^; HRMS (ESI^+^): calc. for C_22_H_21_N_2_O_2_ [M+H]^+^ 345.1578, found 345.1596.

**3705** **1-(4-methylbenzyl)-7-(pyrimidin-5-yl)-1,5-dihydrobenzo[*e*][1,4]oxazepin-2(3*H*)-one**

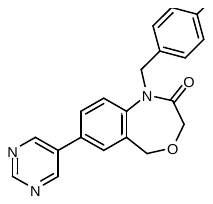

Following **general procedure 2** using Pd(dppf)Cl_2_, bromide **3146** (50 mg, 0.144 mmol) and pyrimidine-5-boronic acid (23 mg, 0.188 mmol) afforded the title compound **3705** as a pale yellow solid (15 mg, 31%) after purification on silica gel (EtOAc:pentane, 2:3); mp = 192-194 °C (EtOAc); ^1^H NMR (500 MHz, CDCl_3_) δ 9.22 (s, 1H), 8.94 (s, 2H), 7.52 (d, *J* = 2.2 Hz, 1H), 7.44 (d, *J* = 8.3 Hz, 1H), 7.16 (d, *J* = 8.1 Hz, 3H), 7.09 (d, *J* = 7.9 Hz, 2H), 5.13 (s, 2H), 4.60 (s, 2H), 4.12 (s, 2H), 2.30 (s, 3H); ^13^C NMR (126 MHz, CDCl_3_) δ 168.4, 158.0, 154.9, 143.4, 137.6, 133.9, 133.1, 132.6, 130.7, 129.6, 129.2, 128.6, 127.9, 122.5, 68.1, 67.9, 50.5, 21.3; *m/z* LRMS (ESI^+^): 346 [M+H]^+^; HRMS (ESI^+^): calc. for C_21_H_20_N_3_O_2_ [M+H]^+^ 346.1550 found 346.1547.

**3707 1-(4-methylbenzyl)-6-(pyridin-4-yl)-1,5-dihydrobenzo[e][1,4]oxazepin-2(3H)-one**

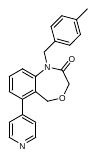

Following **general procedure 2** using Pd(dppf)Cl_2_, bromide **3706** (60 mg, 0.173 mmol) and 4-pyridinylboronic acid (32 mg, 0.260 mmol) afforded the title compound **3707** as an orange solid (55 mg, 92%) after purification on silica gel (MeOH:CH_2_Cl_2_; 2%); mp = 134-136 ºC (MeOH); ^1^H NMR (500 MHz, CDCl_3_) δ 8.68 (s, 2H), 7.47 (t, *J* = 7.9 Hz, 1H), 7.39 – 7.33 (m, 3H), 7.26 (dd, *J* = 7.6, 1.2 Hz, 1H), 7.07 (d, *J* = 7.8 Hz, 2H), 5.13 (s, 2H), 4.40 (s, 2H), 4.20 (s, 2H), 2.29 (s, 3H); ^13^C NMR (126 MHz, CDCl_3_) δ 168.2, 150.1, 147.3, 144.0, 141.2, 137.5, 133.9, 129.9, 129.5, 127.9, 127.7, 127.0, 124.3, 121.5, 67.9, 64.4, 50.8, 21.2; *m/z* LRMS (ESI^+^) 345 [M+H]^+^; HRMS (ESI^+^): calc. for C_22_H_21_N_2_O_2_ [M+H]^+^ requires 345.1598 found 345.1597.

**3710 7-(benzylamino)-1-(4-methylbenzyl)-1,5-dihydrobenzo[*e*][1,4]oxazepin-2(3*H*)-one**

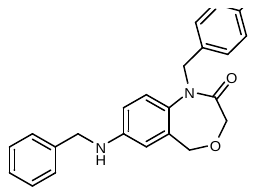

Bromide **3146** (50 mg, 0.144 mmol), benzylamine (19 µL, 0.173 mmol), NaOtBu (28 mg, 0.289 mmol), XPhos (7 mg, 0.014 mmol) and Pd(OAc)_2_ (2 mg, 0.007 mmol) were added sequentially to a vial and degassed before addition of degassed 1,4-dioxane (3 mL). The vial was sealed and heated to 110 ºC for 16 h. The reaction was cooled down to RT and filtered through celite, using EtOAc as an eluent, the filtrate was then concentrated *in vacuo*. Purification on silica gel (EtOAc:pentane, 3:7) afforded the title compound **3710** as a yellow oil (42 mg, 78%); ^1^H NMR (500 MHz, CDCl_3_) δ 7.35 (d, *J* = 3.8 Hz, 4H), 7.32 – 7.27 (m, 1H), 7.13 (d, *J* = 8.0 Hz, 2H), 7.05 (dd, *J* = 8.3, 3.1 Hz, 3H), 6.62 (dd, *J* = 8.7, 2.8 Hz, 1H), 6.51 (d, *J* = 2.7 Hz, 1H), 4.98 (s, 2H), 4.38 (s, 2H), 4.31 (s, 2H), 4.03 (s, 2H), 2.29 (s, 3H); ^13^C NMR (126 MHz, CDCl_3_) δ 168.5, 146.8, 138.9, 137.2, 134.5, 132.5, 130.7, 129.3, 128.9, 128.1, 127.6, 127.6, 122.7, 114.0, 113.7, 50.6, 48.5, 21.2; *m/z* LRMS (ESI^+^) 373 [M+H]^+^; HRMS (ESI^+^): calc. for C_24_H_25_N_2_O_2_ [M+H]^+^ requires 373.1911, found 373.1907.

**3824 1-(4-methylbenzyl)-7-(phenylamino)-1,5-dihydrobenzo[e][1,4]oxazepin-2(3H)-one**

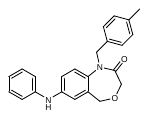

Bromide **3146** (101 mg, 0.289 mmol), aniline (32 µL, 0.347 mmol), NaO*^t^*Bu (56 mg, 0.578 mmol), XPhos (14 mg, 0.029 mmol) and Pd(OAc)_2_ (3 mg, 0.014 mmol) were added sequentially to a vial and degassed before addition of degassed 1,4-dioxane (3 mL). The vial was sealed and heated to 110 ºC for 16 h. The reaction was cooled down to RT and filtered through celite, using EtOAc as an eluent, the filtrate was then concentrated *in vacuo*. Purification on silica gel (acetone:pentane, 3:7) afforded the title compound **3824** as a brown solid (41 mg, 40%) ; mp= 122-124 ºC (CH_2_Cl_2_); ^1^H NMR (500 MHz, CDCl_3_) δ 7.29 (dd, *J* = 8.5, 7.3 Hz, 2H), 7.15 (dd, *J* = 8.4, 2.1 Hz, 3H), 7.10 – 7.02 (m, 5H), 7.01 – 6.95 (m, 1H), 6.95 (d, *J* = 2.7 Hz, 1H), 5.02 (s, 2H), 4.42 (s, 2H), 4.07 (s, 2H), 2.29 (s, 3H); ^13^C NMR (126 MHz, CDCl_3_) δ 168.5, 142.2, 142.2, 137.3, 135.1, 134.4, 130.8, 129.6, 129.4, 128.0, 122.7, 122.1, 118.8, 118.3, 118.2, 77.4, 77.2, 76.9, 50.6, 21.2; *m/z* LRMS (ESI^+^) 359 [M+H]^+^; HRMS (ESI^+^): calc. for C_23_H_23_N_2_O_2_ [M+H]^+^ requires 359.1754, found 359.2904.

**3825 1-benzyl-7-(pyridin-4-yl)-1,5-dihydrobenzo[e][1,4]oxazepin-2(3H)-one**

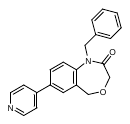

Following **general procedure 2** using Pd(dppf)Cl_2_, bromide **3829** (300 mg, 0.908 mmol) and 4-pyridinylboronic acid (144 mg, 1.17 mmol) afforded the title compound **3825** as a brown solid (96 mg, 32%) after purification on silica gel (MeOH:CH_2_Cl_2_, 2%); mp = 129-130 ºC (MeOH); ^1^H NMR (500 MHz, MeOD) δ 8.59 (d, *J* = 5.4 Hz, 1H), 7.89 (dd, *J* = 8.4, 2.3 Hz, 1H), 7.80 (d, *J* = 2.3 Hz, 1H), 7.76 – 7.71 (m, 2H), 7.62 (d, *J* = 8.5 Hz, 1H), 7.32 – 7.23 (m, 3H), 5.22 (s, 1H), 4.59 (s, 2H), 4.05 (s, 1H); ^13^C NMR (126 MHz, MeOD) δ 170.5, 150.7, 149.1, 144.5, 138.3, 137.3, 131.8, 130.1, 129.8, 129.7, 129.0, 128.7, 123.7, 122.0, 68.7, 68.4; *m/z* LRMS (ESI^+^) 331 [M+H]^+^; HRMS (ESI^+^): calc. for C_21_H_19_N_2_O_2_+H [M+H]^+^ requires 331.1441, found 331.1439.

**4115 1-(4-methylbenzyl)-7-(pyridin-2-yl)-1,5-dihydrobenzo[e][1,4]oxazepin-2(3H)-one**

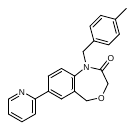

Bromide **3146** (110 mg, 0.328 mmol), 2-pyridyl-MIDA boronate (116 mg, 0.492 mmol), Cu(OAc)_2_ (33 mg, 0.164 mmol), K_2_CO_3_ (227 mg, 1.64 mmol), XPhos (10 mg, 0.020 mmol) and Pd_2_(dba)_3_ (6 mg, 0.066 mmol) were added sequentially to a vial and degassed before addition of degassed DMF. The resulting suspension was degassed further for 5 min, the vial was vealed and heated to 100 °C for 16 h. The reaction was cooled down before addition of EtOAc (10 mL) and H_2_O:brine (1:1, 30 mL). The organic phase was washed further with H_2_O:brine (1:1, 2 x 30 mL), dried (Na_2_SO_4_), filtered and concentrated *in vacuo*. The residue was purified on silica gel (MeOH:CH_2_Cl_2_, 3%) to afford the title product **4115** (113 mg, 99%) as a beige solid; ^1^H NMR (400 MHz, CDCl_3_) δ 8.67 (ddd, *J* = 4.8, 1.8, 1.0 Hz, 1H), 8.01 (dd, *J* = 8.4, 2.2 Hz, 1H), 7.94 (d, *J* = 2.1 Hz, 1H), 7.79 – 7.70 (m, 1H), 7.69 (dt, *J* = 8.0, 1.2 Hz, 1H), 7.38 (d, *J* = 8.4 Hz, 1H), 7.23 (ddd, *J* = 7.3, 4.8, 1.3 Hz, 1H), 7.14 (d, *J* = 8.1 Hz, 2H), 7.04 (d, *J* = 7.9 Hz, 2H), 5.11 (s, 2H), 4.58 (s, 2H), 4.09 (s, 2H), 2.26 (s, 3H); ^13^C NMR (101 MHz, CDCl_3_) δ 168.5, 155.9, 149.9, 143.0, 137.7, 137.3, 137.0, 134.0, 129.9, 129.4, 129.1, 128.4, 127.9, 122.5, 121.7, 120.4, 68.2, 67.7, 50.3; ; *m/z* LRMS (ESI^+^) 345 [M+H]^+^; HRMS (ESI^+^): calc. for C_22_H_21_N_2_O_2_ [M+H]^+^ requires 345.1598 found 345.1599.

**4116 7-(2-methoxypyridin-4-yl)-1-(4-methylbenzyl)-1,5-dihydrobenzo[e][1,4]oxazepin-2(3H)-one**

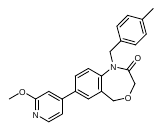

Following **general procedure 3**, **3146** (110 mg, 0.328 mmol) and 2-methoxypyridine boronic acid (55 mg, 0.36 mmol) afforded the title product **4116** (119 mg, 97%) as a light yellow oil that solidified on standing after purification on silica gel (EtOAc:pentane, 2:3); ^1^H NMR (400 MHz, CDCl_3_) δ 8.21 (dd, *J* = 5.4, 0.7 Hz, 1H), 7.63 (dd, *J* = 8.4, 2.2 Hz, 1H), 7.54 (d, *J* = 2.2 Hz, 1H), 7.37 (d, *J* = 8.4 Hz, 1H), 7.15 (d, *J* = 8.1 Hz, 2H), 7.08 (d, *J* = 7.8 Hz, 2H), 7.07 (dd, *J* = 5.5, 1.6 Hz, 1H), 6.91 (dd, *J* = 1.6, 0.7 Hz, 1H), 5.11 (s, 2H), 4.58 (s, 2H), 4.10 (s, 2H), 3.98 (s, 3H), 2.29 (s, 3H); ^13^C NMR (101 MHz, CDCl_3_) δ 168.5, 165.1, 149.7, 147.5, 143.3, 137.5, 136.5, 134.0, 130.2, 129.5, 129.2, 128.6, 127.8, 122.0, 115.1, 108.4, 68.2, 67.9, 53.8, 50.5, 21.2; *m/z* LRMS (ESI^+^) 375 [M+H]^+^; HRMS (ESI^+^): calc. for C_23_H_23_N_2_O_2_ [M+H]^+^ requires 375.1709 found 375.1709.

**4117 7-(2-aminopyridin-4-yl)-1-(4-methylbenzyl)-1,5-dihydrobenzo[e][1,4]oxazepin-2(3H)-one**

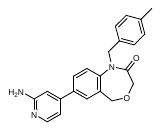

Following **general procedure 3**, **3146** (110 mg, 0.328 mmol) and 2-aminopyridine-4-boronic pinacol ester (79 mg, 0.361 mmol) afforded the title product **4237** (113 mg, 96%) as a light yellow oil that solidified on standing after purification on silica gel (MeOH:CH_2_Cl_2_, 4%); ^1^H NMR (400 MHz, MeOD) δ 7.92 (dd, *J* = 5.5, 0.8 Hz, 1H), 7.71 (dd, *J* = 8.4, 2.2 Hz, 1H), 7.60 (d, *J* = 2.2 Hz, 1H), 7.52 (d, *J* = 8.4 Hz, 1H), 7.13 (d, *J* = 8.2 Hz, 2H), 7.04 (dd, *J* = 8.4, 0.8 Hz, 2H), 6.83 (dd, *J* = 5.5, 1.7 Hz, 1H), 6.80 (dd, *J* = 1.6, 0.8 Hz, 1H), 5.47 (s, 2H), 5.12 (s, 2H), 4.86 (s, 4H), 4.51 (s, 2H), 3.99 (s, 2H), 2.23 (s, 3H); ^13^C NMR (101 MHz, MeOD) δ 170.4, 161.4, 150.2, 148.6, 143.9, 138.6, 138.4, 135.3, 131.5, 130.2, 130.2, 129.8, 129.5, 129.0, 123.3, 112.1, 107.4, 68.7, 68.4, 50.8, 25.0; *m/z* LRMS (ESI^+^) 360 [M+H]^+^; HRMS (ESI^+^): calc. for C_22_H_22_N_3_O_2_ [M+H]^+^ requires 360.1712 found 360.1711.

**4118 1-methyl-7-(pyridin-4-yl)-1,5-dihydrobenzo[e][1,4]oxazepin-2(3H)-one**

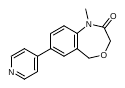

Following **general procedure 3**, 7-bromo-1-methyl-1,5-dihydrobenzo[e][1,4]oxazepin-3(2H)-one (30 mg, 0.118 mmol) and 4-pyridyl boronic acid hydrate (16 mg, 0.129 mmol) afforded the title product **4237** (22 mg, 74%) as a beige solid after purification on silica gel (MeOH:CH_2_Cl_2_, 2%); ^1^H NMR (400 MHz, CDCl_3_) δ 8.72 – 8.66 (m, 2H), 7.74 (dd, *J* = 8.4, 2.2 Hz, 1H), 7.63 (d, *J* = 2.2 Hz, 1H), 7.56 – 7.46 (m, 2H), 7.36 (d, *J* = 8.4 Hz, 1H), 4.73 (s, 2H), 4.08 (s, 2H), 3.48 (s, 3H) ; ^13^C NMR (126 MHz, CDCl_3_) δ 168.5, 150.5, 147.0, 144.6, 136.1, 129.8, 129.1, 128.7, 121.5, 121.4, 68.1, 67.8, 34.8; *m/z* LRMS (ESI^+^) 255 [M+H]^+^; HRMS (ESI^+^): calc. for C_15_H_15_N_2_O_2_ [M+H]^+^ requires 255.1134 found 255.1135.

**4120 1-(cyclopropylmethyl)-7-(pyridin-4-yl)-1,5-dihydrobenzo[e][1,4]oxazepin-2(3H)-one**

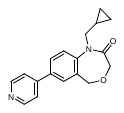

Following **general procedure 3**, **4119** (134 mg, 0.453 mmol) and 4-pyridyl boronic acid hydrate (61 mg, 0.498 mmol) afforded the title product **4120** (130 mg, 97%) as an orange oil that solidified on standing after purification on silica gel (MeOH:CH_2_Cl_2_, 4%); ^1^H NMR (500 MHz, DMSO) δ 8.69 (s, 2H), 8.01 (d, *J* = 2.2 Hz, 1H), 7.98 (dd, *J* = 8.4, 2.3 Hz, 1H), 7.79 (d, *J* = 4.6 Hz, 2H), 7.66 (d, *J* = 8.4 Hz, 1H), 4.72 (s, 2H), 3.86 (s, 2H), 3.84 (d, *J* = 7.2 Hz, 2H), 0.98 (tt, *J* = 7.6, 4.7 Hz, 1H), 0.42 – 0.32 (m, 2H), 0.22 – 0.15 (m, 2H); ^13^C NMR (126 MHz, DMSO) δ 167.3, 150.3, 145.6, 143.3, 134.8, 130.1, 128.7, 128.4, 122.7, 67.0, 66.8, 50.1, 10.0, 3.5; *m/z* LRMS (ESI^+^) 295 [M+H]^+^; HRMS (ESI^+^): calc. for C_18_H_19_N_2_O_2_ [M+H]^+^ requires 295.1447 found 295.1448.

**4122 1-(4-methylbenzyl)-7-(thiazol-5-yl)-1,5-dihydrobenzo[e][1,4]oxazepin-2(3H)-one**

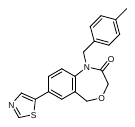

Following **general procedure 3**, **3146** (100 mg, 0.290 mmol) and thiazole-5-boronic pinacol ester (73 mg, 0.348 mmol) afforded the title product **4122** (98 mg, 97%) as an off white solid purification on silica gel (MeOH:CH_2_Cl_2_, 4%); ^1^H NMR (400 MHz, MeOD) δ 8.96 (s, 1H), 8.18 (s, 1H), 7.74 (dd, *J* = 8.4, 2.3 Hz, 1H), 7.65 (d, *J* = 2.3 Hz, 1H), 7.53 (d, *J* = 8.5 Hz, 1H), 7.15 (d, *J* = 8.0 Hz, 2H), 7.06 (d, *J* = 8.2 Hz, 2H), 5.13 (s, 2H), 4.51 (s, 2H), 4.01 (s, 2H), 2.24 (s, 3H); ^13^C NMR (101 MHz, MeOD) δ 170.4, 154.9, 143.4, 140.2, 139.6, 138.6, 135.3, 131.9, 130.9, 130.3, 129.6, 129.5, 129.1, 129.1, 123.9, 68.5, 68.4, 25.0; *m/z* LRMS (ESI^+^) 351 [M+H]^+^; HRMS (ESI^+^): calc. for C_20_H_19_N_2_O_2_S [M+H]^+^ requires 351.1167 found 351.1166

**4123 7-(1-methyl-1H-pyrazol-5-yl)-1-(4-methylbenzyl)-1,5-dihydrobenzo[e][1,4]oxazepin-2(3H)-one**

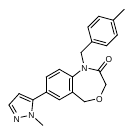

Following **general procedure 3**, **XXX** (100 mg, 0.290 mmol) and 1-methylpyrazole-5- boronic acid (72 mg, 0.348 mmol) afforded the title product **4237** (96 mg, 95%) as a light orange oil that solidified on standing after purification on silica gel (MeOH:CH_2_Cl_2_, 4%); ^1^H NMR (400 MHz, MeOD) δ 7.57 (d, *J* = 1.3 Hz, 2H), 7.47 (d, *J* = 2.0 Hz, 1H), 7.46 (t, *J* = 1.3 Hz, 1H), 7.15 (d, *J* = 8.1 Hz, 2H), 7.05 (d, *J* = 7.4 Hz, 1H), 6.36 (d, *J* = 2.0 Hz, 1H), 5.14 (s, 1H), 4.88 (s, 1H), 4.52 (s, 2H), 4.01 (s, 2H), 3.84 (s, 3H), 2.23 (s, 3H); ^13^C NMR (101 MHz, MeOD) δ 170.4, 143.9, 143.6, 139.4, 138.6, 135.3, 131.6, 131.4, 130.3, 130.2, 129.0, 123.4, 107.4, 68.5, 68.4, 37.7, 21.1; *m/z* LRMS (ESI^+^) 348 [M+H]^+^; HRMS (ESI^+^): calc. for C_21_H_22_N_3_O_2_ [M+H]^+^ requires 348.1712 found 348.1713

**3699 2-(4-methylbenzyl)-6-(pyridin-4-yl)-3,4-dihydroisoquinolin-1(2H)-one**

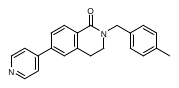

Following **general procedure 2** using Pd(dppf)Cl_2_, compound **3698** (50 mg, 0.151 mmol) and 4-pyridinylboronic acid (24 mg, 0.197 mmol), afforded the title compound as a brown solid (48 mg, 96%) after purification by flash column chromatography (MeOH:CH_2_Cl_2_, 5%); mp = 151-152 ºC (MeOH); ^1^H NMR (500 MHz, CDCl_3_) δ 8.69 (d, *J* = 4.7 Hz, 2H), 8.26 (d, *J* = 8.0 Hz, 1H), 7.62 (dd, *J* = 8.1, 1.9 Hz, 1H), 7.52 (d, *J* = 6.1 Hz, 1H), 7.42 (d, *J* = 1.8 Hz, 1H), 7.24 (d, *J* = 8.1 Hz, 2H), 7.15 (d, *J* = 7.9 Hz, 2H), 4.78 (s, 2H), 3.52 (t, *J* = 6.6 Hz, 2H), 3.01 (t, *J* = 6.6 Hz, 2H), 2.34 (s, 3H); ^13^C NMR (126 MHz, CDCl_3_) δ 164.1, 150.5, 147.5, 141.4, 139.0, 137.4, 134.3, 130.1, 129.5, 128.3, 125.9, 125.7, 121.8, 50.3, 45.3, 28.4, 21.3; *m/z* LRMS (ESI^+^) 329 [M+H]^+^; HRMS (ESI^+^): calc. for C_22_H_21_N_3_O [M+H]^+^ requires 329.1648 found 329.1646.

**4236 2-(4-methylbenzyl)-6-(pyridin-3-yl)-3,4-dihydroisoquinolin-1(2H)-one**

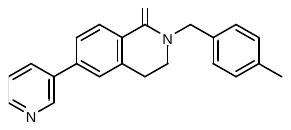

Following **general procedure 3**, **3698** (80 mg, 0.243 mmol) and 3-pyridyl boronic acid (33 mg, 0.267 mmol) afforded the title product **4236** (76 mg, 99%) as a brown oil that solidified on standing after purification on silica gel (MeOH:CH_2_Cl_2_, 4%); ^1^H NMR (400 MHz, Acetone) δ 8.92 (d, *J* = 2.6 Hz, 1H), 8.60 (dd, *J* = 4.9, 1.6 Hz, 1H), 8.15 (d, *J* = 8.1 Hz, 1H), 8.11 – 8.03 (m, 1H), 7.71 (dd, *J* = 8.1, 1.9 Hz, 1H), 7.62 (t, *J* = 1.2 Hz, 1H), 7.48 (dd, *J* = 8.1, 4.7 Hz, 1H), 7.28 (d, *J* = 7.9 Hz, 2H), 7.17 (d, *J* = 7.9 Hz, 2H), 4.76 (s, 2H), 3.57 (dd, *J* = 7.1, 6.2 Hz, 2H), 3.08 (t, *J* = 6.6 Hz, 2H), 2.31 (s, 3H); ^13^C NMR (101 MHz, CDCl_3_) δ 164.3, 148.9, 148.1, 141.0, 139.0, 137.3, 136.0, 134.8, 134.4, 132.2, 129.5, 129.3, 128.2, 128.2, 126.0, 125.7, 123.9, 50.3, 45.3, 28.3, 21.2; *m/z* LRMS (ESI^+^) 329 [M+H]^+^; HRMS (ESI^+^): calc. for C_22_H_21_N_3_O [M+H]^+^ requires 329.1648 found 329.1647.

**4237 6-(1-methyl-1H-pyrazol-5-yl)-2-(4-methylbenzyl)-3,4-dihydroisoquinolin-1(2H)-one**

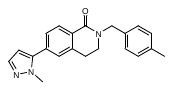

Following **general procedure 3**, **3698** (80 mg, 0.243 mmol) and 1-methylpyrazole-5- boronic acid (34 mg, 0.267 mmol) afforded the title product **4237** (71 mg, 88%) as a brown oil after purification on silica gel (MeOH:CH_2_Cl_2_, 6%); ^1^H NMR (400 MHz, MeOD) δ 8.07 (d, *J* = 8.0 Hz, 1H), 7.48 (dd, *J* = 2.0, 1.1 Hz, 1H), 7.44 (dd, *J* = 8.0, 1.7 Hz, 1H), 7.32 (t, *J* = 1.3 Hz, 1H), 7.19 (d, *J* = 7.8 Hz, 2H), 7.10 (d, *J* = 7.7 Hz, 2H), 6.39 (dd, *J* = 2.1, 1.0 Hz, 1H), 4.70 (s, 2H), 3.85 (d, *J* = 1.1 Hz, 3H), 3.51 – 3.43 (m, 2H), 2.94 (t, *J* = 6.6 Hz, 2H), 2.26 (s, 3H)^13^C NMR (101 MHz, MeOD) δ 165.7, 144.2, 140.6, 139.5, 138.4, 135.3, 135.1, 130.3, 130.1, 129.4, 129.0, 128.5, 128.2, 107.6, 46.5, 37.9, 28.7, 21.2; *m/z* LRMS (ESI^+^) 332 [M+H]^+^; HRMS (ESI^+^): calc. for C_21_H_22_N_3_O [M+H]^+^ requires 332.1763 found 332.1762.

**4238 6-(2-aminopyridin-4-yl)-2-(4-methylbenzyl)-3,4-dihydroisoquinolin-1(2H)-one**

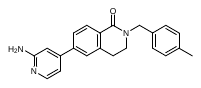

Following **general procedure 3**, **3698** (80 mg, 0.243 mmol) and 2-amino-pyridine-4-boronic pinacol ester (59 mg, 0.267 mmol) afforded the title product **4238** (79 mg, 95%) as a brown oil that solidified on standing after purification on silica gel (MeOH:CH_2_Cl_2_, 8%); ^1^H NMR (600 MHz, DMSO) δ 8.01 (d, *J* = 8.1 Hz, 1H), 7.99 (d, *J* = 5.0 Hz, 1H), 7.62 (dd, *J* = 8.1, 1.8 Hz, 1H), 7.55 (d, *J* = 1.9 Hz, 1H), 7.21 (d, *J* = 8.0 Hz, 2H), 7.15 (d, *J* = 7.9 Hz, 2H), 6.81 (dd, *J* = 5.4, 1.7 Hz, 1H), 6.74 (dd, *J* = 1.6, 0.8 Hz, 1H), 6.01 (s, 2H), 4.68 (s, 2H), 3.49 (t, *J* = 6.6 Hz, 2H), 3.01 (t, *J* = 6.6 Hz, 2H), 2.28 (s, 3H); ^13^C NMR (151 MHz, DMSO) δ 163.0, 160.4, 148.5, 147.3, 141.4, 139.4, 136.3, 134.6, 129.1, 128.3, 127.6, 125.4, 124.8, 110.0, 105.2, 49.4, 45.1, 27.4, 20.7; *m/z* LRMS (ESI^+^) 344 [M+H]^+^; HRMS (ESI^+^): calc. for C_22_H_22_N_3_O [M+H]^+^ requires 344.1763 found 344.1764.

**4239 2-(4-methylbenzyl)-6-(thiazol-5-yl)-3,4-dihydroisoquinolin-1(2H)-one**

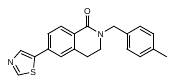

Following **general procedure 3**, **3698** (80 mg, 0.243 mmol) and thiazole boronic acid (56 mg, 0.267 mmol) afforded the title product 4239 (78 mg, 97%) as a light orange oil that solidified on standing after purification on silica gel (MeOH:CH_2_Cl_2_, 6%); ^1^H NMR (400 MHz, CDCl_3_) δ 8.82 (s, 1H), 8.19 (d, *J* = 8.1 Hz, 1H), 8.16 (s, 1H), 7.57 (dd, *J* = 8.0, 1.6 Hz, 1H), 7.37 (s, 1H), 7.24 (d, *J* = 14.2 Hz, 4H), 7.15 (d, *J* = 7.8 Hz, 2H), 4.76 (s, 2H), 3.50 (dd, *J* = 8.4, 4.8 Hz, 2H), 2.97 (t, *J* = 6.6 Hz, 2H), 2.34 (s, 3H); ^13^C NMR (126 MHz, CDCl_3_) δ 164.0, 139.2, 137.4, 134.3, 129.6, 129.6, 129.5, 129.5, 129.5, 128.3, 125.8, 125.5, 50.4, 45.2, 28.3, 21.3; *m/z* LRMS (ESI^+^) 335 [M+H]^+^; HRMS (ESI^+^): calc. for C_20_H_19_N_2_OS [M+H]^+^ requires 335.1218 found 335.1218.

#### NMR Spectra

3700

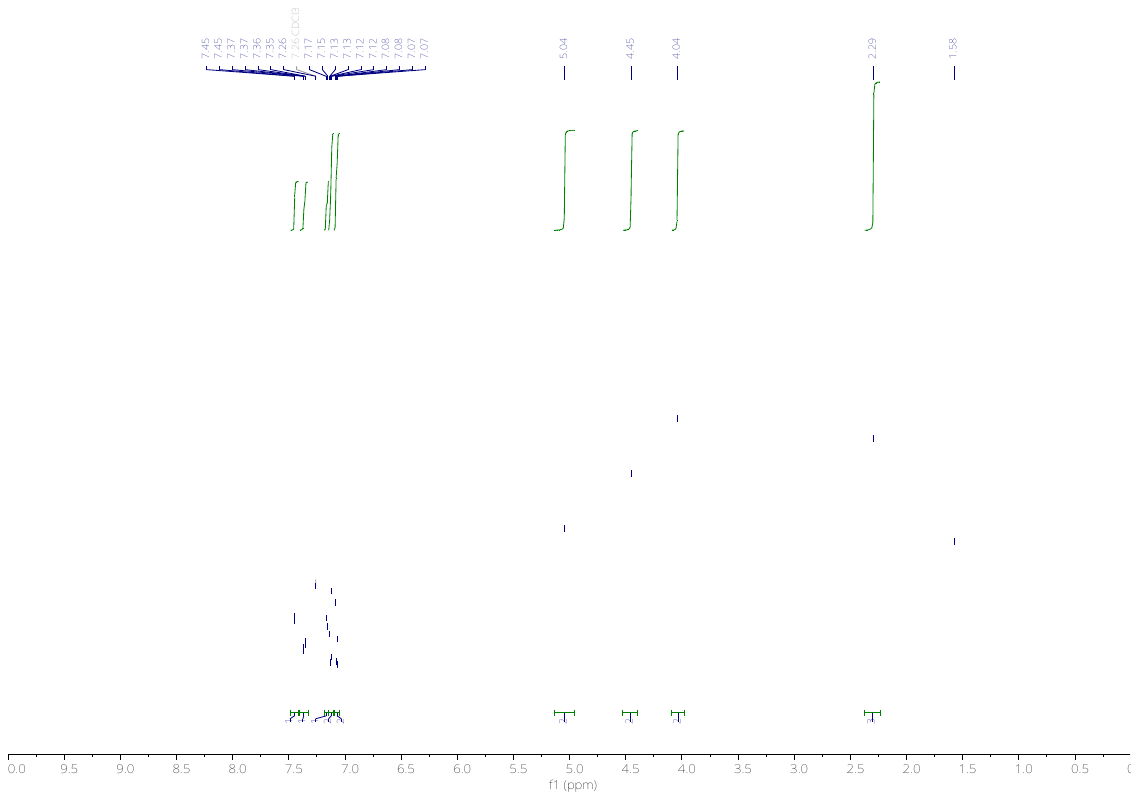

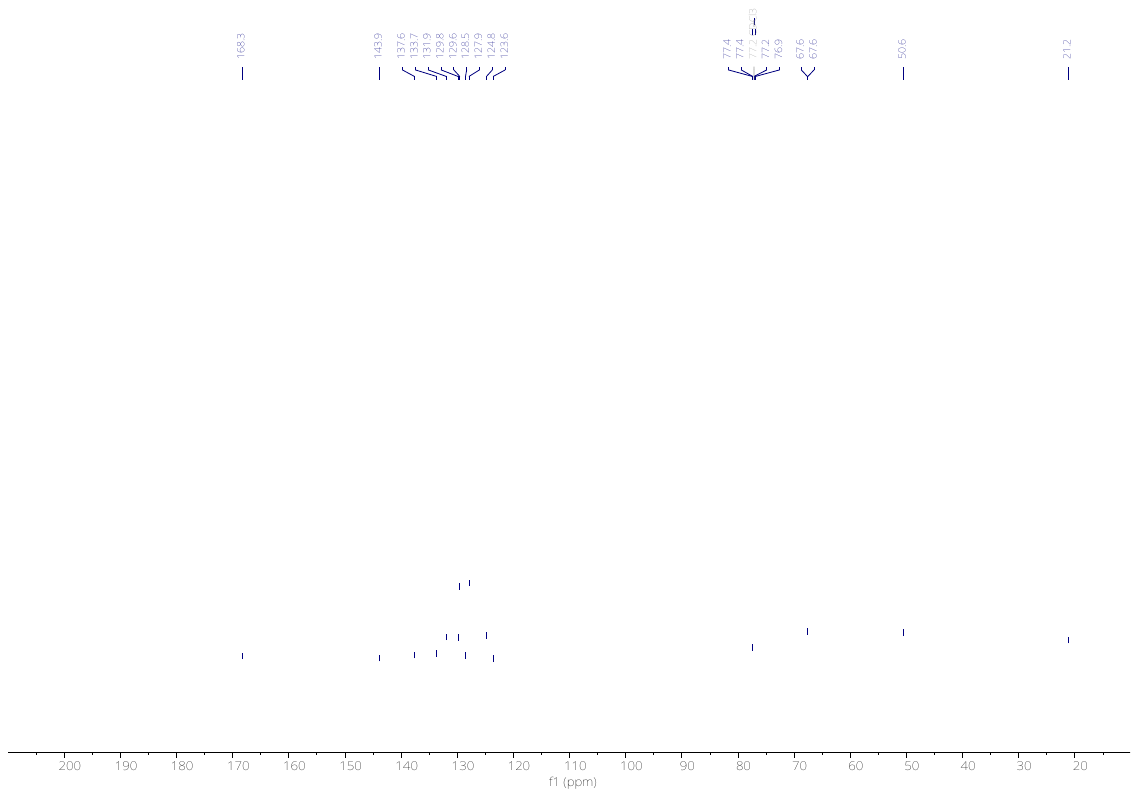

3702

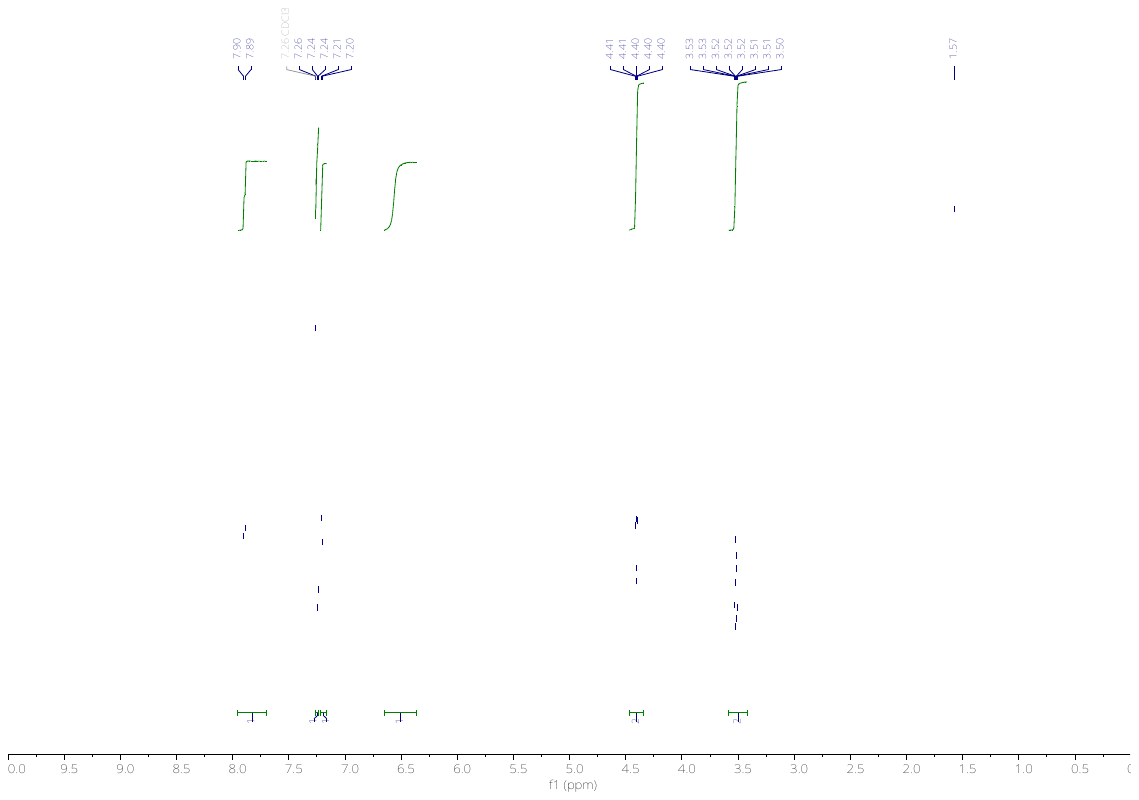

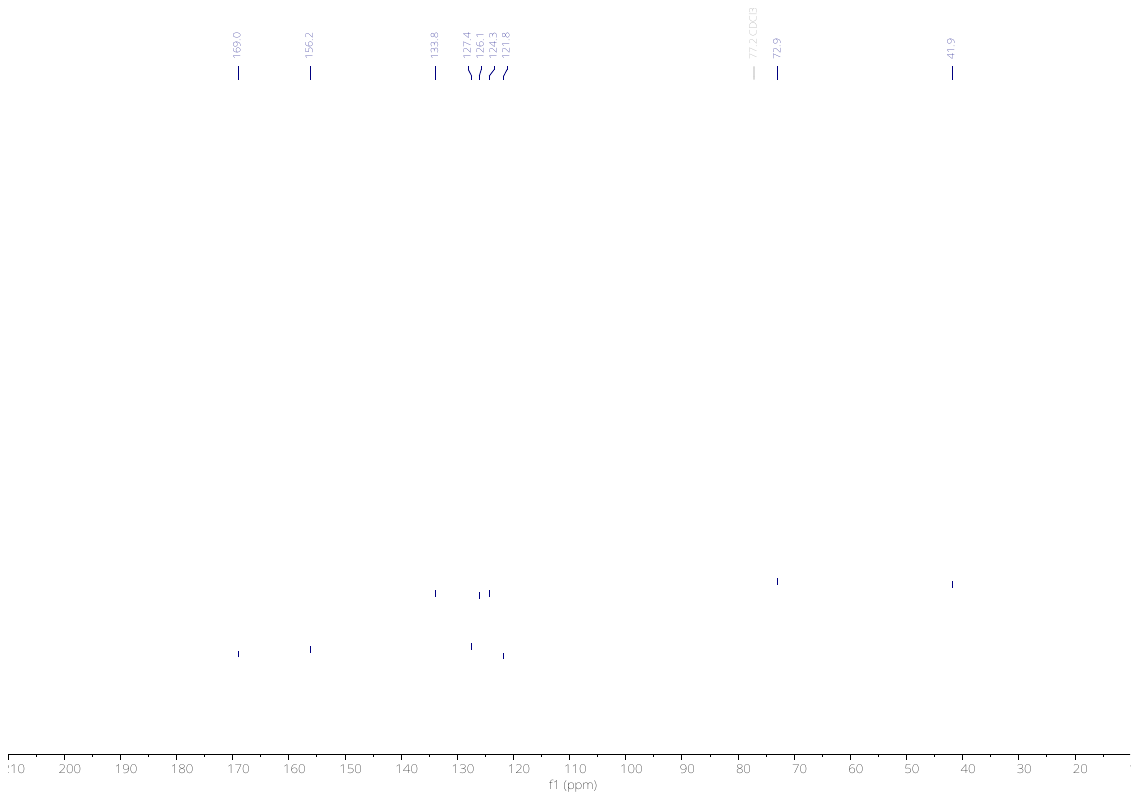

3703

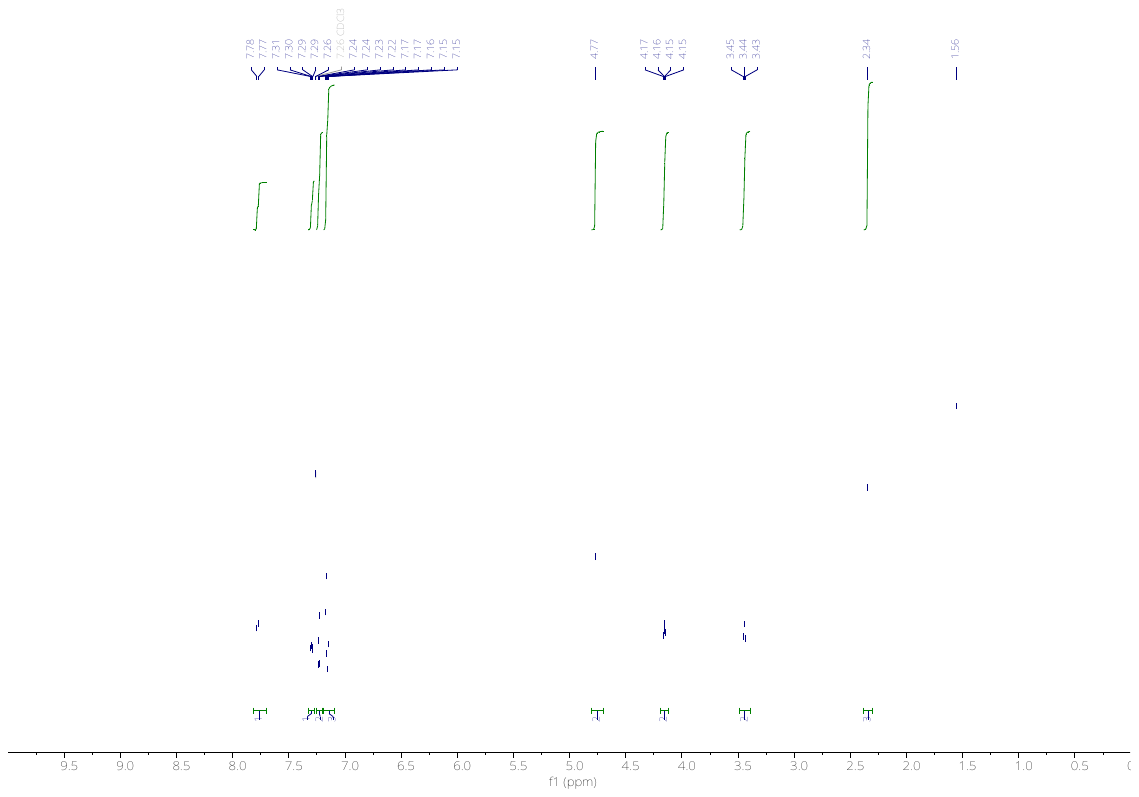

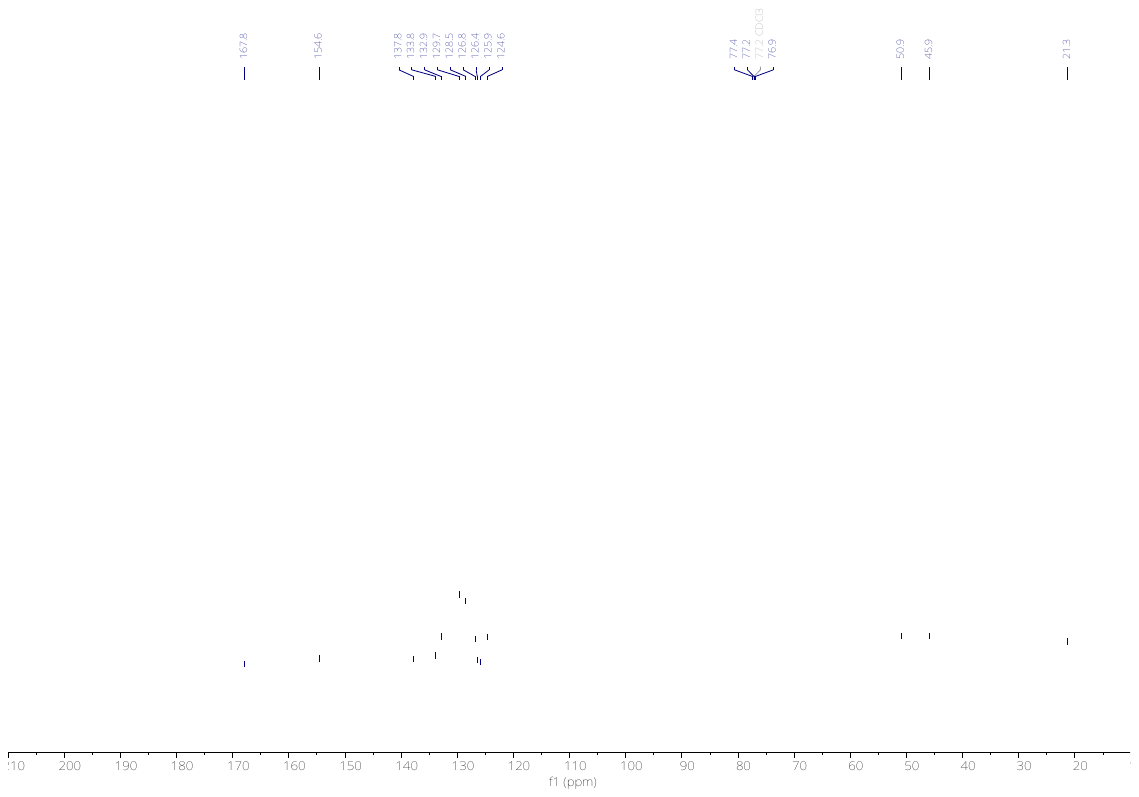

3705

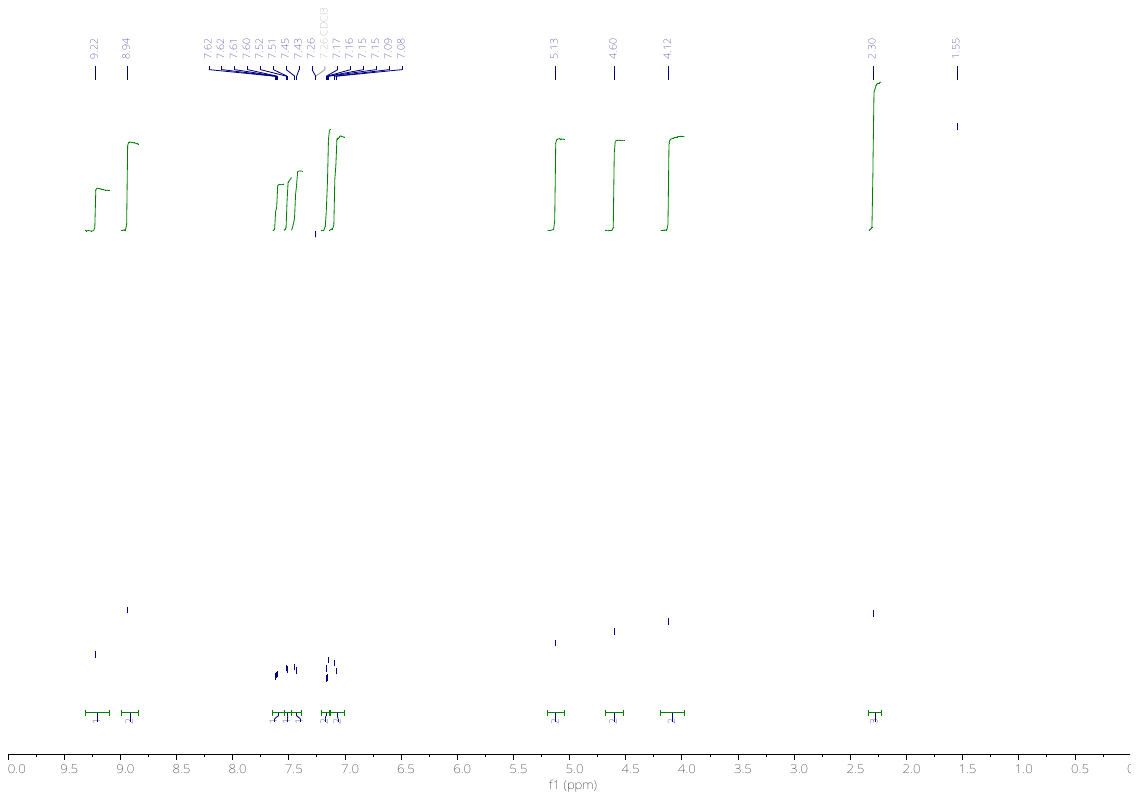

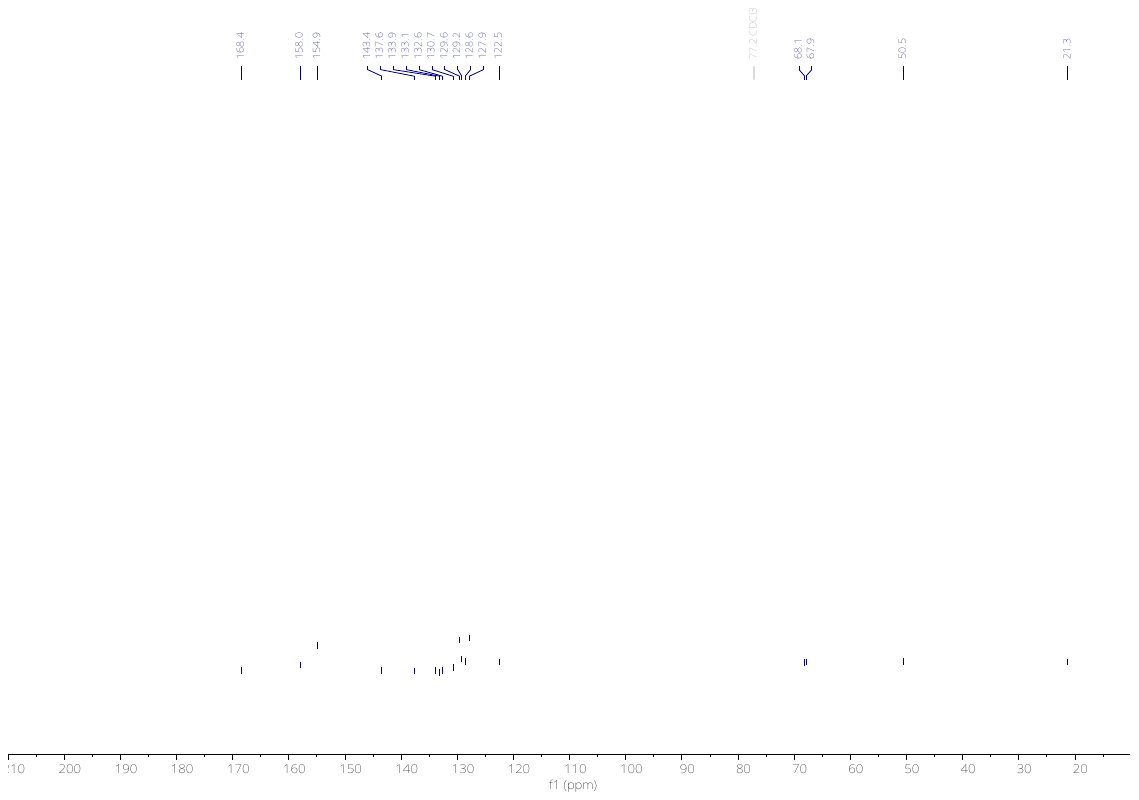

3706

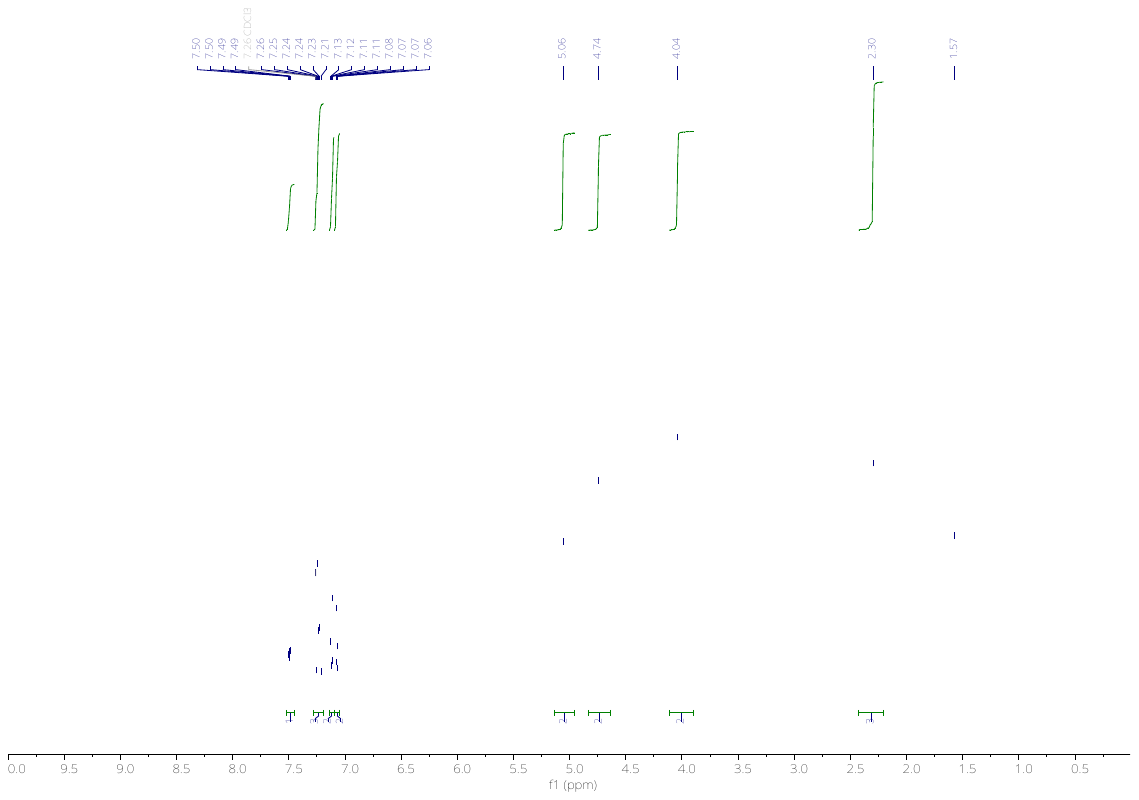

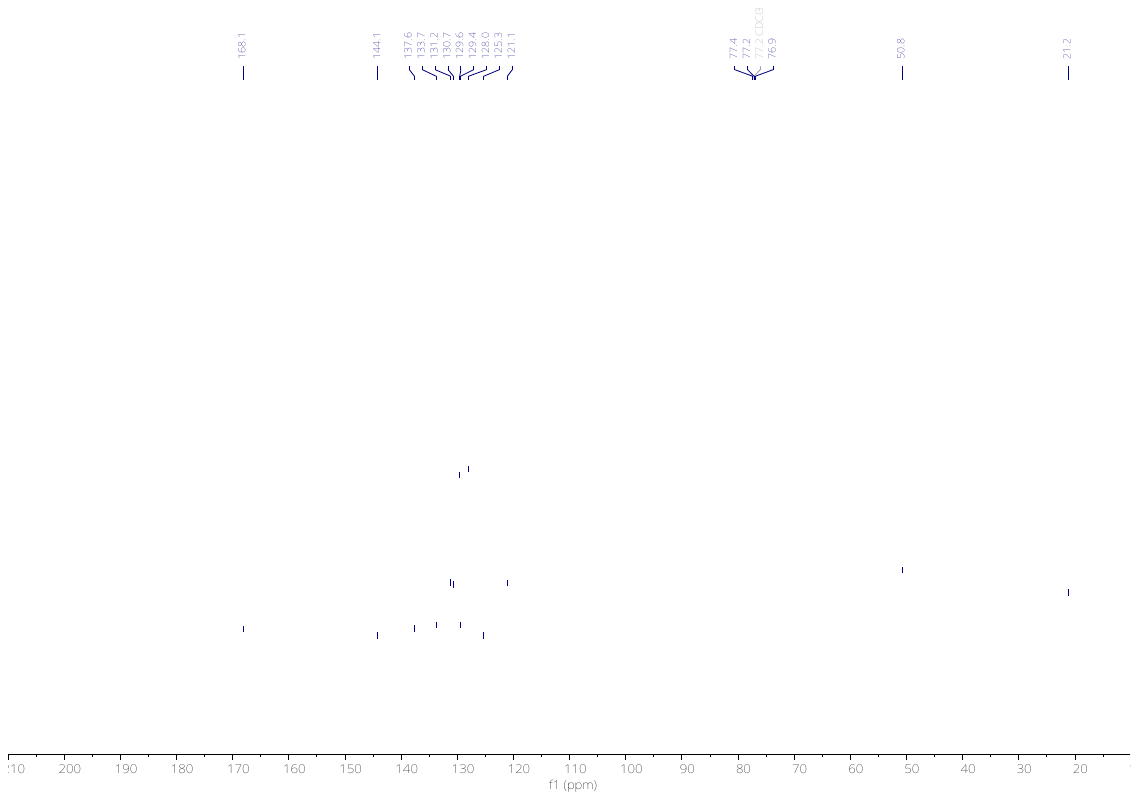

3827

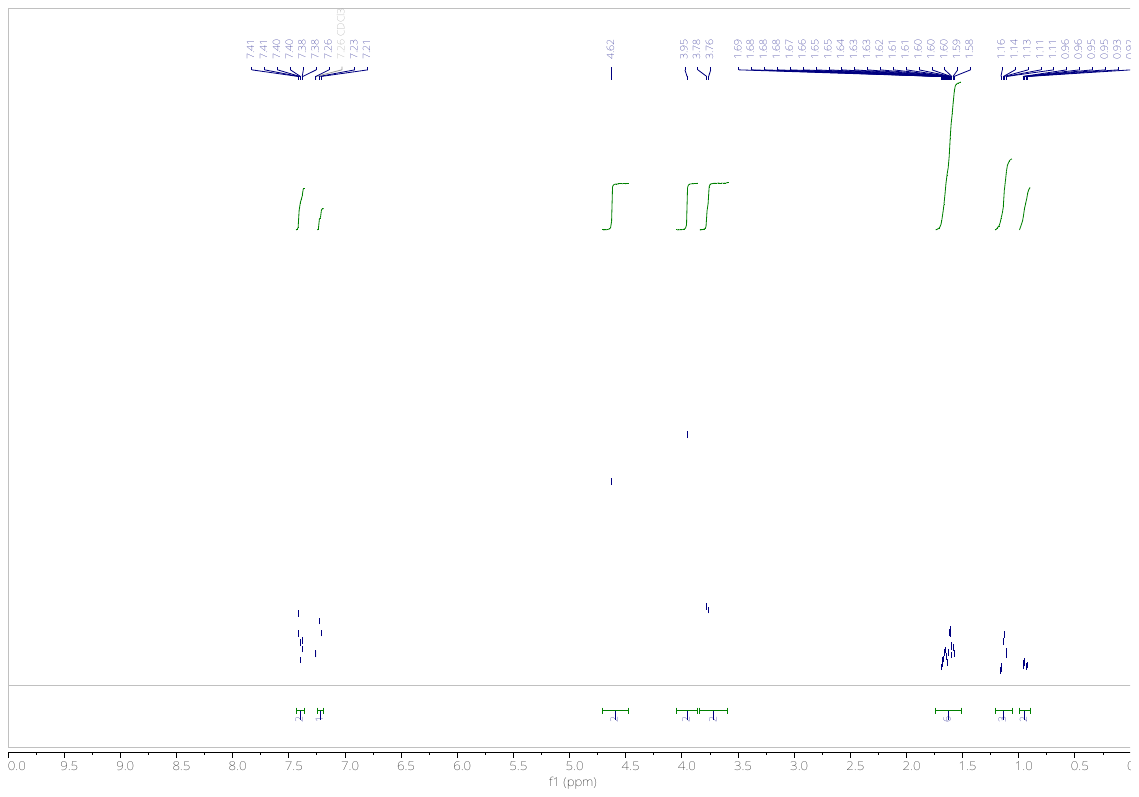

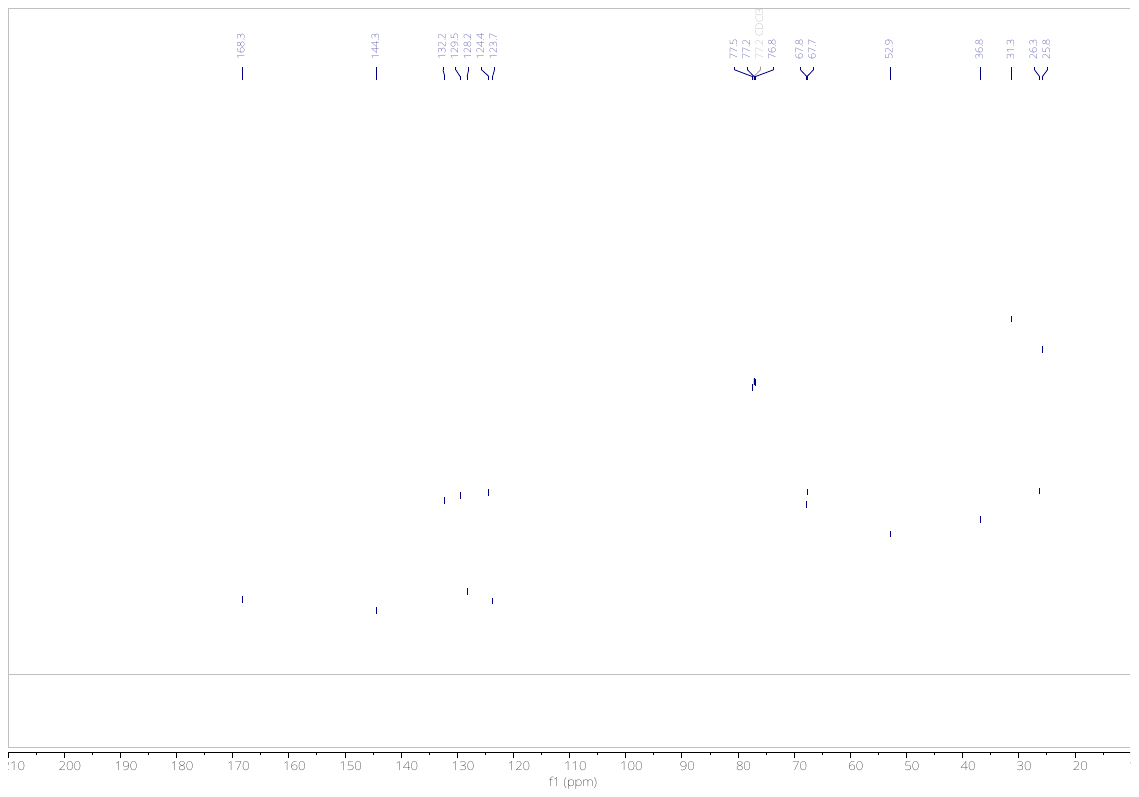

3829

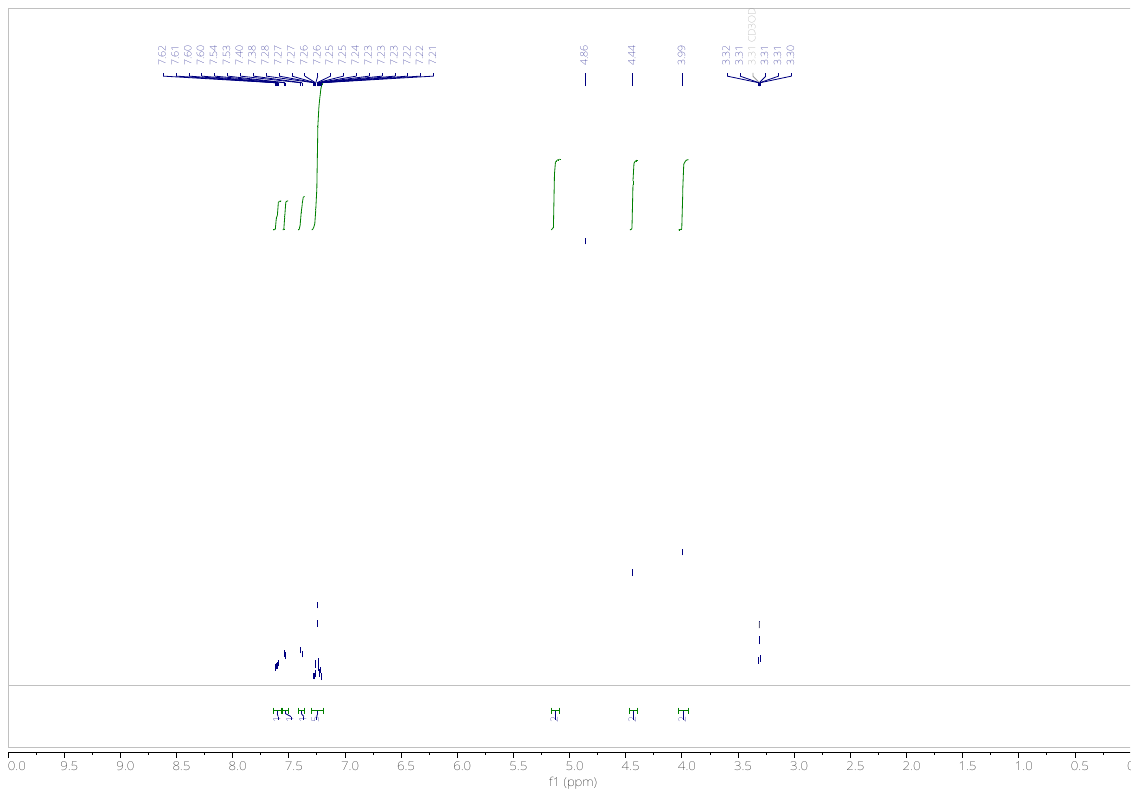

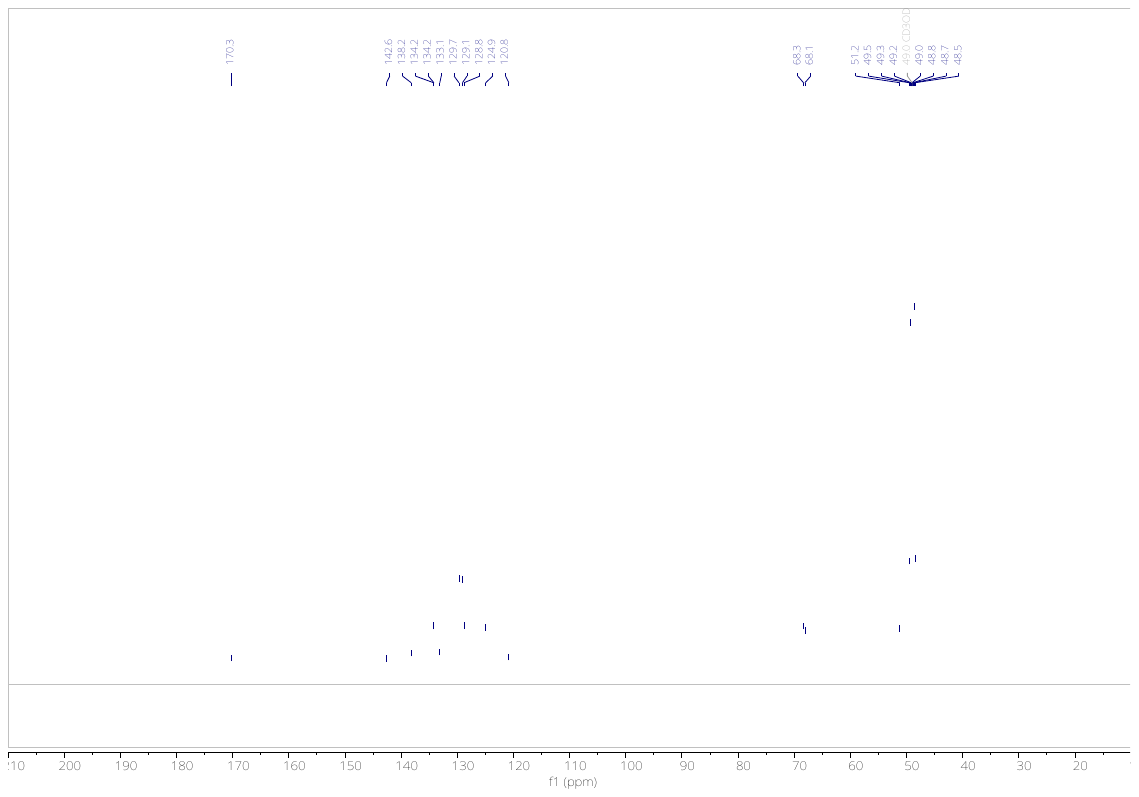

3698

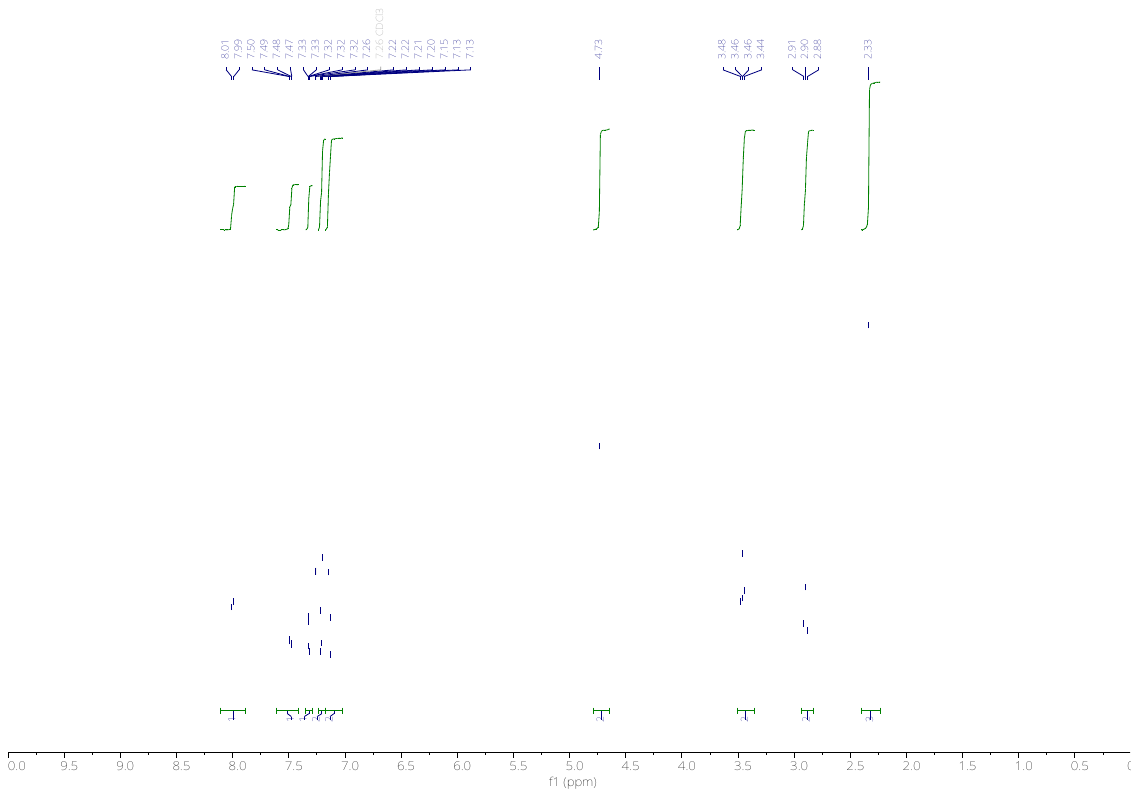

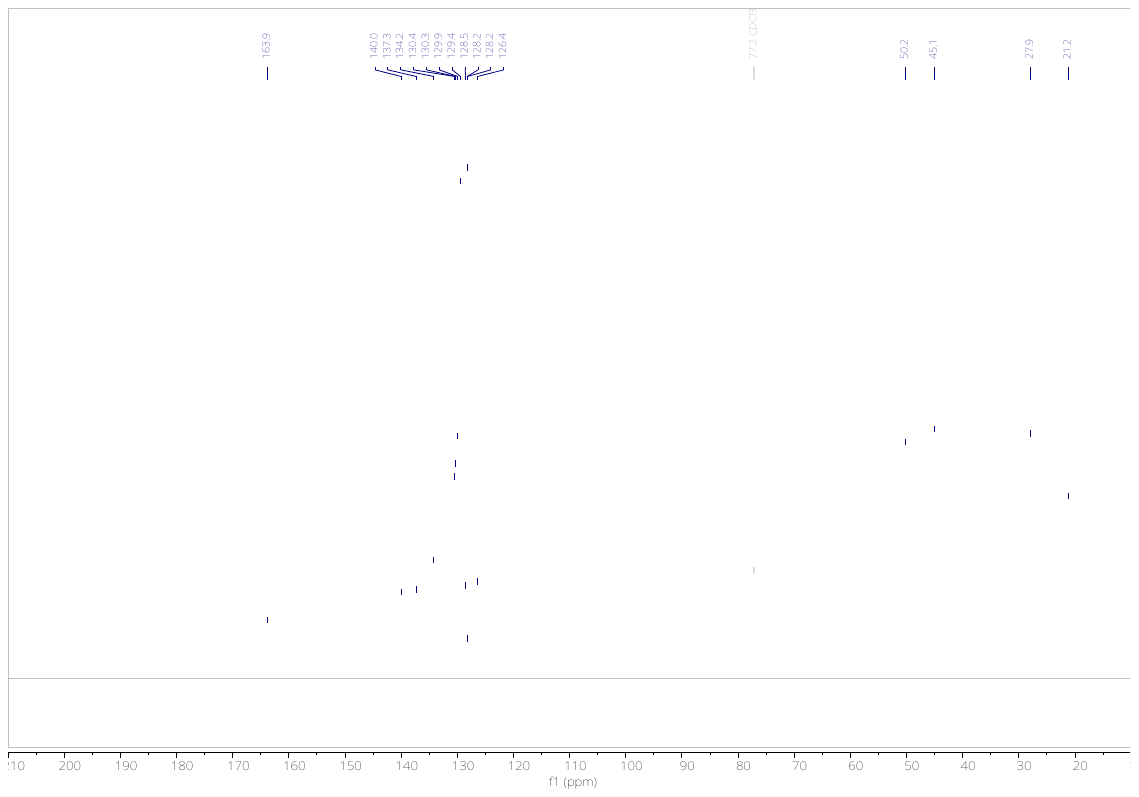

4119

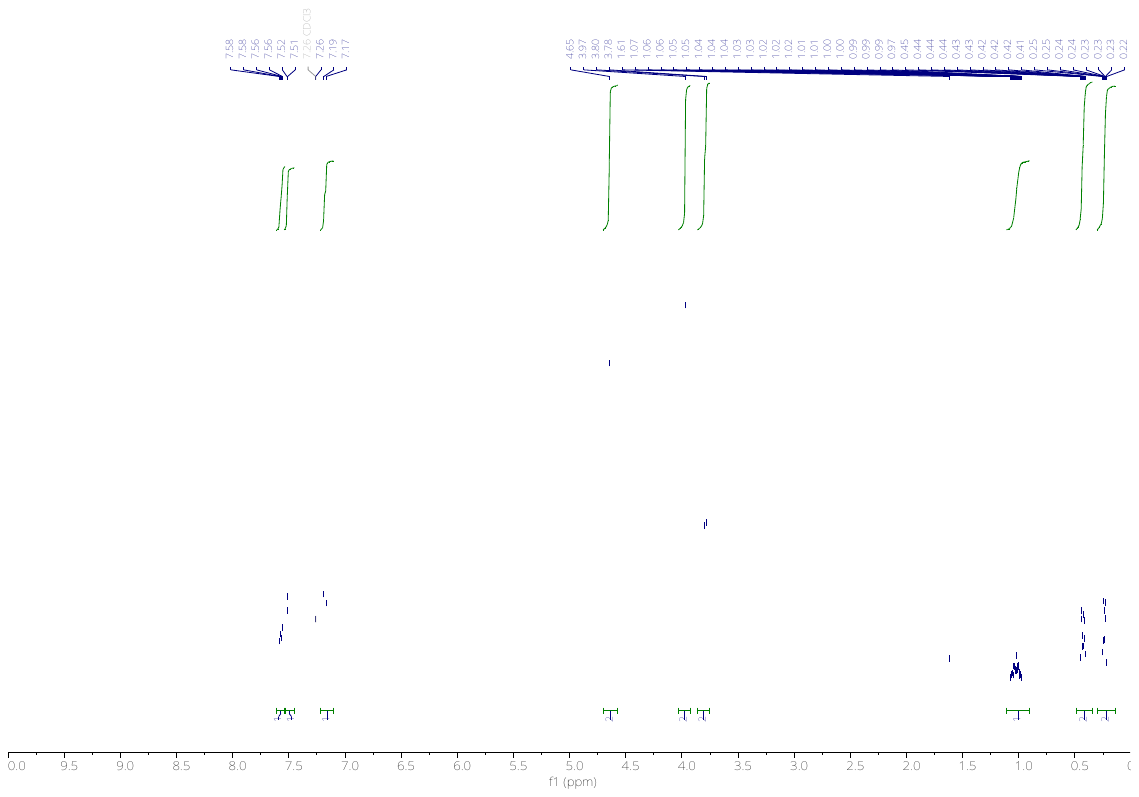

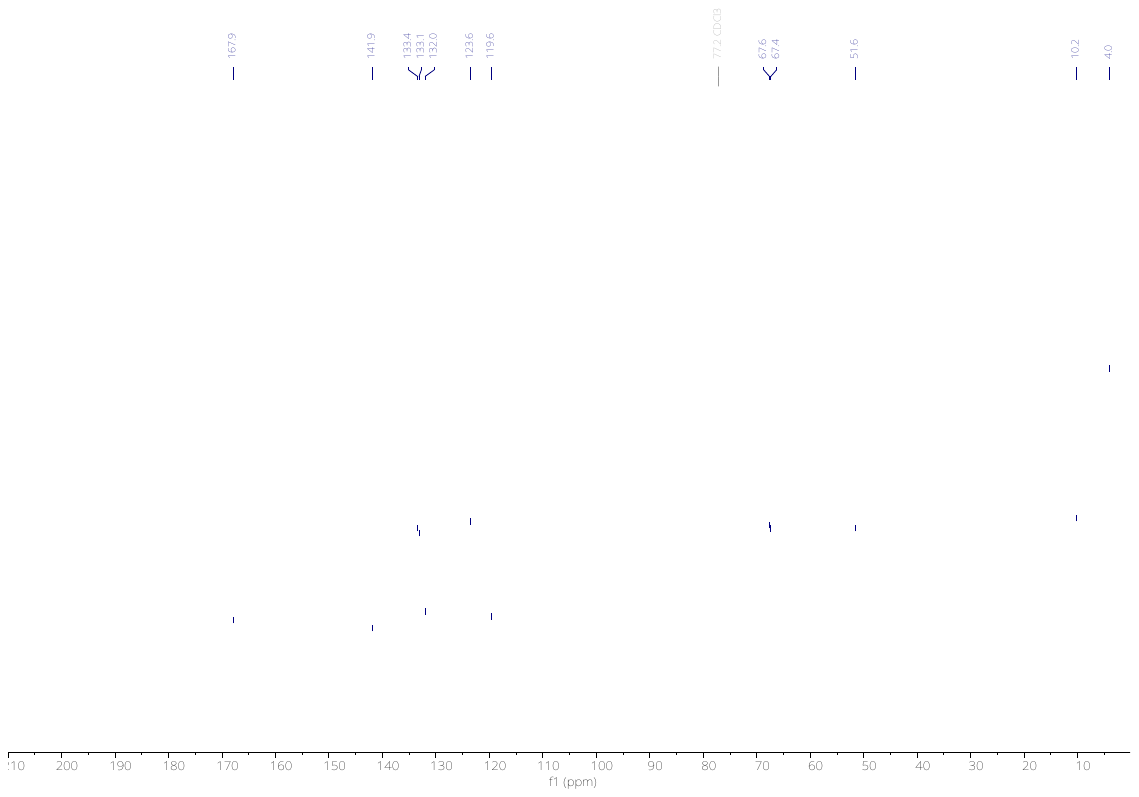

4118a

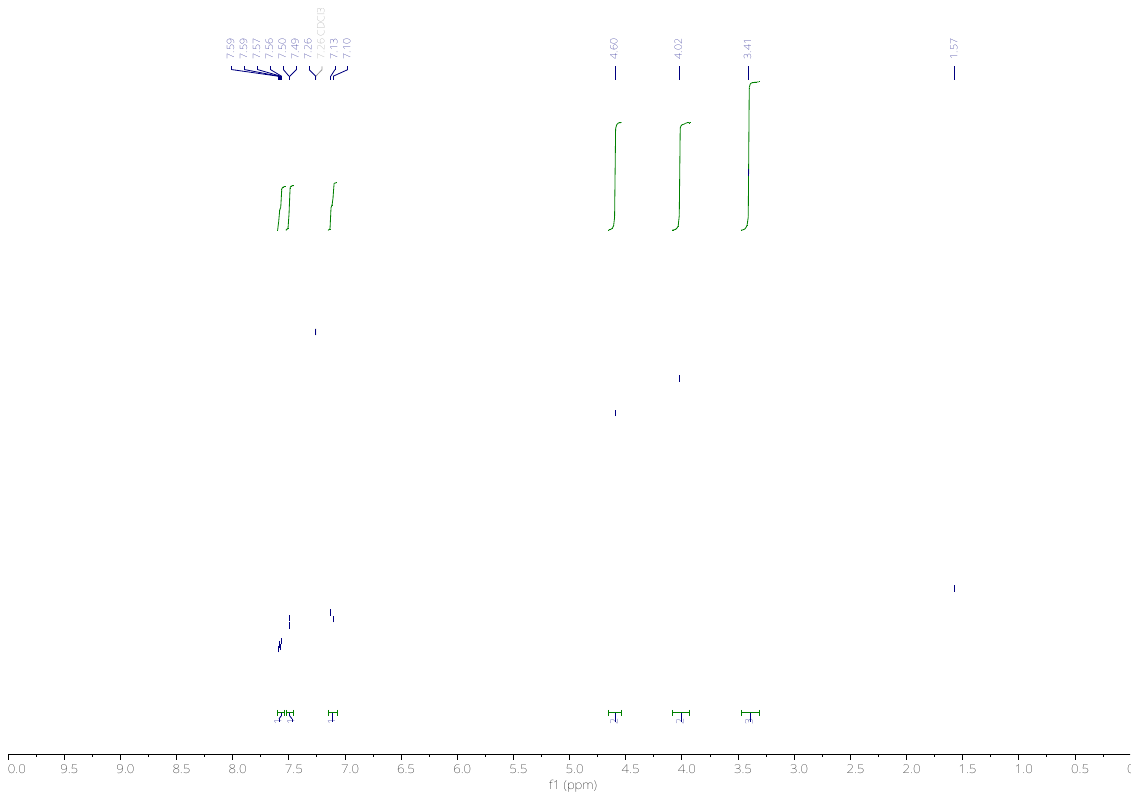

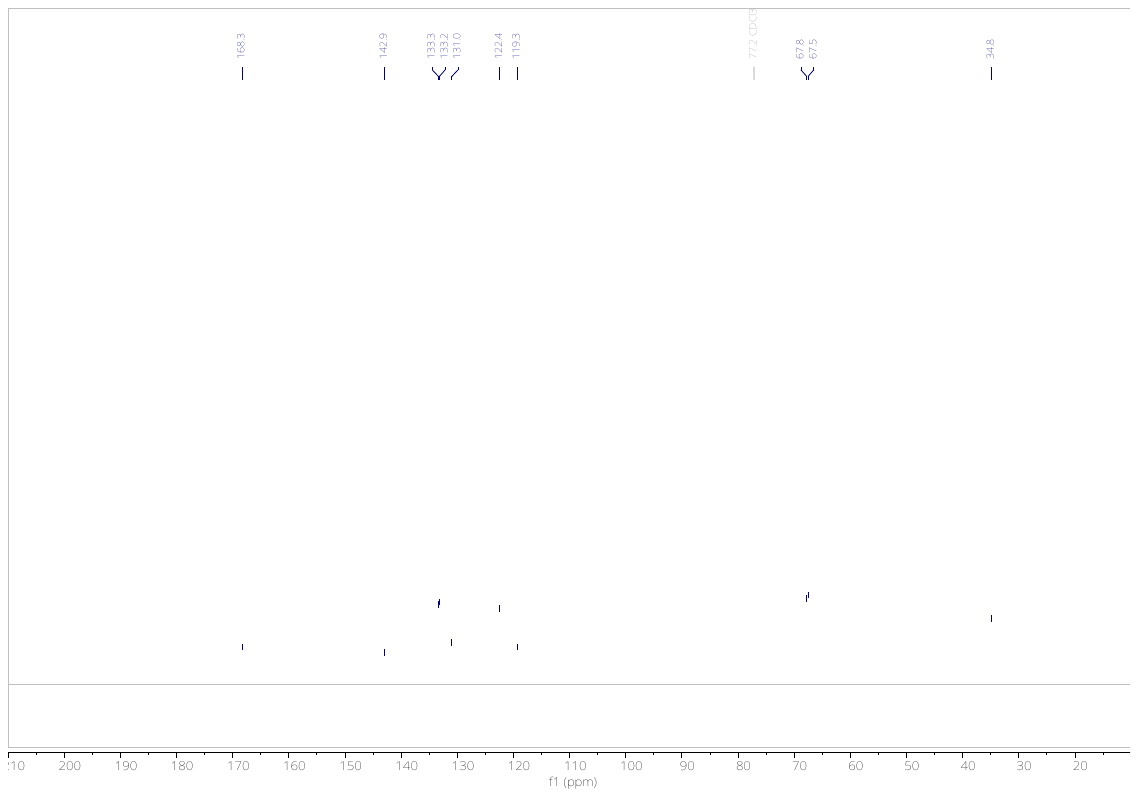

3145

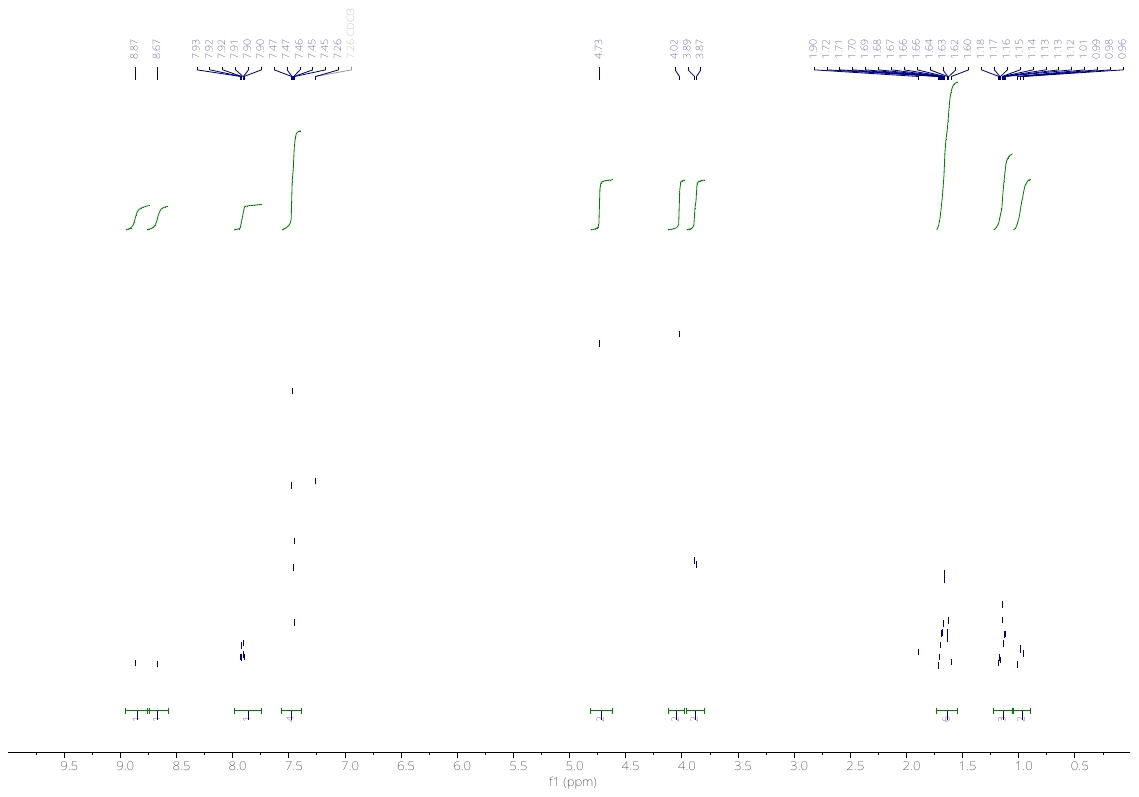

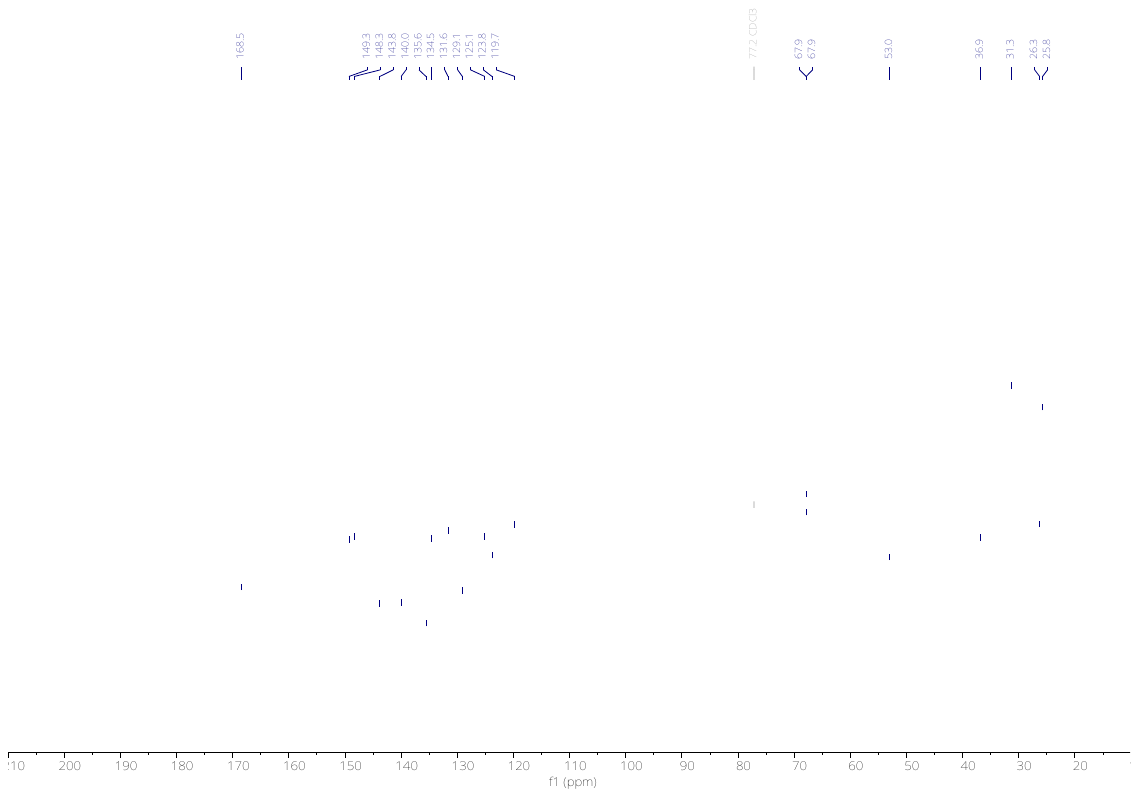

3593

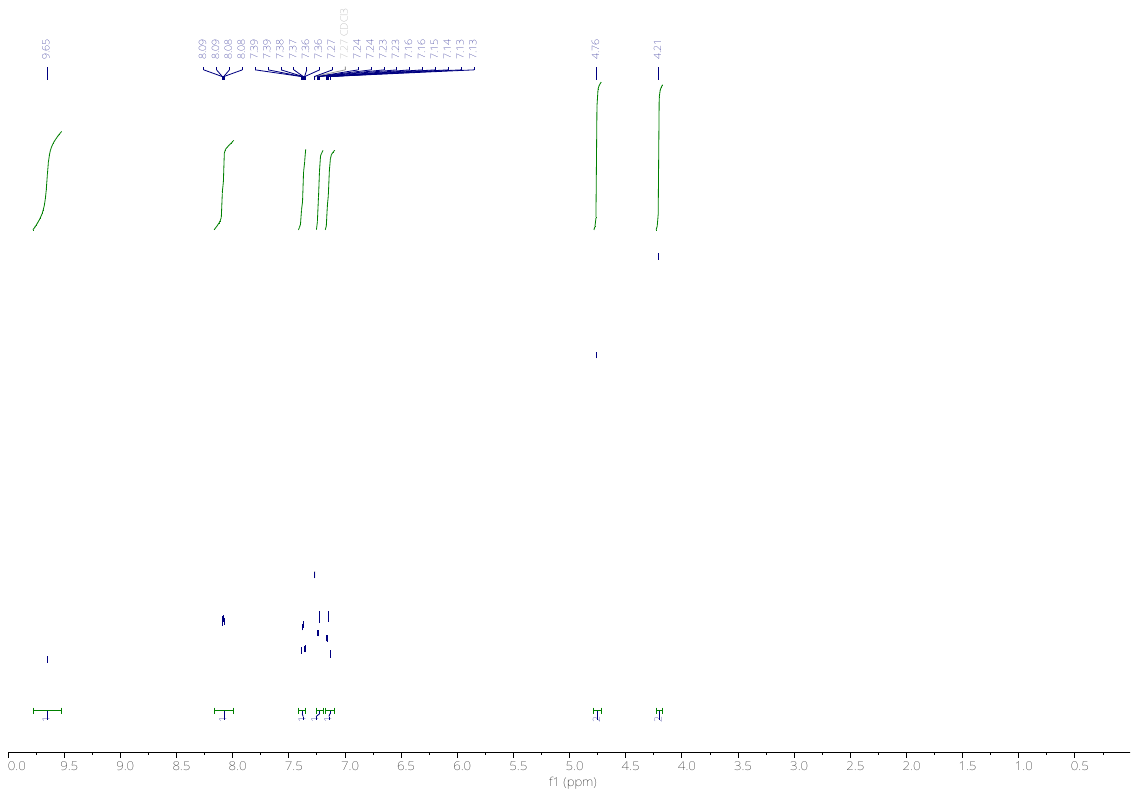

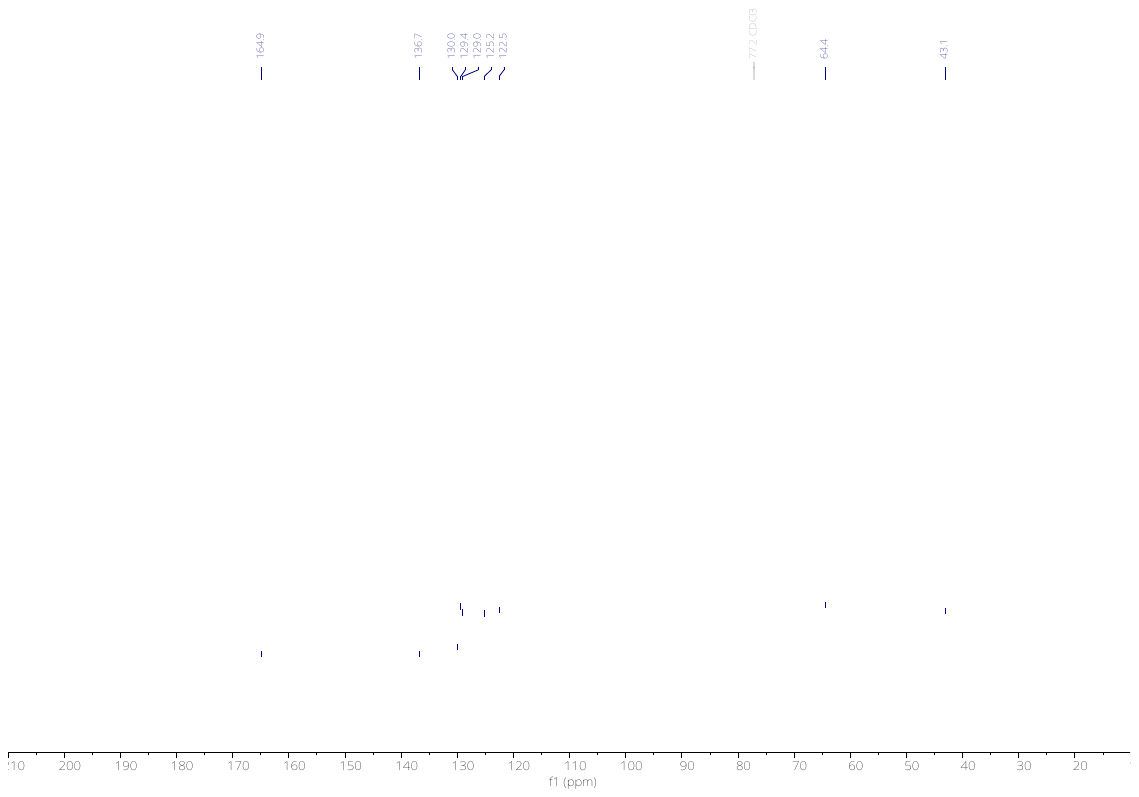

3594

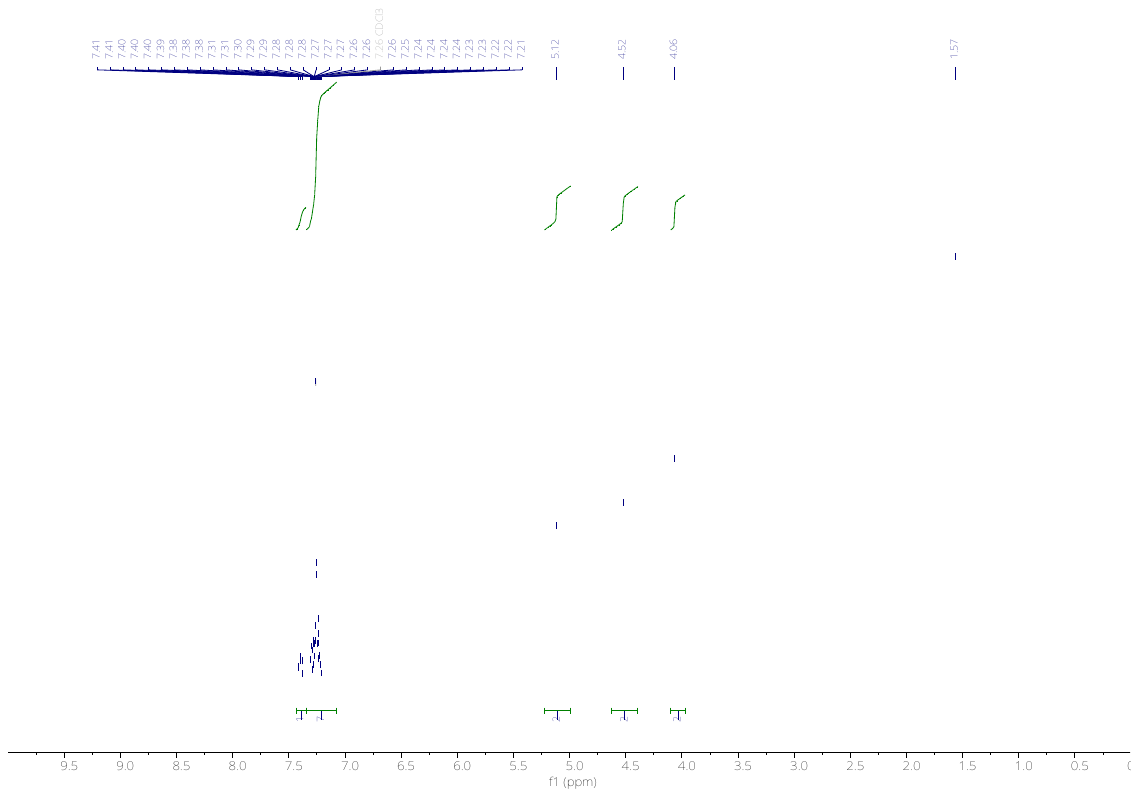

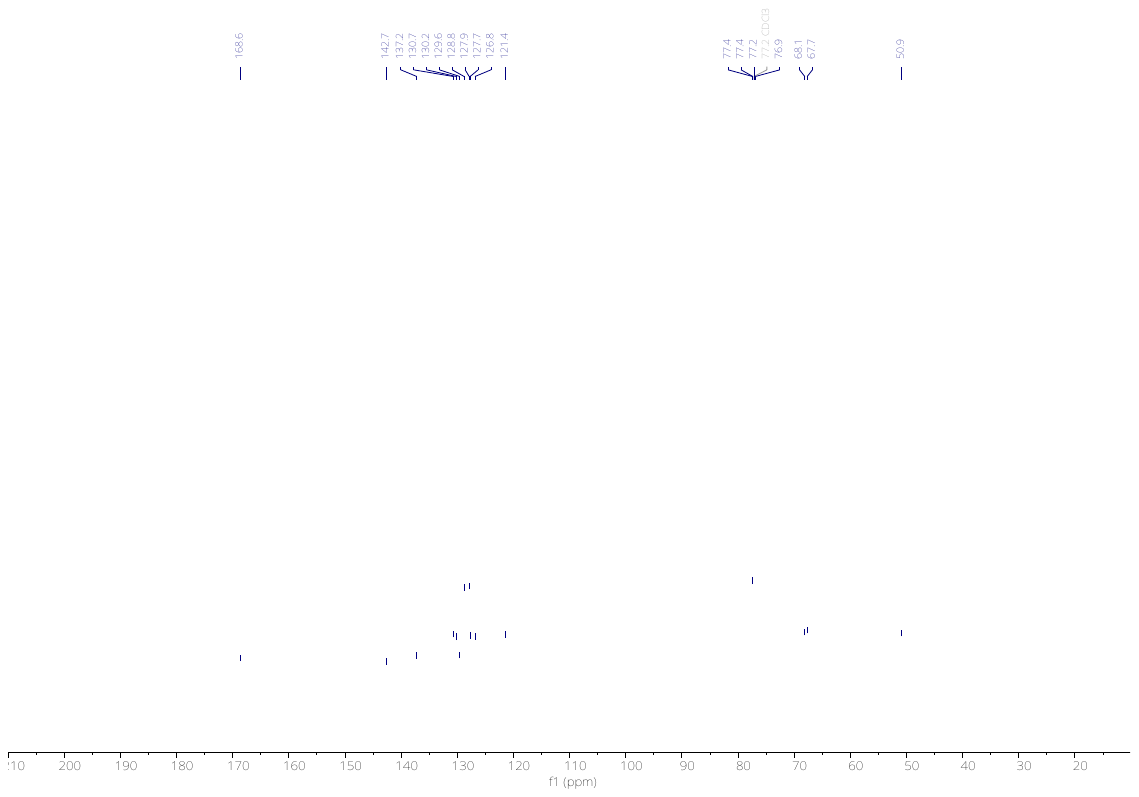

3596

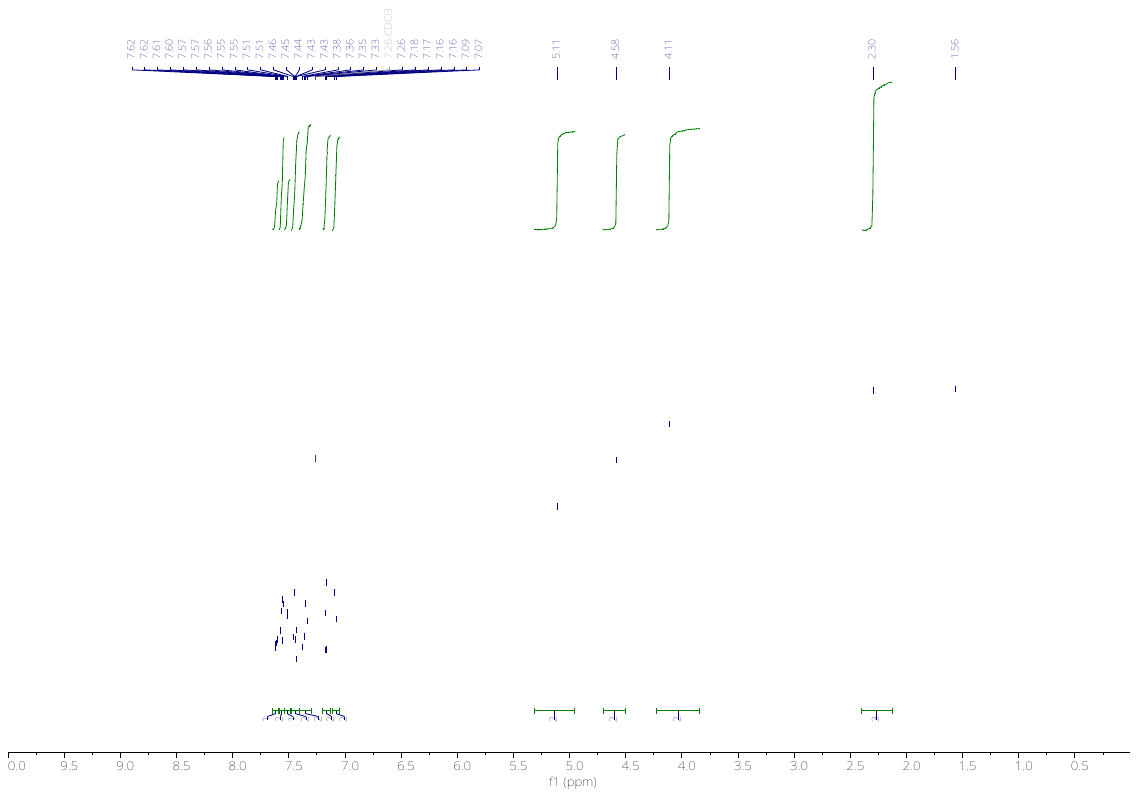

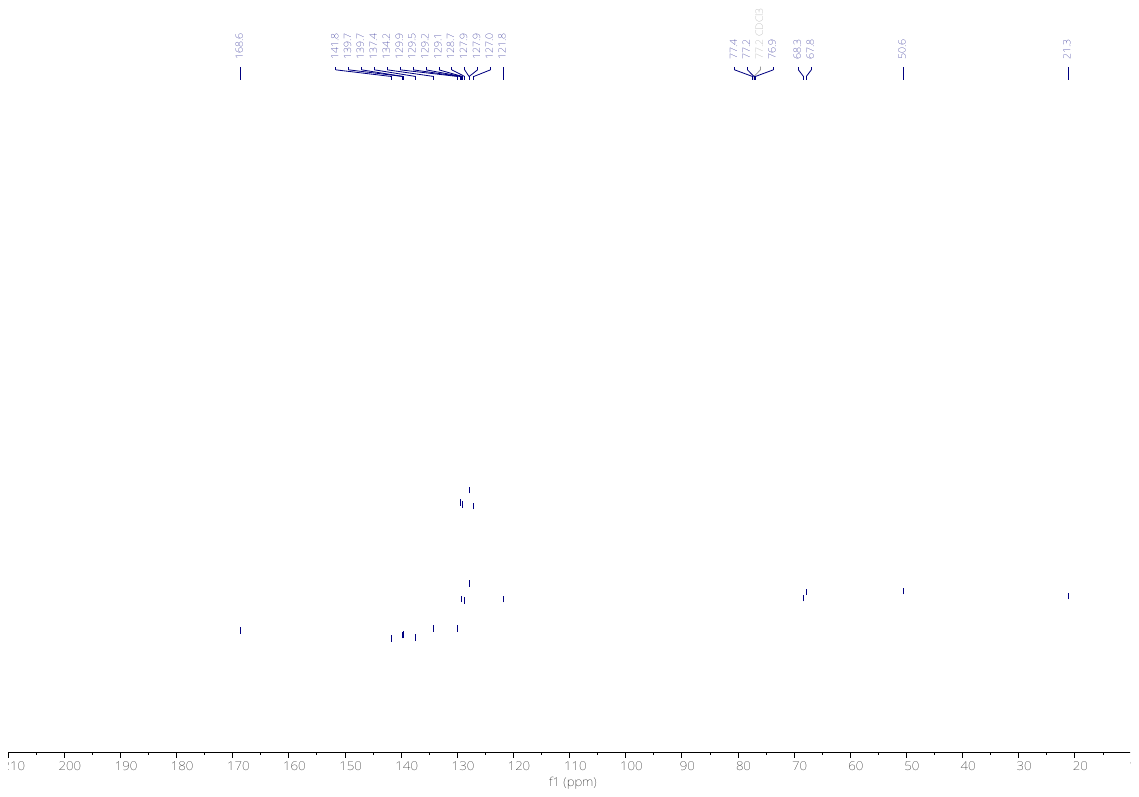

3599

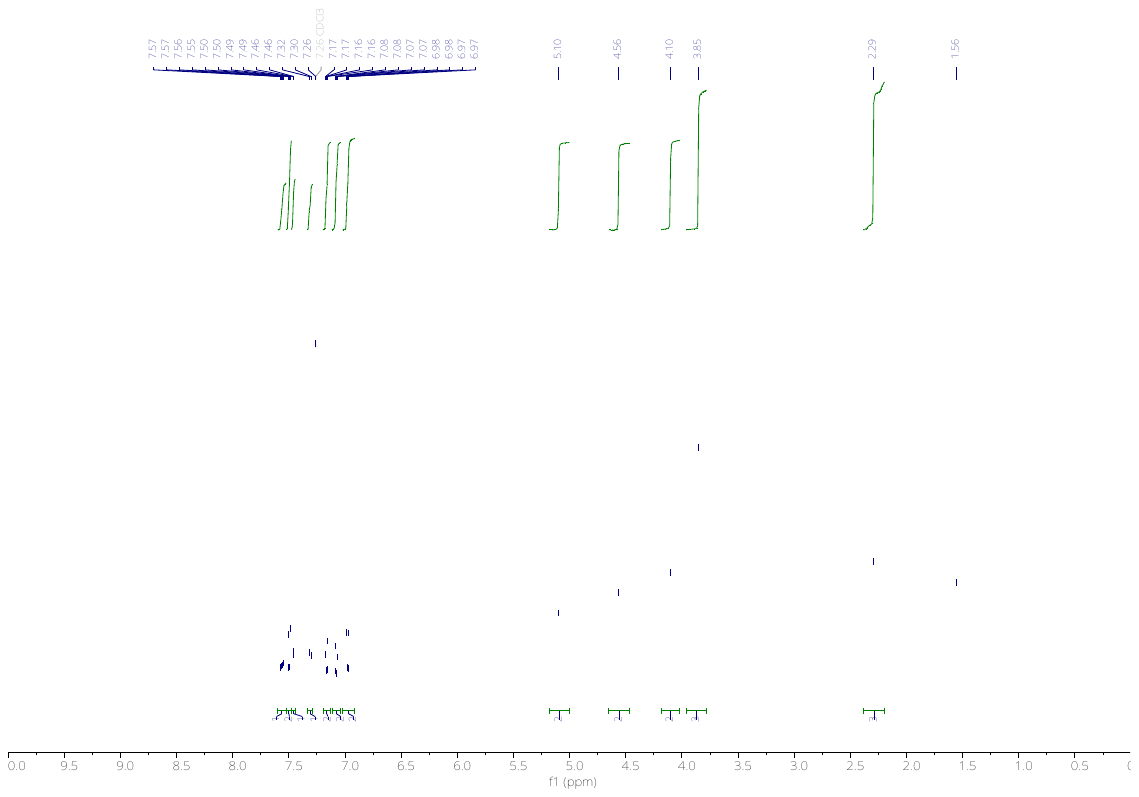

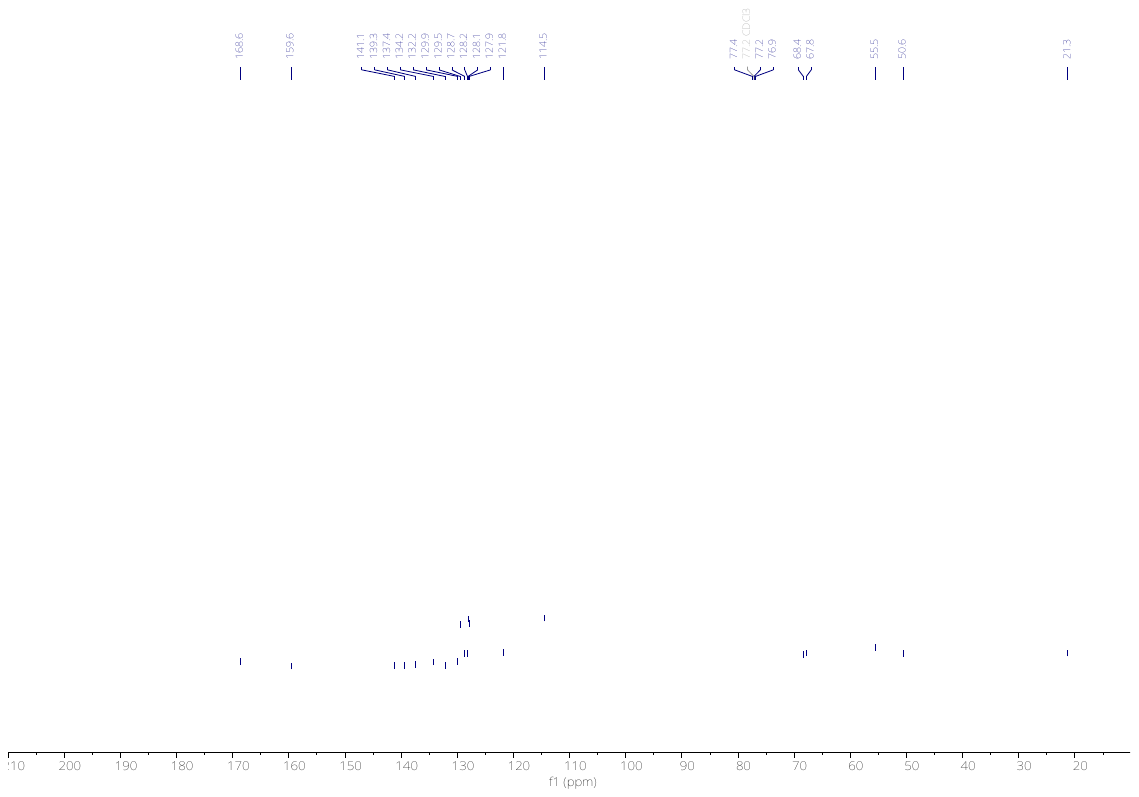

3600

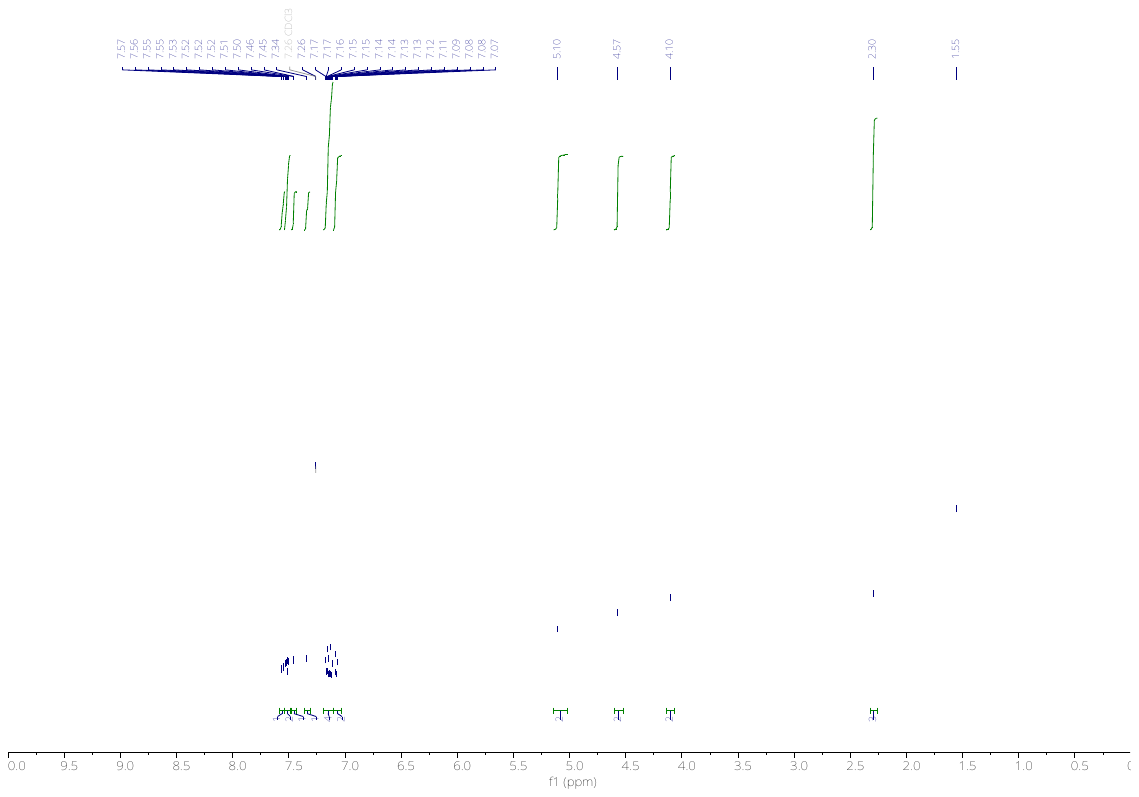

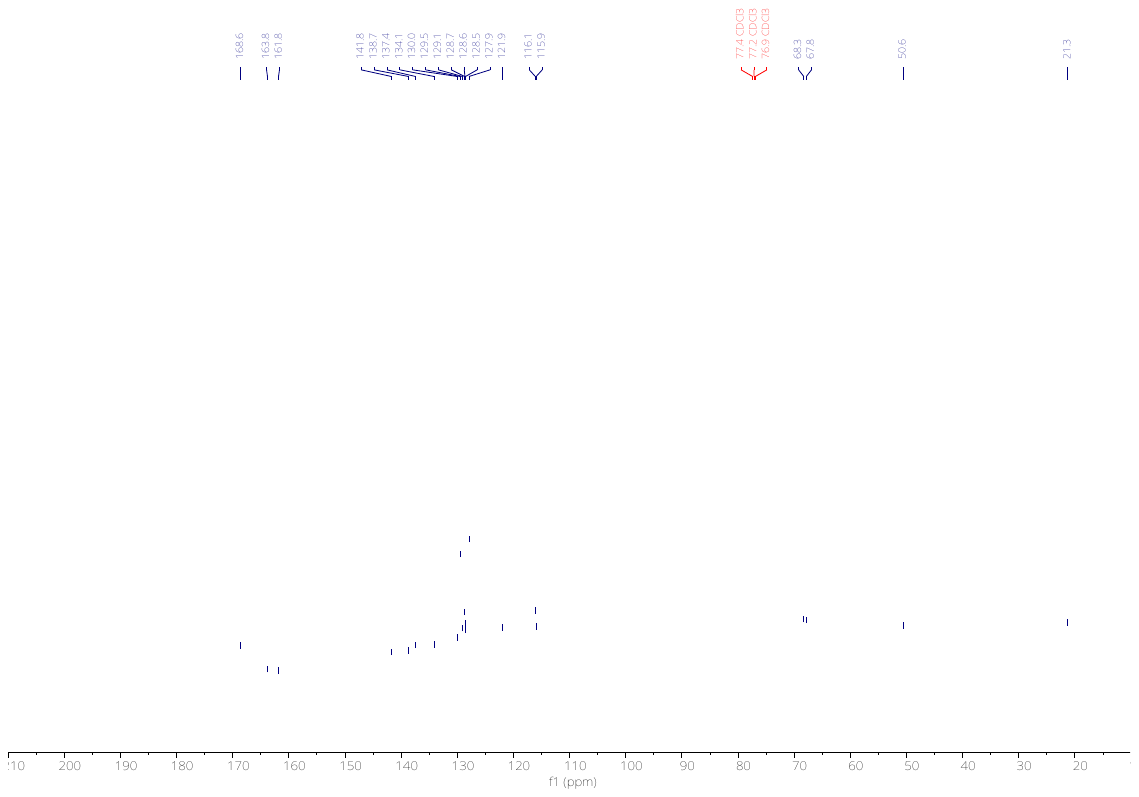

3601

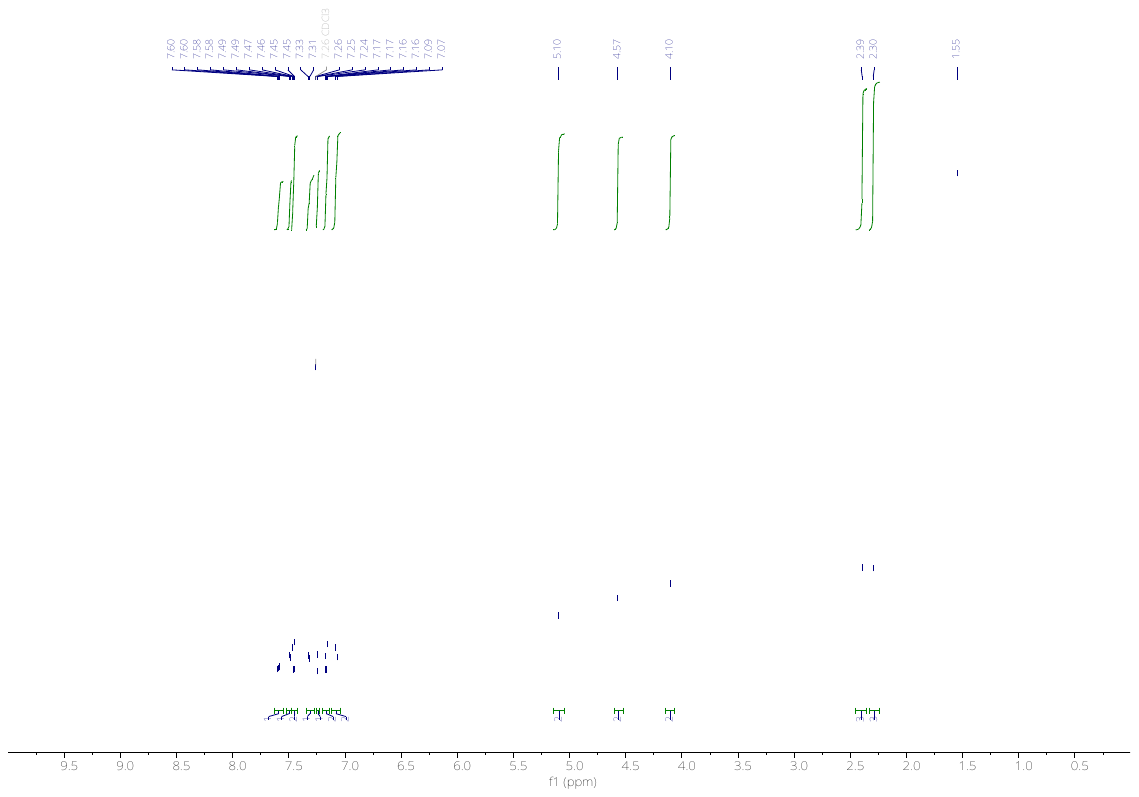

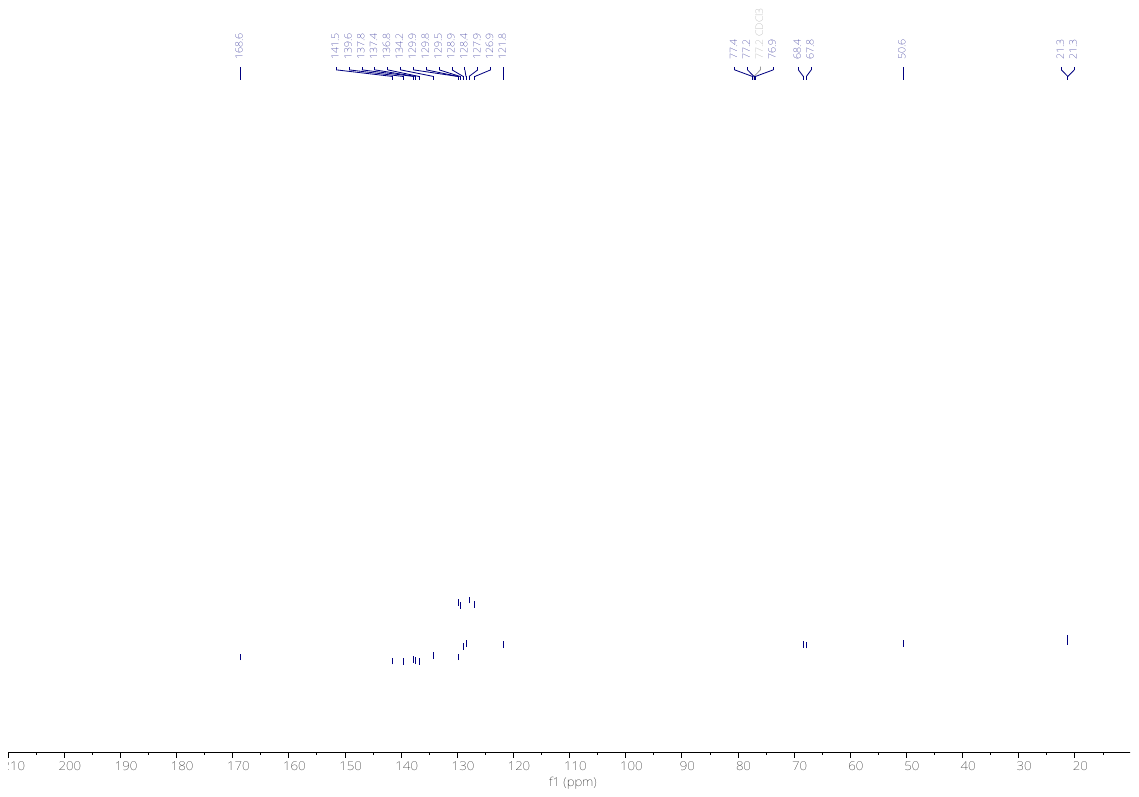

3697

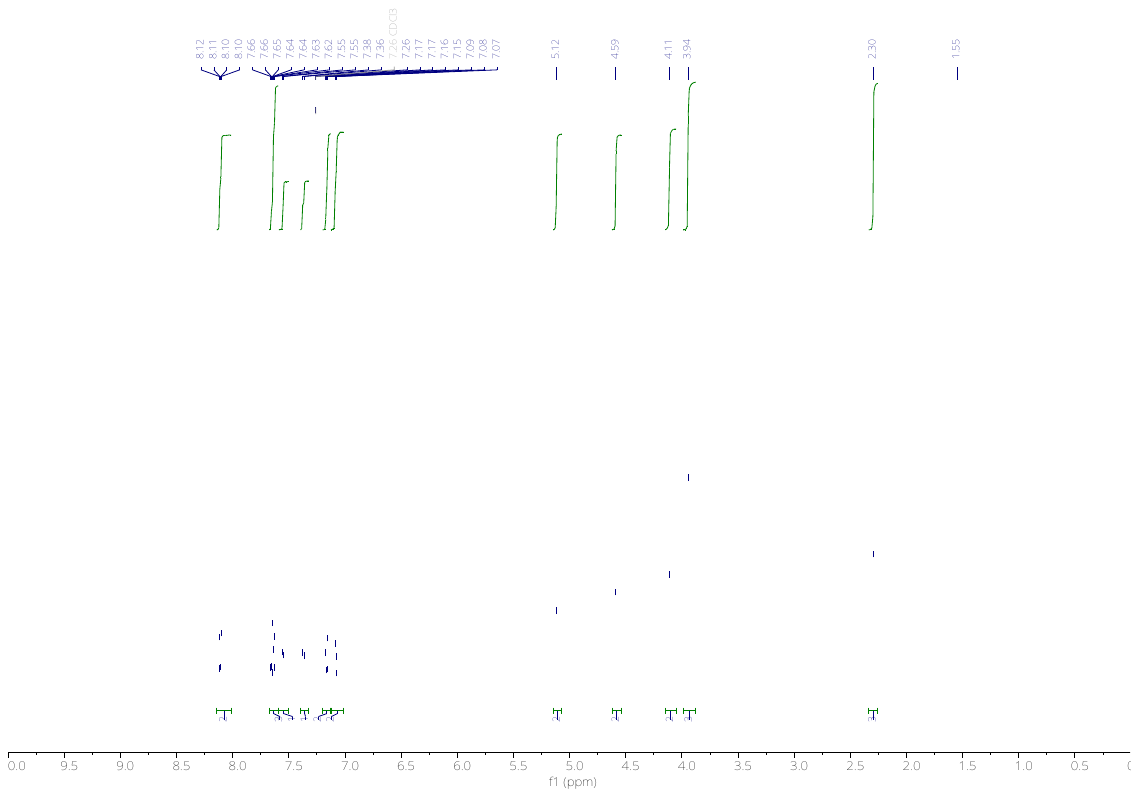

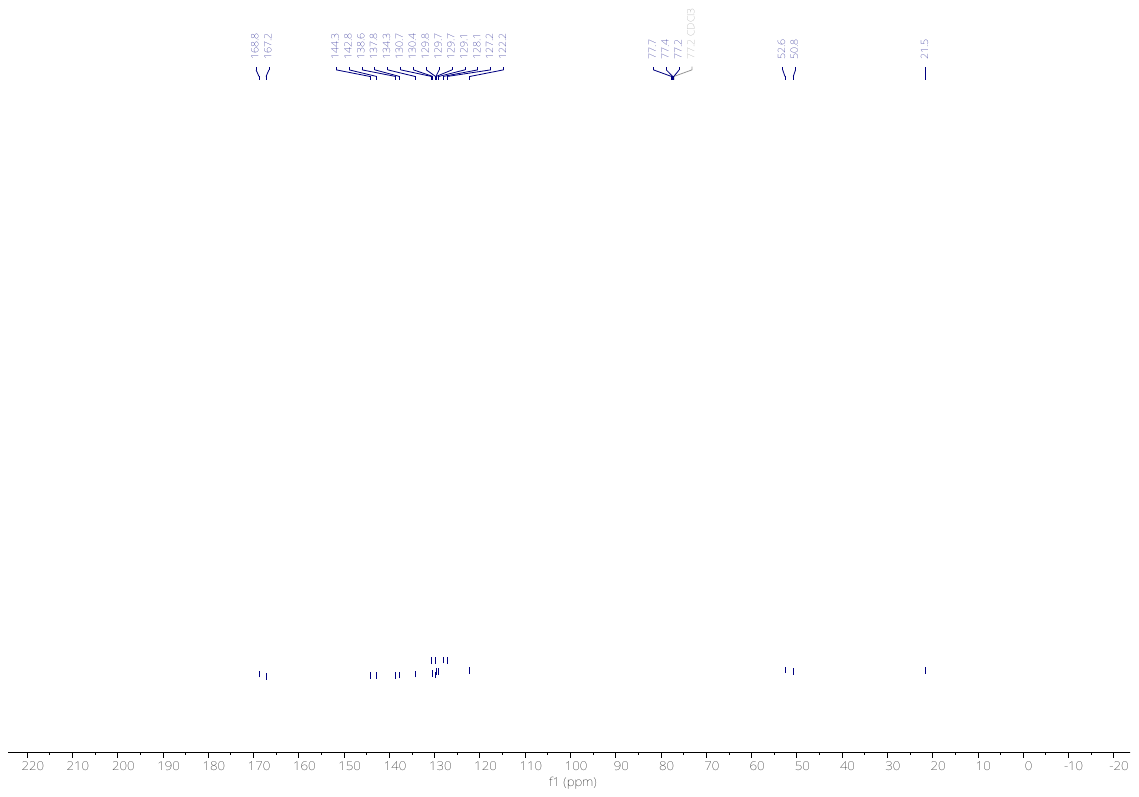

3701

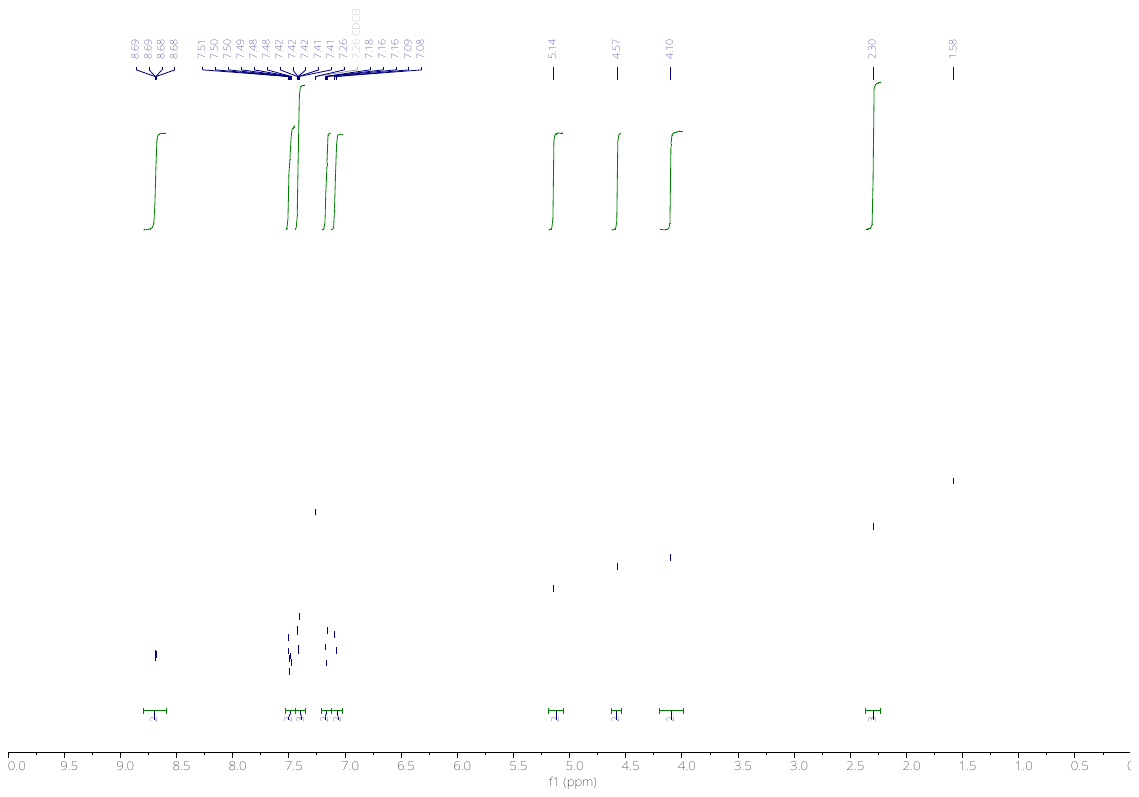

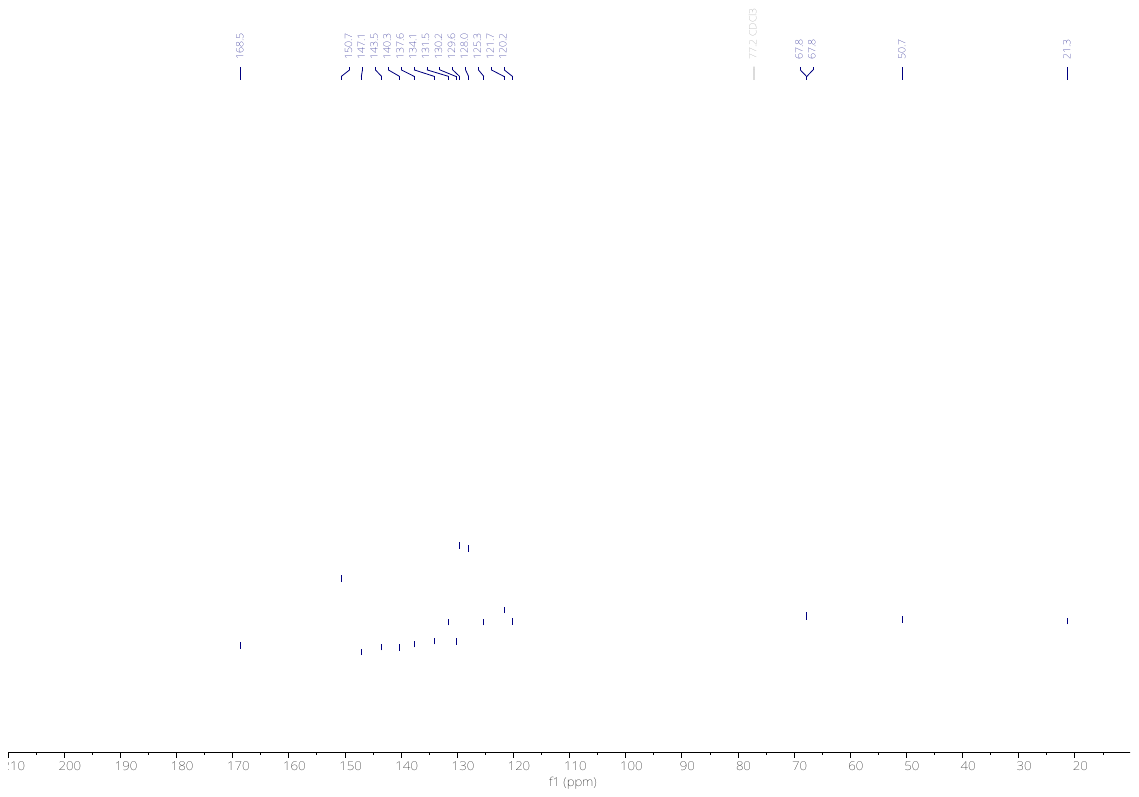

3704

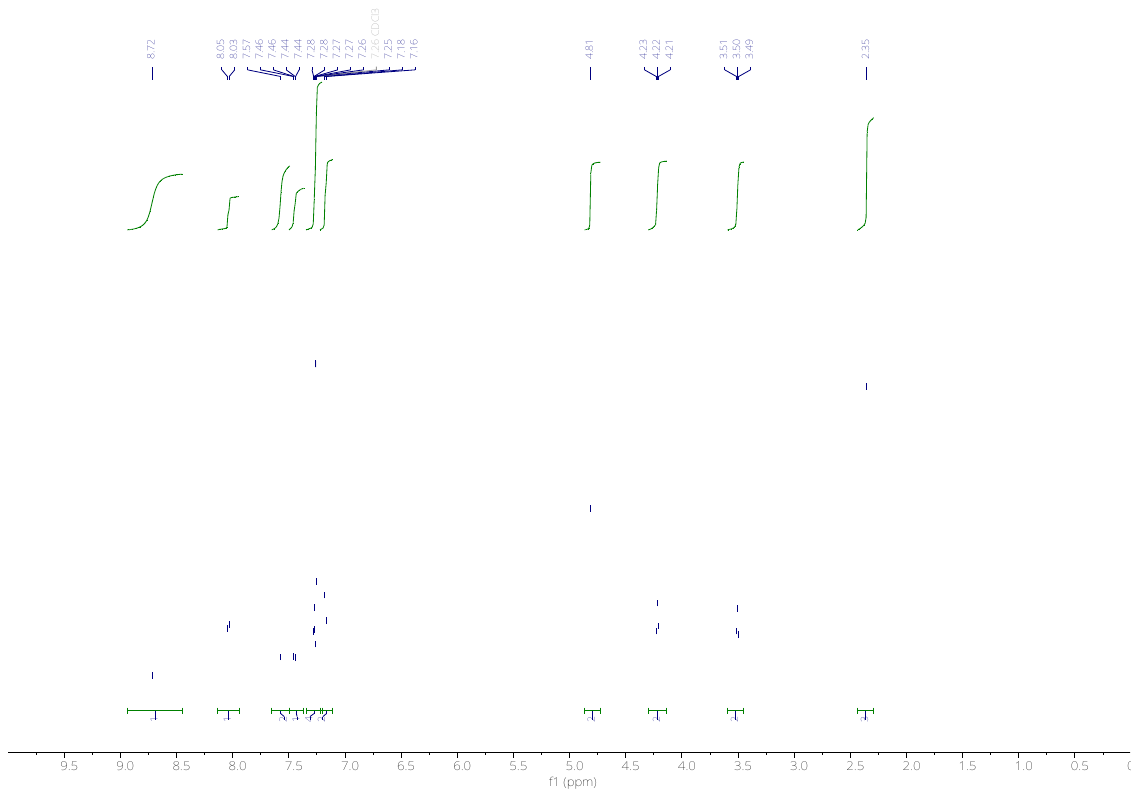

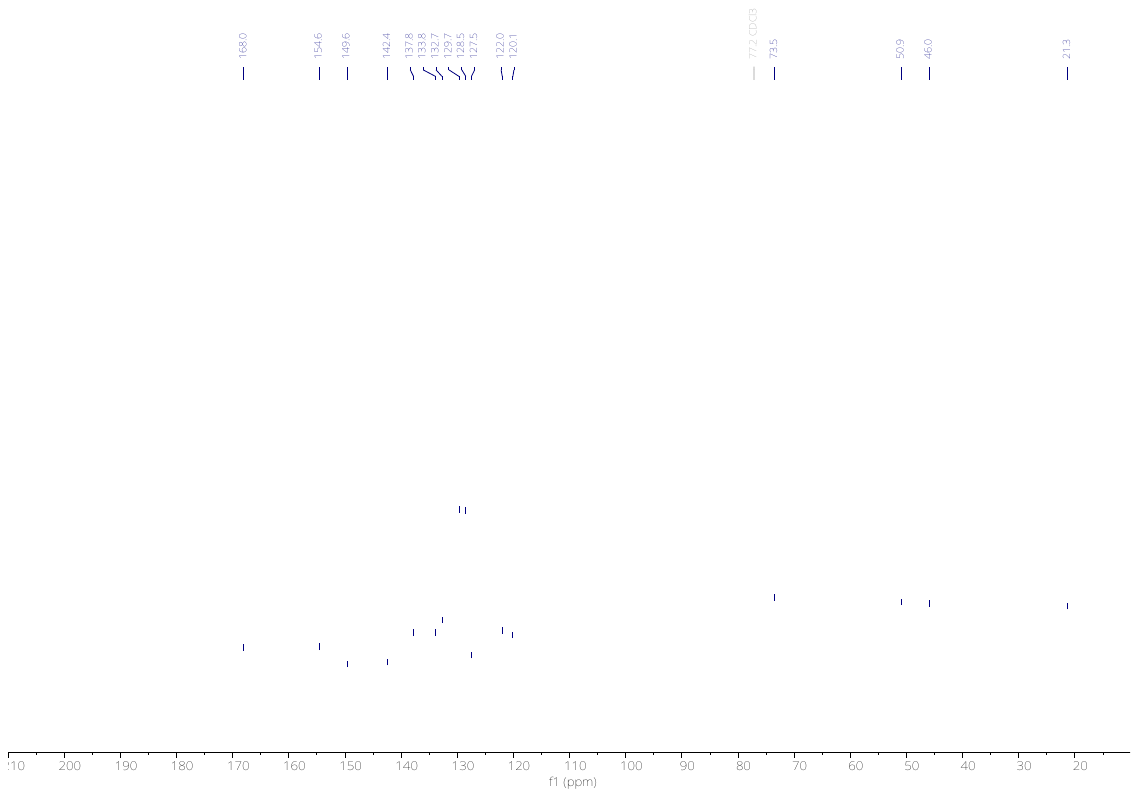

3707

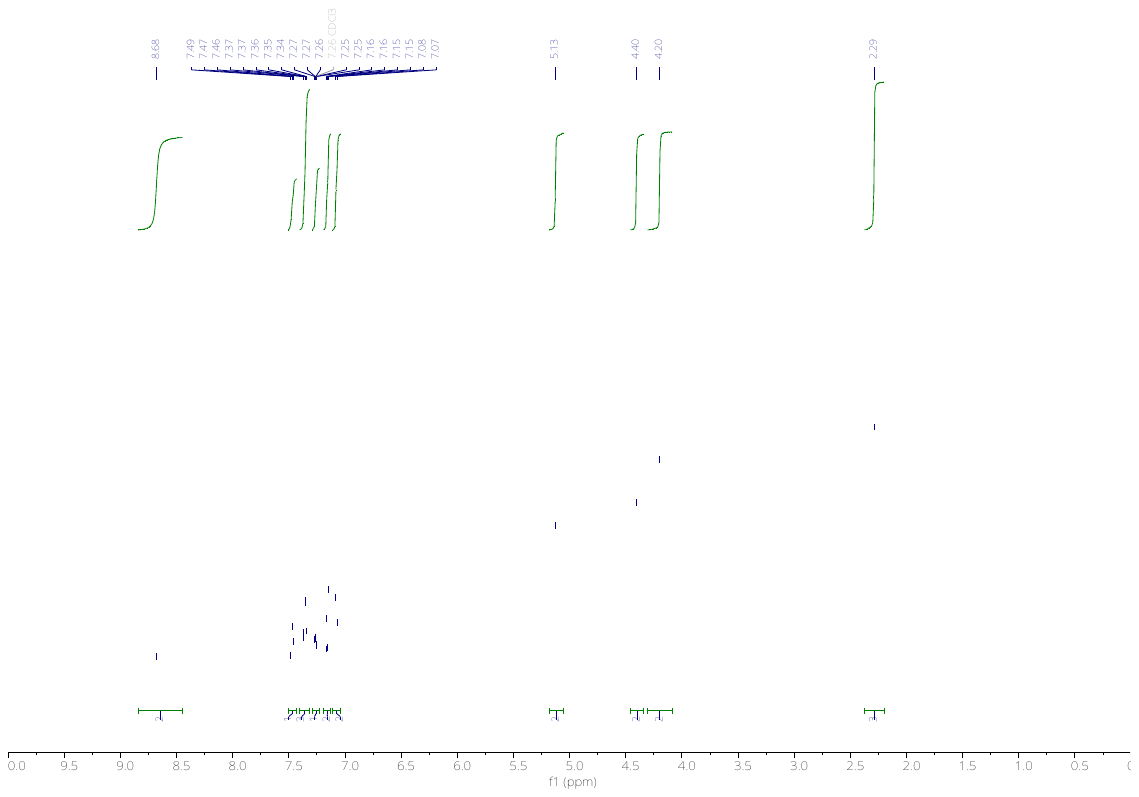

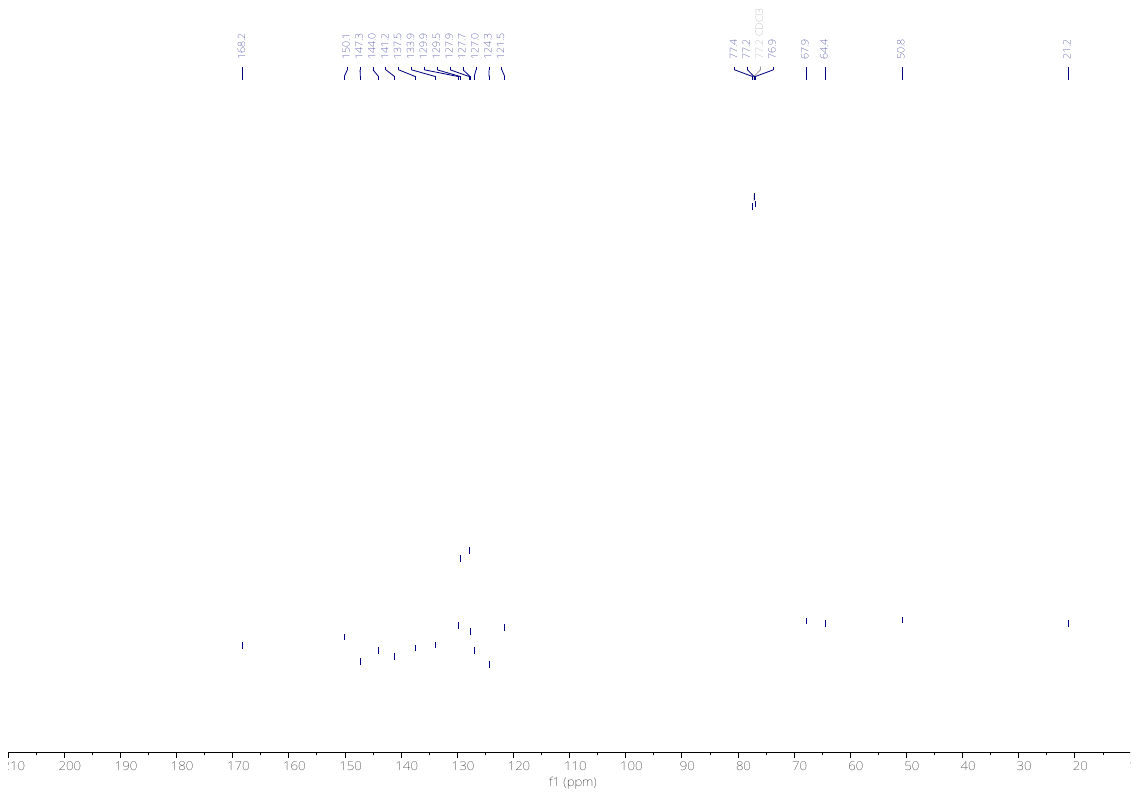

3710

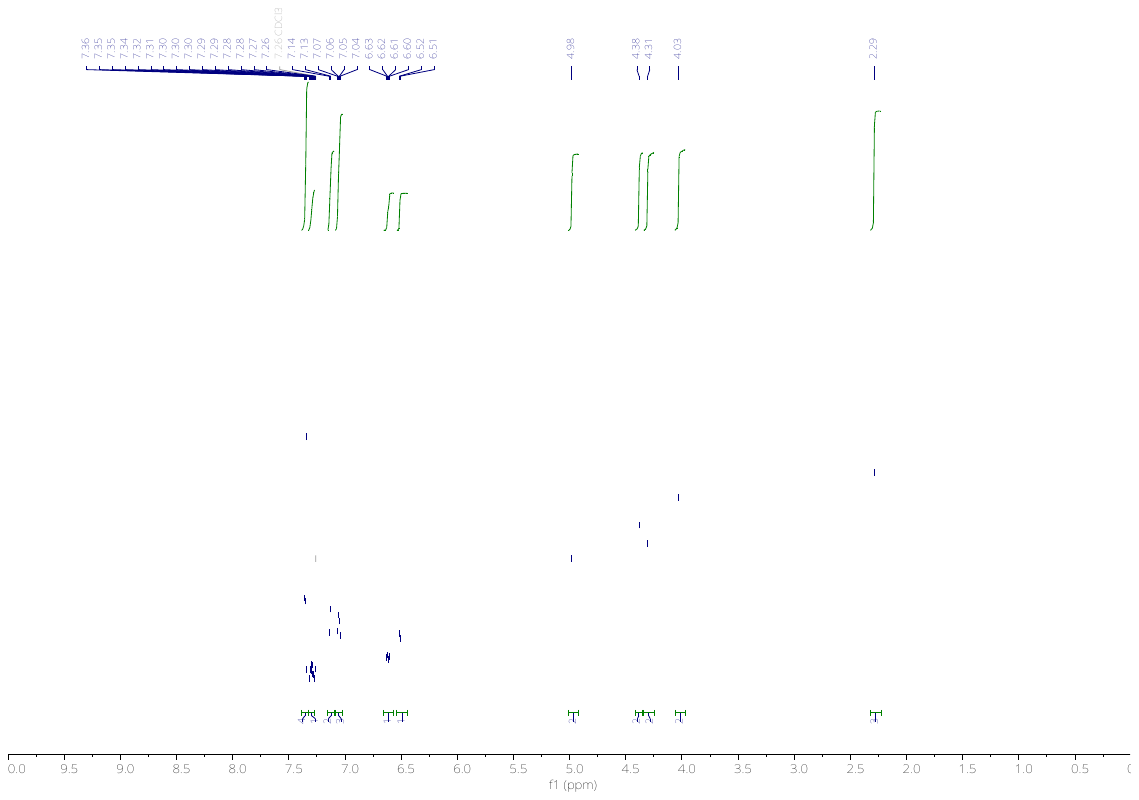

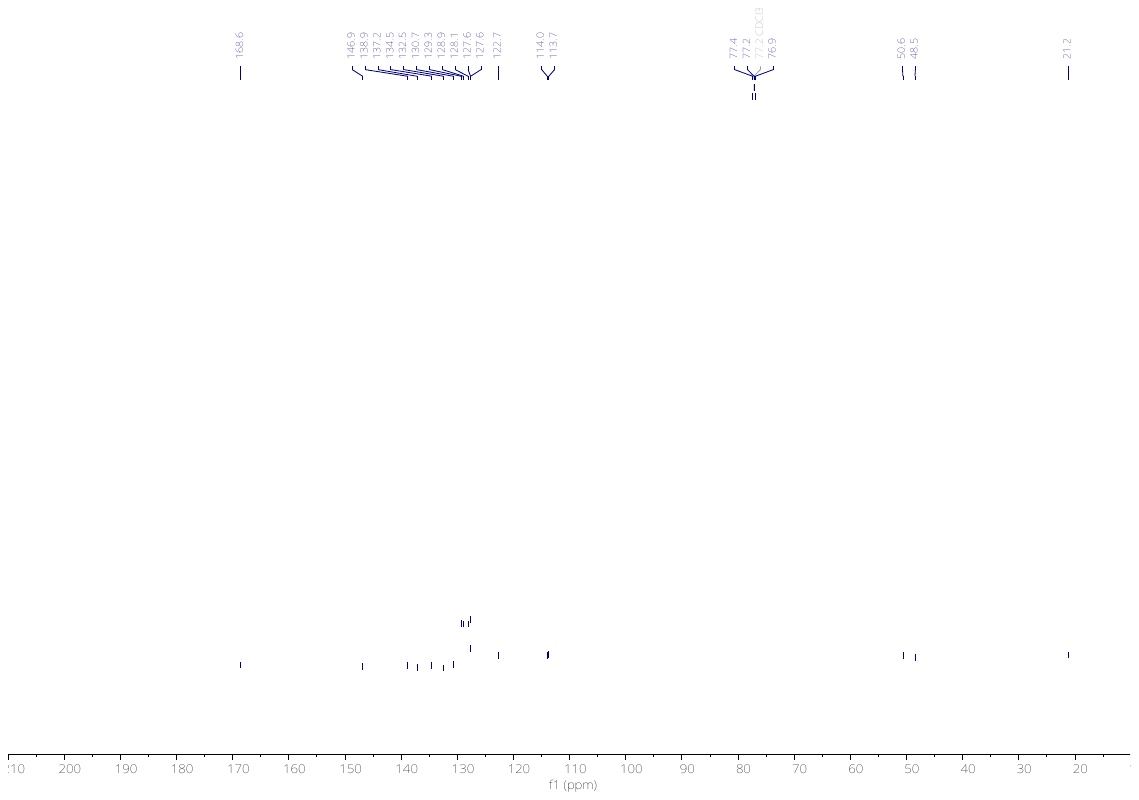

3824

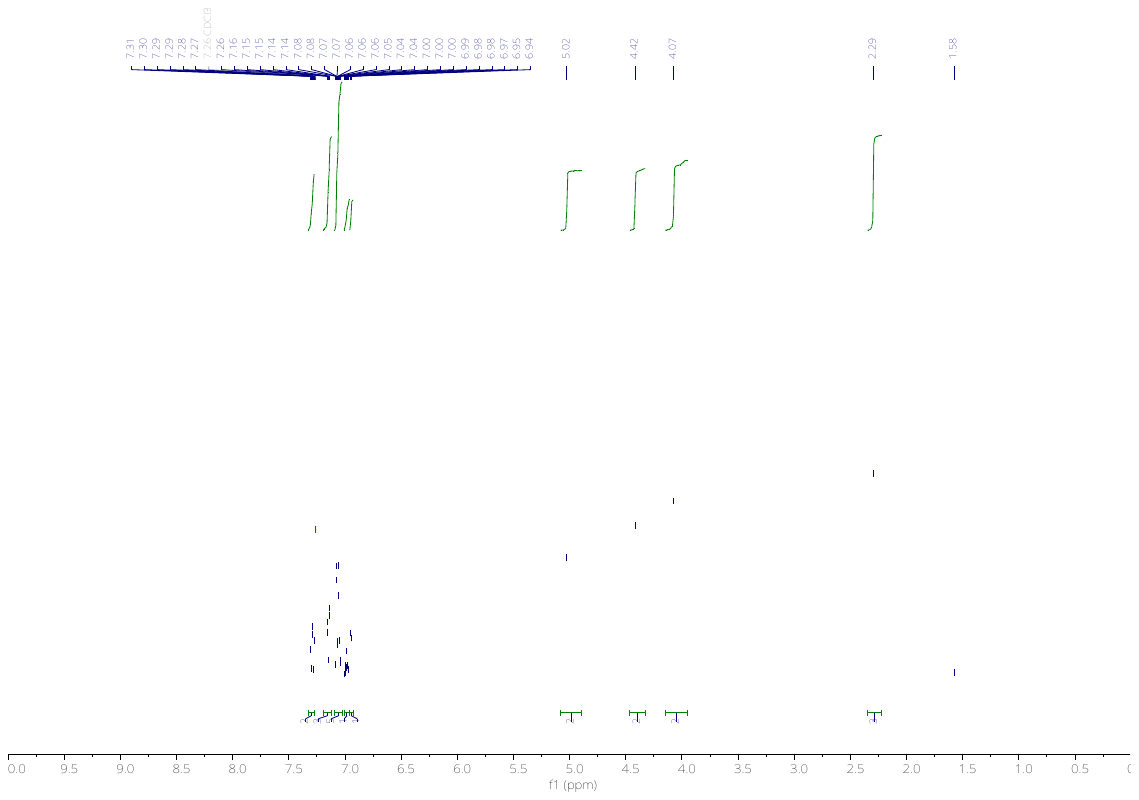

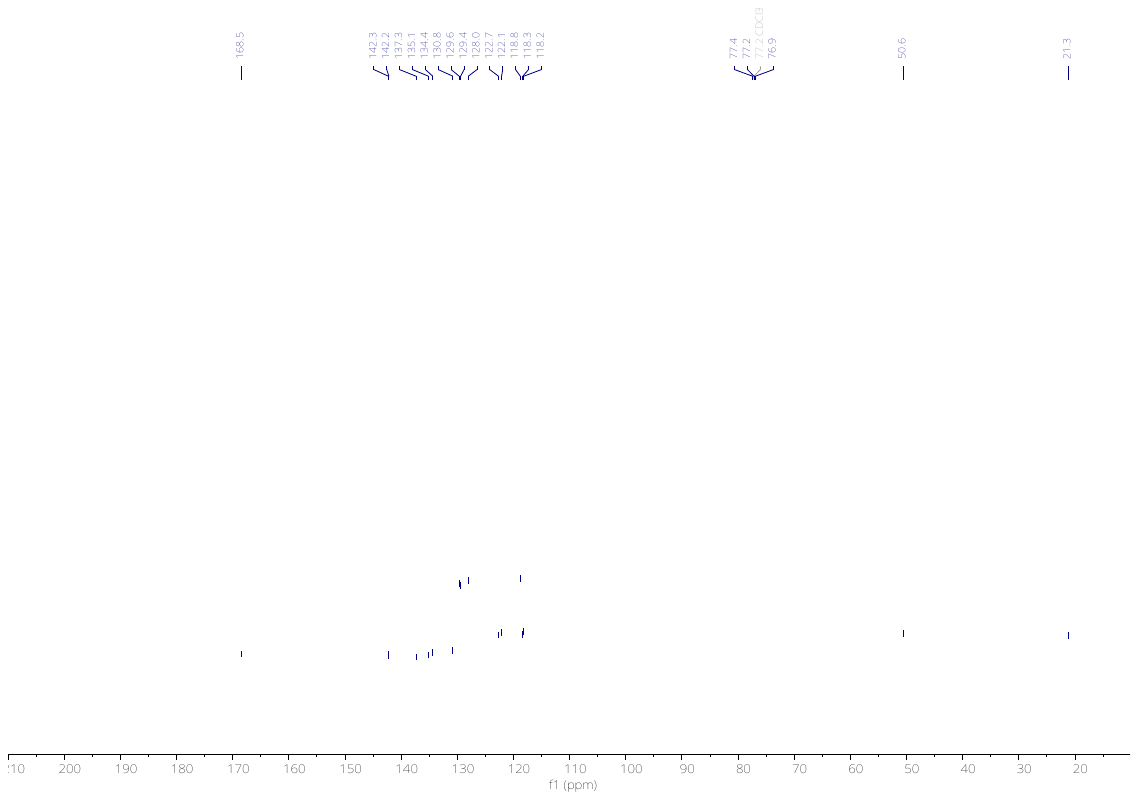

3825

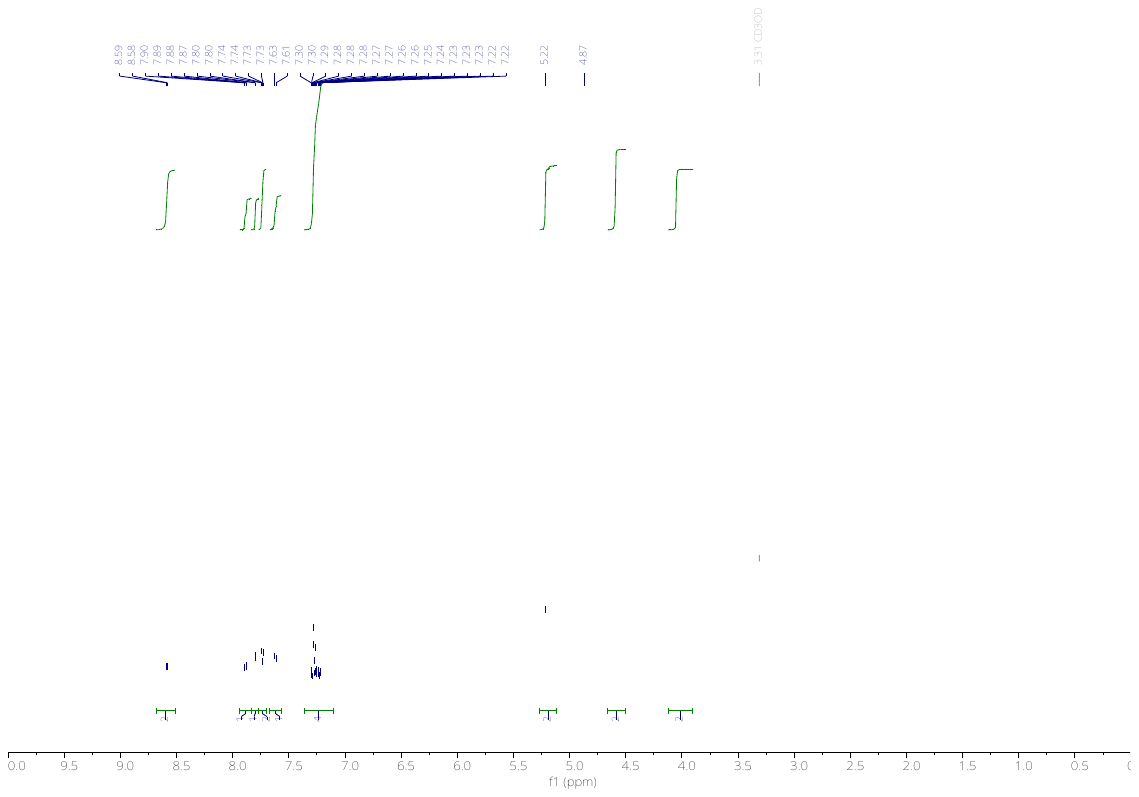

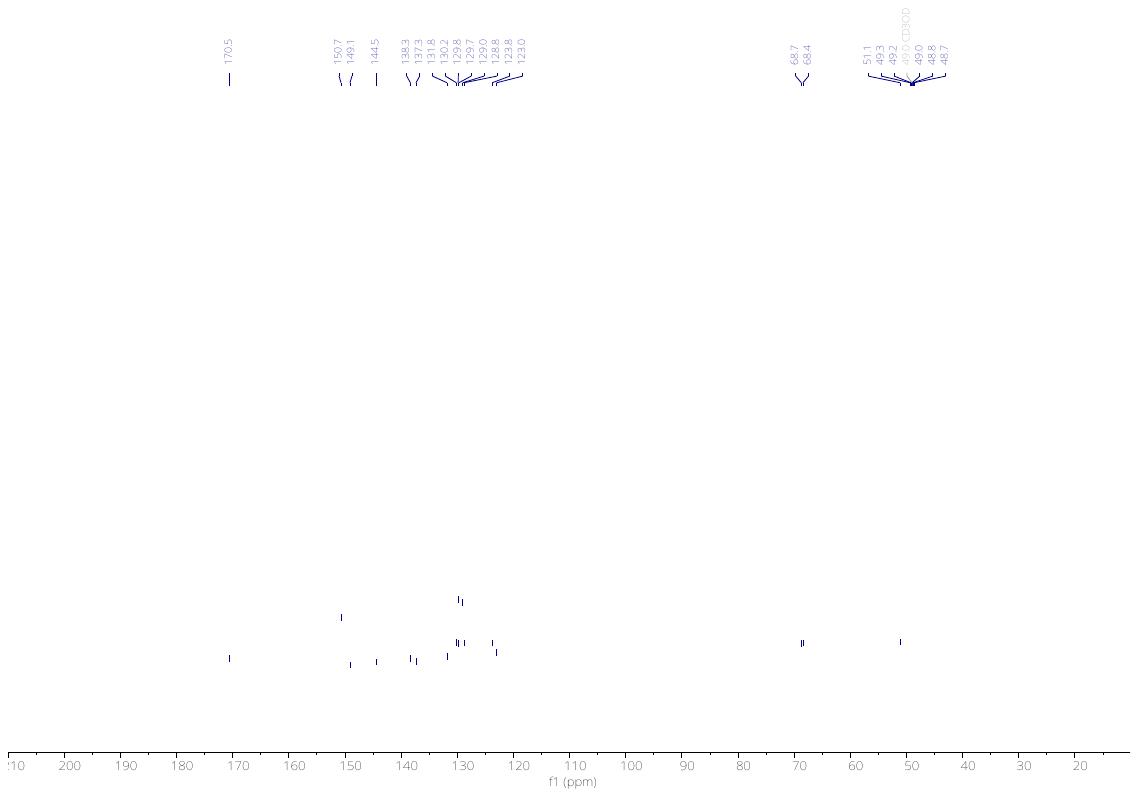

4115

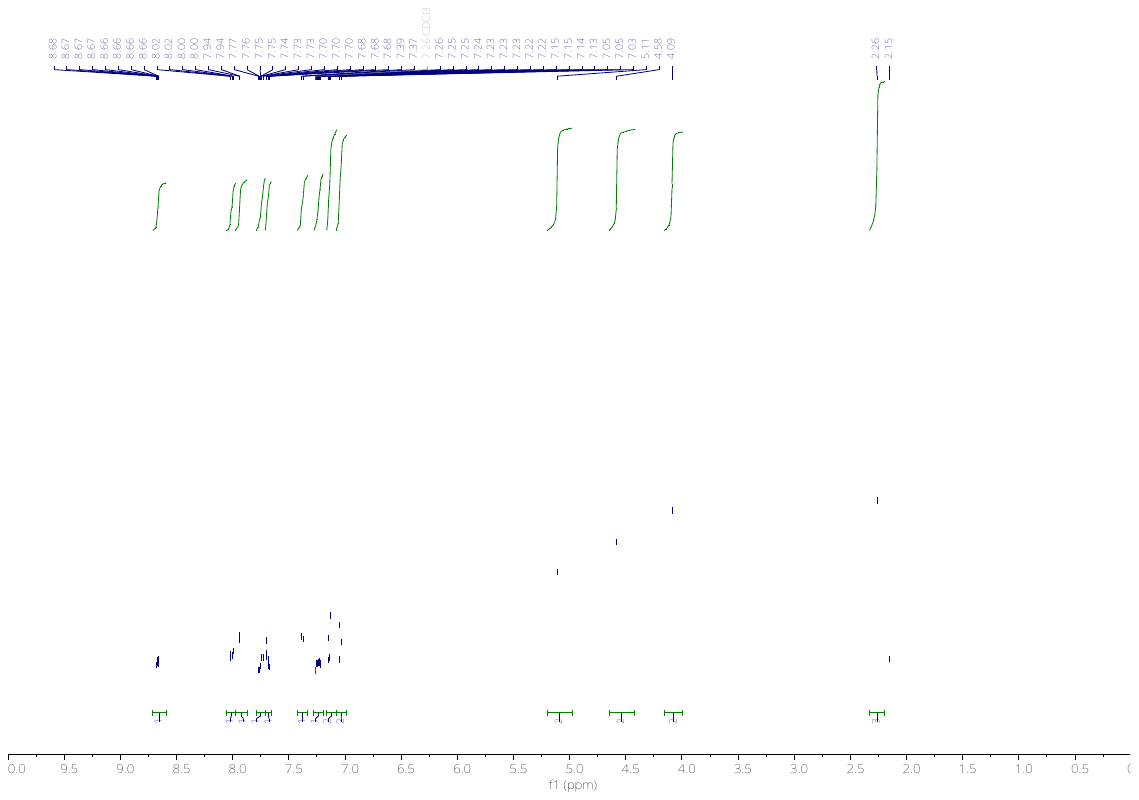

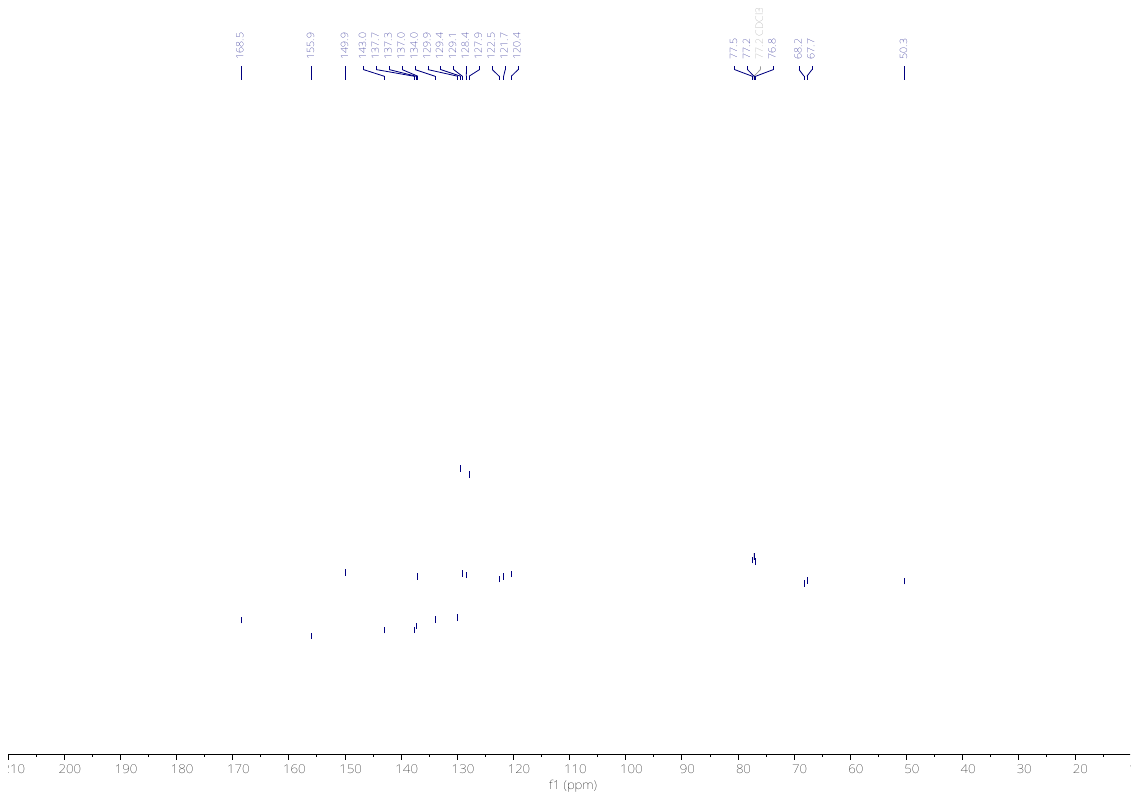

4116

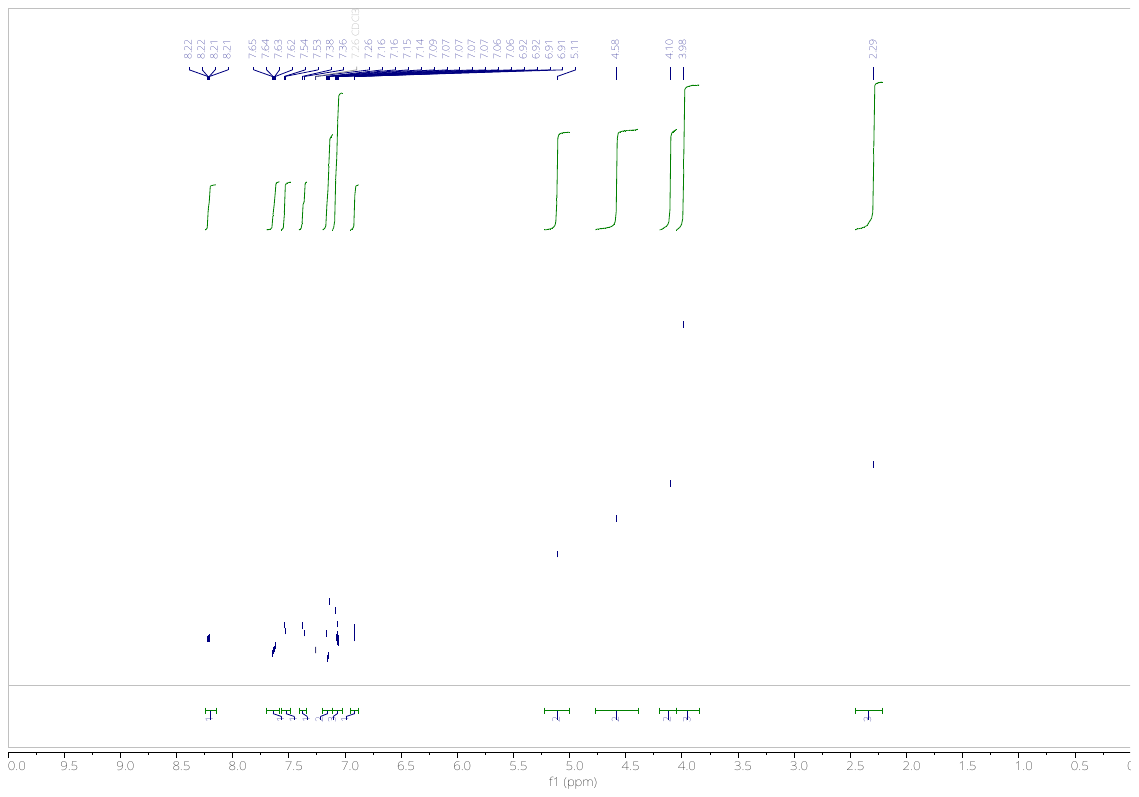

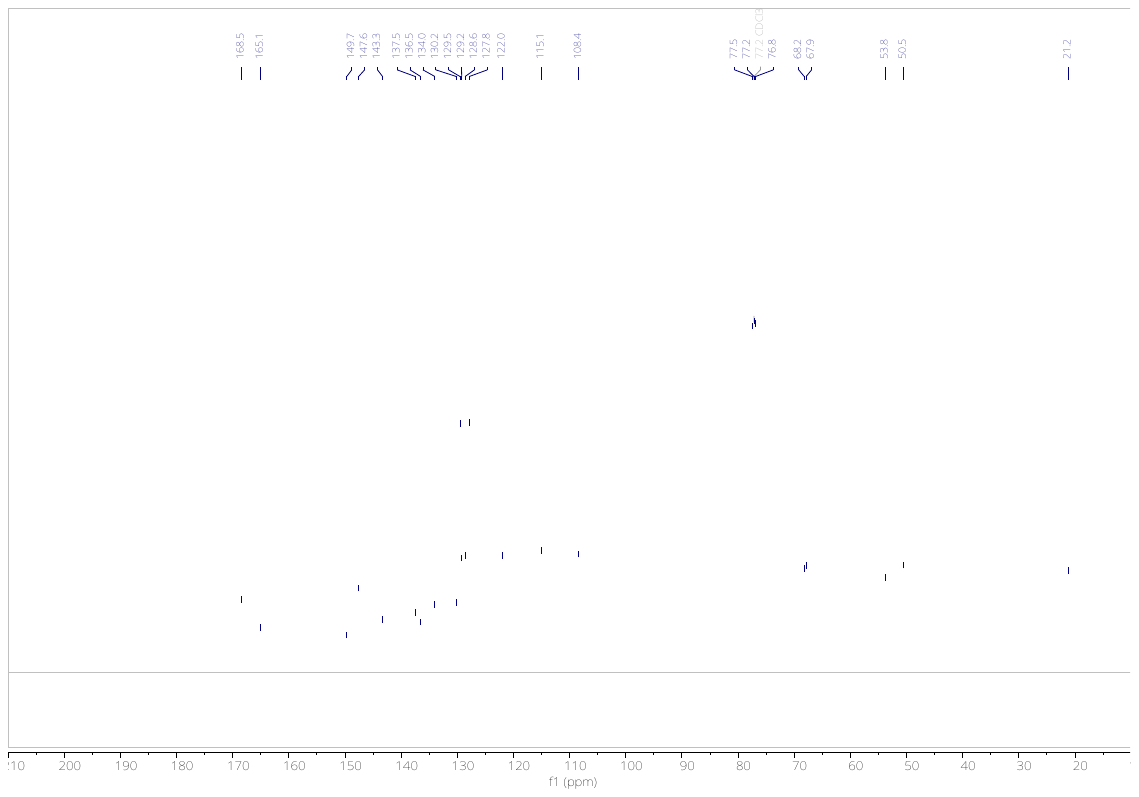

4117

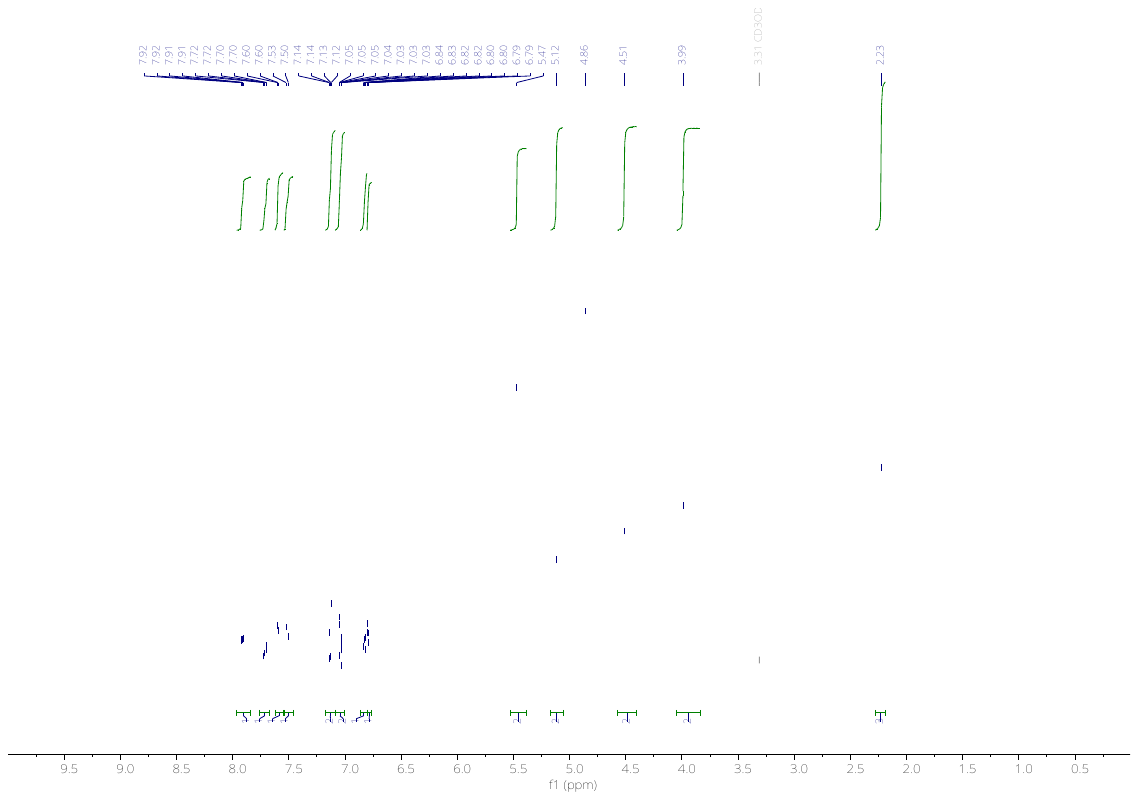

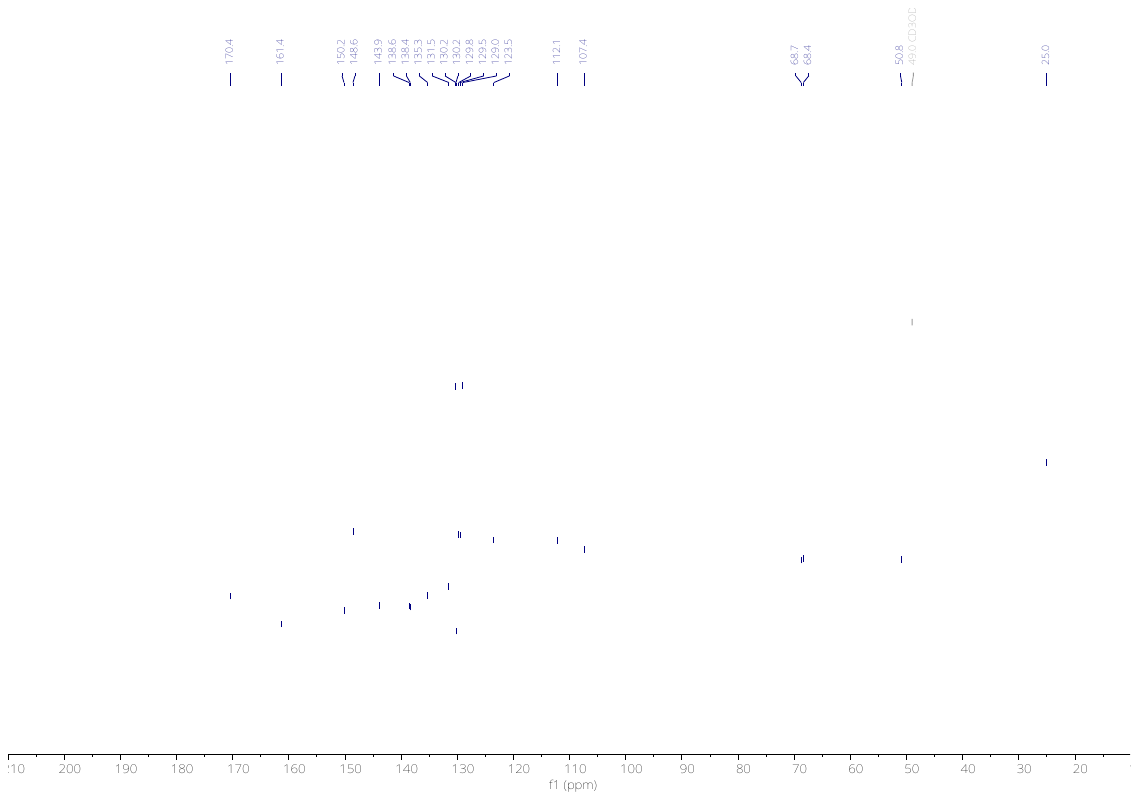

4118

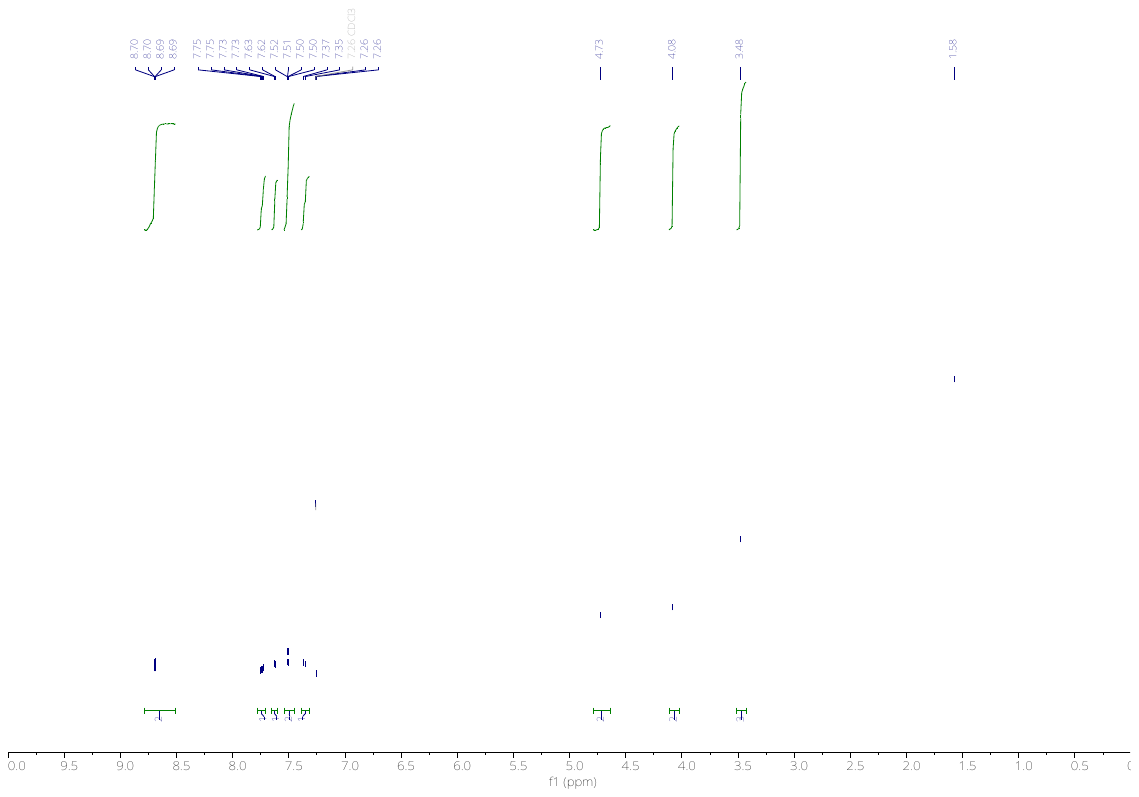

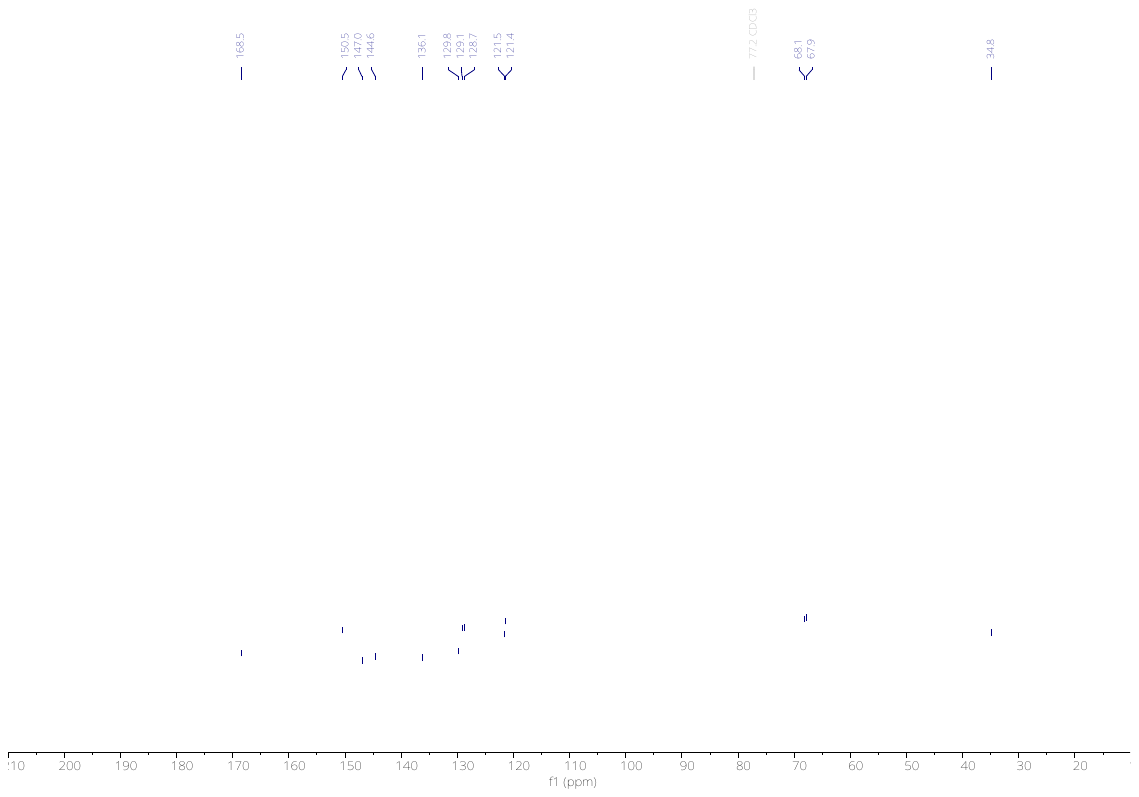

4120

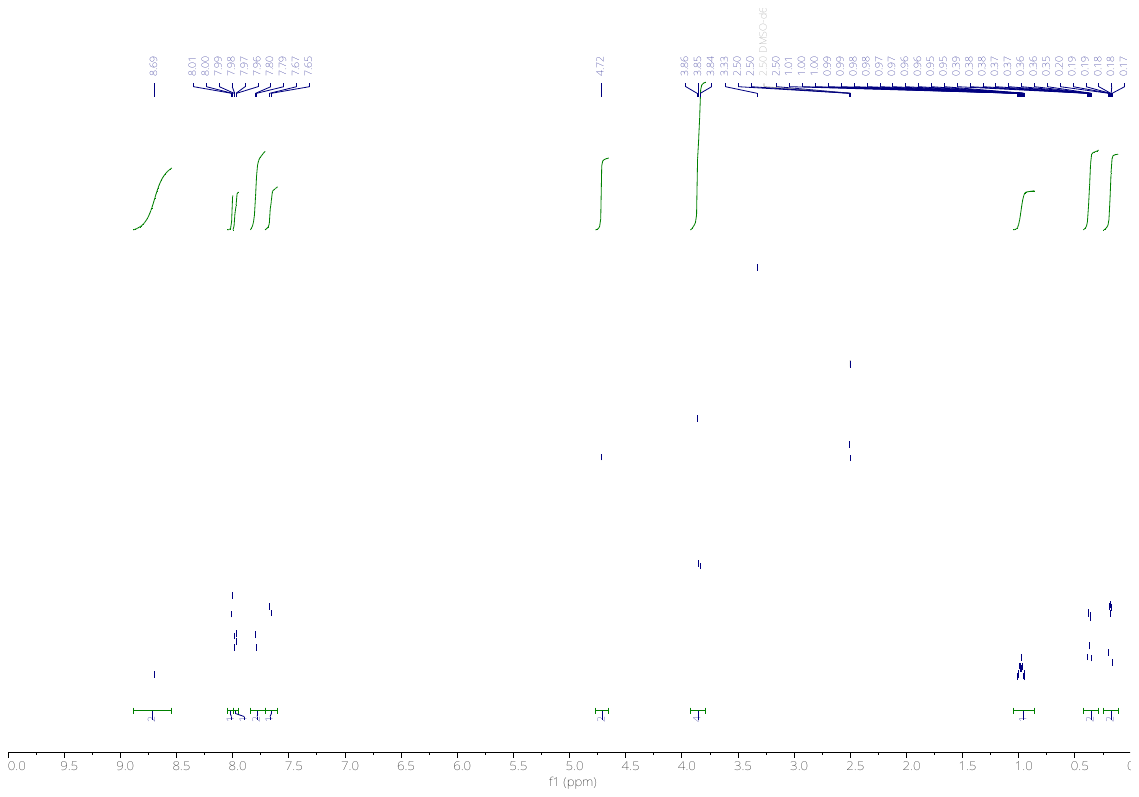

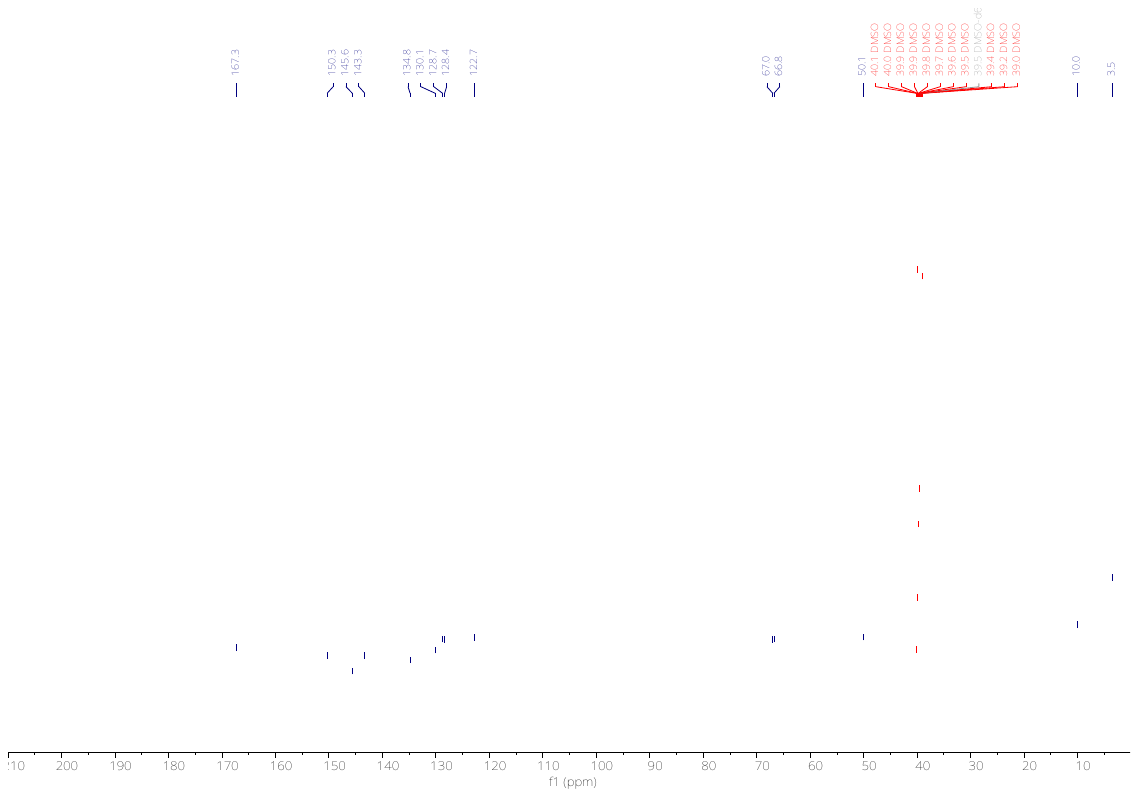

4122

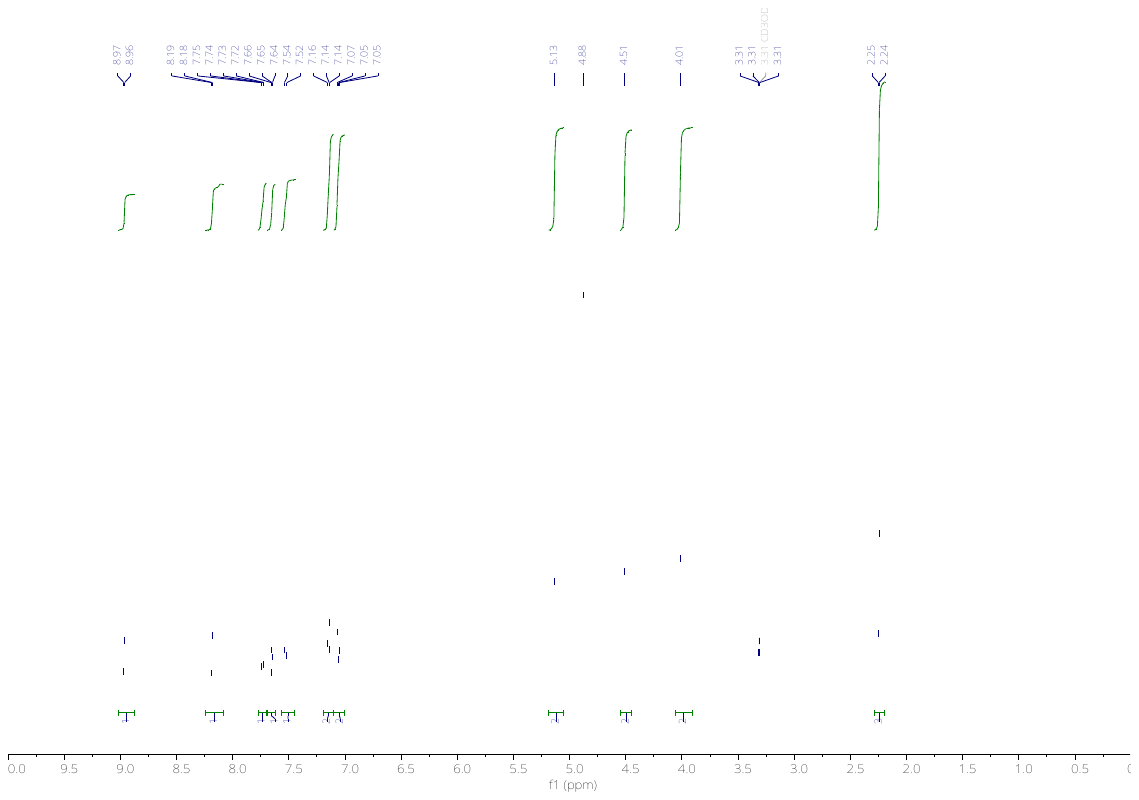

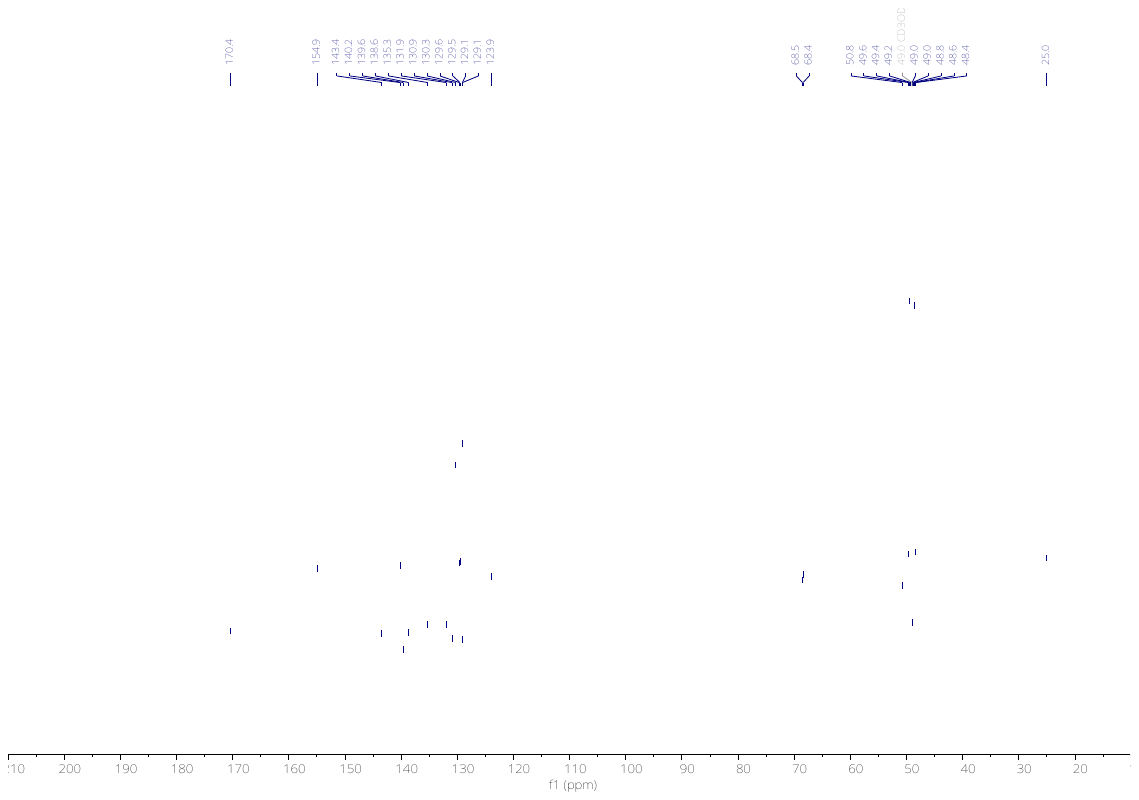

4123

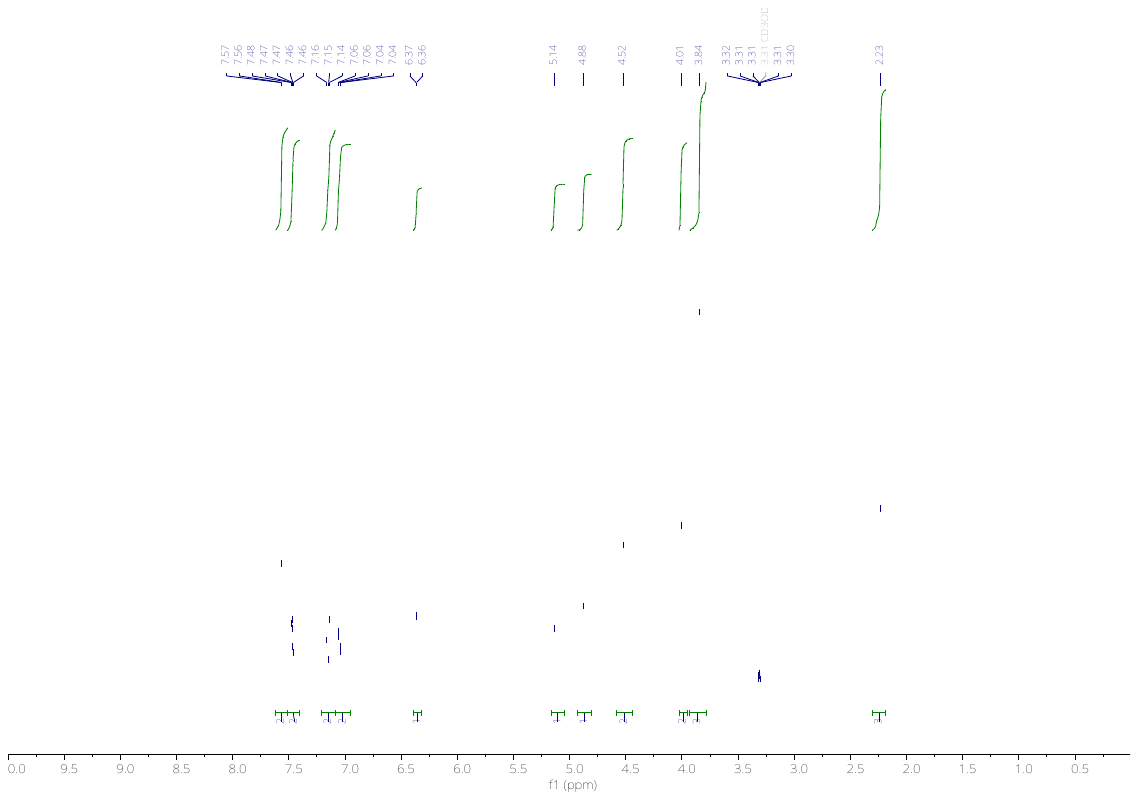

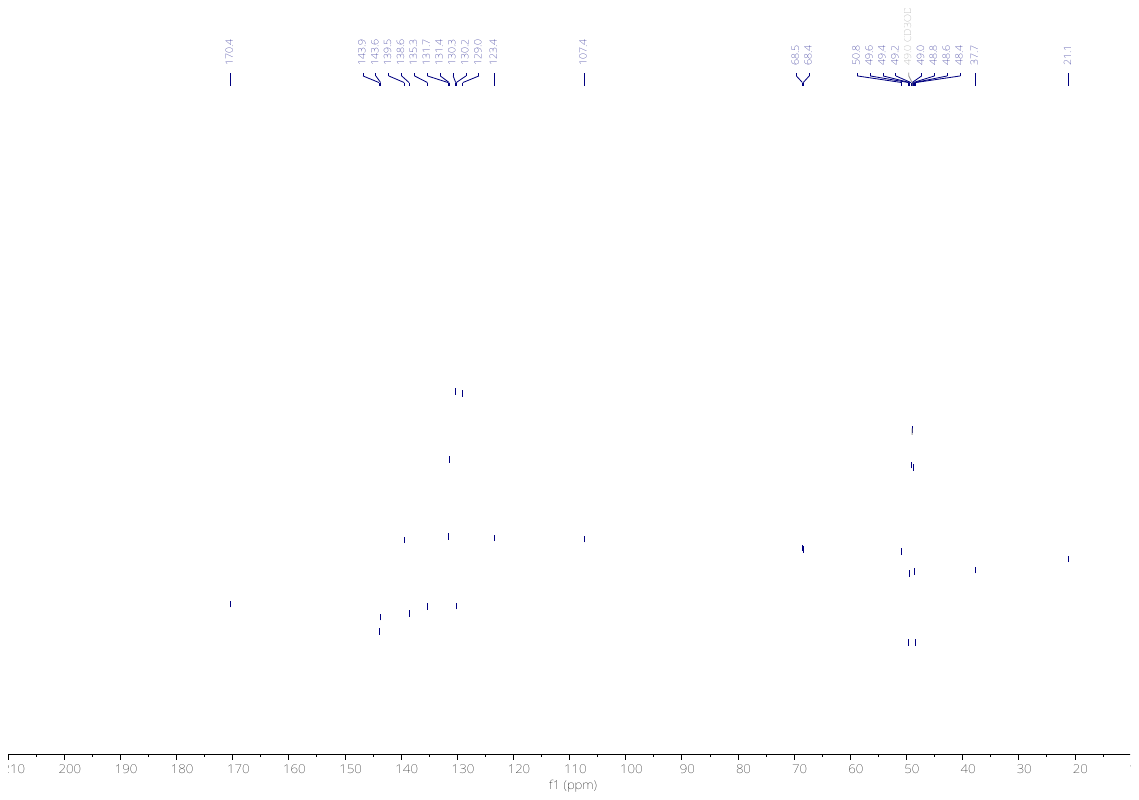

3699

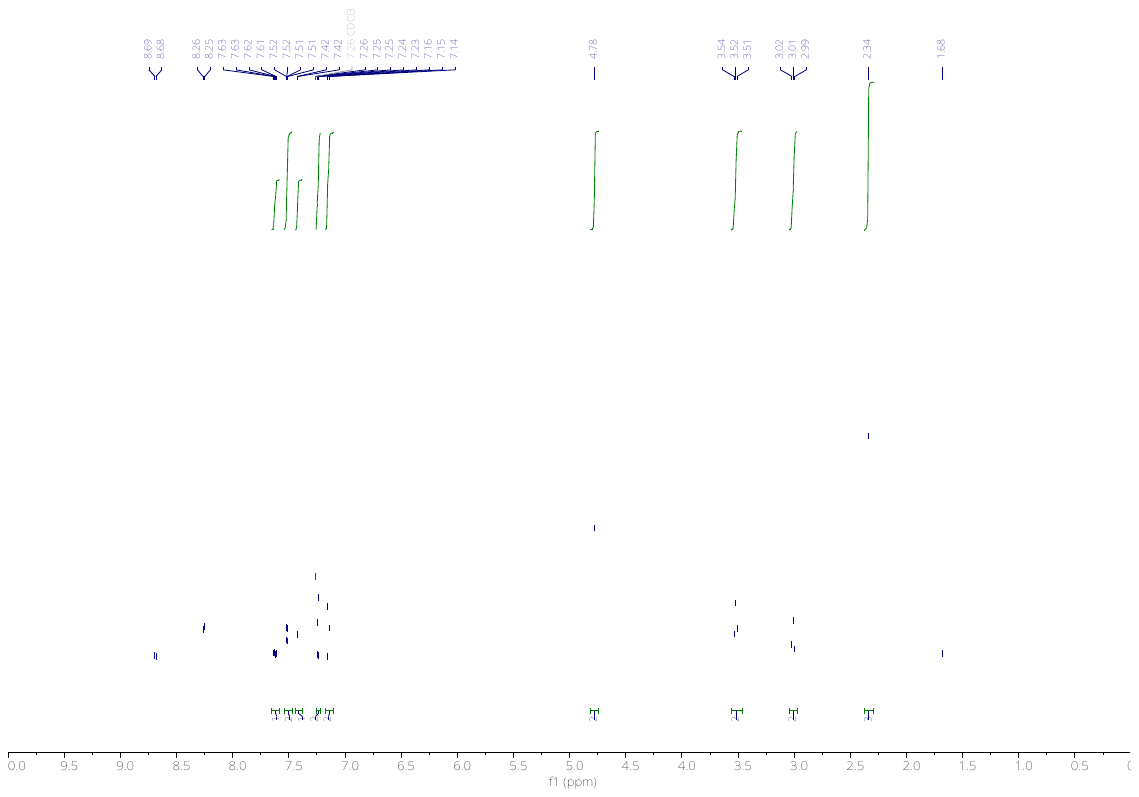

4236

4237

4238

4239

#### References

Coppola, G.M., Damon, R.E., Kukkola, P.J., Stanton, J.L., 2004. Amide derivatives and their use as inhibitors of 11-beta-hydroxysteroid dehydrogenase type 1. WO2004065351A1.

Partridge, F.A., Murphy, E.A., Willis, N.J., Bataille, C.J.R., Forman, R., Heyer-Chauhan, N., Marinič, B., Sowood, D.J.C., Wynne, G.M., Else, K.J., Russell, A.J., Sattelle, D.B., 2017. Dihydrobenz[*e*][1,4]oxazepin-2(3*H*)-ones, a new anthelmintic chemotype immobilising whipworm and reducing infectivity *in vivo*. PLoS Negl Trop Dis 11, e0005359. https://doi.org/10.1371/journal.pntd.0005359

Zablocki, J.A., Elzein, E., Li, X., Koltun, D.O., Parkhill, E.Q., Kobayashi, T., Martinez, R., Corkey, B., Jiang, H., Perry, T., Kalla, R., Notte, G.T., Saunders, O., Graupe, M., Lu, Y., Venkataramani, C., Guerrero, J., Perry, J., Osier, M., Strickley, R., Liu, G., Wang, W.-Q., Hu, L., Li, X.-J., El-Bizri, N., Hirakawa, R., Kahlig, K., Xie, C., Li, C.H., Dhalla, A.K., Rajamani, S., Mollova, N., Soohoo, D., Lepist, E.-I., Murray, B., Rhodes, G., Belardinelli, L., Desai, M.C., 2016. Discovery of Dihydrobenzoxazepinone (GS-6615) Late Sodium Current Inhibitor (Late Na i), a Phase II Agent with Demonstrated Preclinical Anti-Ischemic and Antiarrhythmic Properties. Journal of Medicinal Chemistry acs.jmedchem.6b00939. https://doi.org/10.1021/acs.jmedchem.6b00939
